# Human septins in cells organize as octamer-based filaments mediating actin-membrane anchoring

**DOI:** 10.1101/2022.02.23.481653

**Authors:** Carla Silva Martins, Cyntia Taveneau, Gerard Castro-Linares, Mikhail Baibakov, Nicolas Buzhinsky, Mar Eroles, Violeta Milanović, Francois Iv, Léa Bouillard, Alex Llewellyn, Maxime Gomes, Mayssa Belhabib, Mira Kuzmić, Pascal Verdier-Pinard, Stacey Lee, Ali Badache, Sanjay Kumar, Cristel Chandre, Sophie Brasselet, Felix Rico, Olivier Rossier, Gijsje H. Koenderink, Jerome Wenger, Stéphanie Cabantous, Manos Mavrakis

## Abstract

Septins are cytoskeletal proteins conserved from algae and protists to mammals. Septin knock-out animals have established that septins are essential for animal physiology, but their molecular function remains elusive. A unique feature of septins is their presence as heteromeric complexes that polymerize into filaments in solution and on lipid membranes. Although animal septins associate extensively with actin-based structures in cells, whether actin-decorating septins organize as filaments and if septin organization impacts septin function is not known. Customizing a tripartite split-GFP complementation assay for probing the presence and composition of septin filaments *in situ* in cells, we show that all septins decorating actin stress fibers are present as filaments whose integrity depends on octameric septin protomers. Atomic force microscopy nanoindentation measurements on cells confirmed that cell stiffness depends on the presence of octamer-containing septin filaments. Super-resolution structured illumination microscopy revealed septin fibers with widths compatible with their organization as paired septin filaments. Nanometer-resolved distance measurements and single-protein tracking further showed that actin-associated septin filaments are membrane-bound and largely immobilized. Finally, reconstitution assays on supported lipid bilayers showed that septin filaments mediate actin-membrane anchoring. We propose that septin organization as octamer-based filaments is essential for septin function in anchoring and stabilizing actin fibers at the plasma membrane.

## Introduction

Septins comprise a family of cytoskeletal proteins conserved from algae and protists to mammals (Cao et al., 2007; Momany et al., 2008; Nishihama et al., 2011; Pan et al., 2007). Septins were discovered in budding yeast already 50 years ago (Hartwell, 1971; Hartwell et al., 1970) as mutants that result in cytokinesis defects (Hartwell, 1971), and follow-up studies in animal model systems established that they are also required for animal cell division (Echard et al., 2004; Estey et al., 2010; Founounou et al., 2013; Kechad et al., 2012; Kinoshita et al., 1997; Neufeld and Rubin, 1994; Surka et al., 2002). However, septins are expressed in practically all human tissues, including non-dividing neurons (Karlsson et al., 2021). There is compelling evidence that septins play roles in a wide range of biological processes in non-dividing animal cells and tissues, including cell motility, sperm integrity, neuron development, tissue morphogenesis, and host-pathogen interactions (Fares et al., 1995; Finger et al., 2003; Gilden et al., 2012; Ihara et al., 2005; Kim et al., 2010; Kissel et al., 2005; Kuo et al., 2012; Mostowy et al., 2010; Mostowy et al., 2011; Nguyen et al., 2000; Shindo and Wallingford, 2014; Steels et al., 2007; Tada et al., 2007; Tooley et al., 2009; Xie et al., 2007). The embryonic lethality of mouse and *Drosophila* septin knock-outs (Adam et al., 2000; Fuchtbauer et al., 2011; Menon et al., 2014; Roseler et al., 2011) emphasizes their essential contribution to animal physiology and development, yet the precise molecular basis of their function in dividing and non-dividing cells remains elusive.

Biochemical isolation of native septins from budding yeast, *Drosophila* and mammalian cells and tissues revealed that septins exist as stable heteromeric complexes that can polymerize into filaments and higher-order filament assemblies (Field et al., 1996; Frazier et al., 1998; Hsu et al., 1998; Kim et al., 2011; Kinoshita et al., 2002; Sellin et al., 2011). The isolation of recombinant septin complexes helped establish that septin complexes are palindromes, with each septin in two copies and in a specific position within the complex, with each monomer interacting with its neighbors by alternating interfaces, named NC (from the N- and C-terminal domains) and G (from the GTP-binding domain) (Bertin et al., 2008; DeRose et al., 2020; Farkasovsky et al., 2005; Garcia et al., 2011; Huijbregts et al., 2009; Iv et al., 2021; John et al., 2007; Kinoshita et al., 2002; Kumagai et al., 2019; Mavrakis et al., 2014; Mendonca et al., 2019; Rosa et al., 2020; Sala et al., 2016; Sirajuddin et al., 2007; Soroor et al., 2021; Versele and Thorner, 2004). Human septins are classified in four homology groups, namely the SEPT2 group (SEPT1, 2, 4, and 5), SEPT6 group (SEPT6, 8, 10, 11, and 14), SEPT7 group (SEPT7), and SEPT3 group (SEPT3, 9, and 12) (Kinoshita, 2003). Cell-isolated human septins exist as stable hexamers and octamers (Kim et al., 2011; Sellin et al., 2011; Sellin et al., 2014), with hexamers composed of septins from the SEPT2, SEPT6, SEPT7 groups, and octamers containing additional septins from the SEPT3 group (Fig. 1A). Whereas SEPT3 and SEPT12 are enriched in the brain and the testis, respectively, SEPT9 is expressed in practically all human cell lines and tissues (Karlsson et al., 2021; Uhlen et al., 2015).

**Figure 1.**
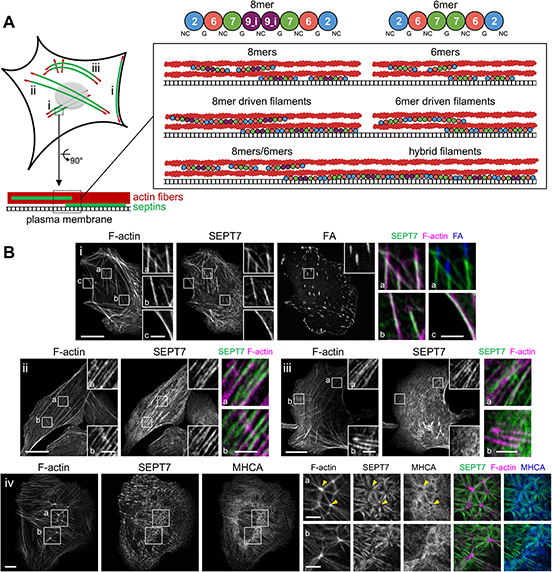
SEPT7 distribution on different types of stress fibers in U2OS cells. **(A)** The scheme on the left depicts septin-decorated stress fibers (SFs) in a mammalian cell. Septins (green) decorate different types of actin SFs (red), notably peripheral and ventral SFs (i), perinuclear actin caps (ii) and transverse arcs (iii), shown in the respective panels (i-iii) in (B). The cartoons on the right show human septin octamers and hexamers associating with stress fibers either as single protomers (top), as octamer and hexamer-driven filaments (middle), or as hybrid filaments from octamer and hexamer co-polymerization (bottom). Septins can associate exclusively with SFs or also with the plasma membrane. **(B)** Representative confocal micrographs of SEPT7 immunostainings showing examples of SEPT7 localizing (i) to ventral (a,b) and peripheral (c) SFs and excluded from focal adhesions (FA) (a), (ii) to perinuclear actin caps (a,b), (iii) to transverse arcs (a) and excluded from dorsal SFs (b), and (iv) to ventral actin nodes (a,b). Cells are co-stained for F-actin (phalloidin), the FA protein paxillin, and myosin heavy chain (MHCA). Yellow arrowheads in (iv) point to two actin nodes. Scale bars in large fields of views, 10 μm. Scale bars in insets, 2 μm (i-iii) and 5 μm (iv).

The feature of septins that led to the hypothesis that they make up the fourth cytoskeleton element is the capacity of recombinant and cell-isolated septins to self-assemble into filaments (Mostowy and Cossart, 2012; Valadares et al., 2017). The most convincing evidence that septins form filaments *in vivo*, and that cell viability depends on their ability to assemble into filaments, comes from electron microscopy and functional data in budding yeast (Bertin et al., 2012; Byers and Goetsch, 1976; McMurray et al., 2011; Ong et al., 2014; Rodal et al., 2005). The conservation of septins and the ability of recombinant and cell-purified mammalian septin hexamers and octamers (hereafter referred to as protomers) to self-assemble into filaments in solution and on lipid membranes (DeRose et al., 2020; Iv et al., 2021; Leonardo et al., 2021; Soroor et al., 2021; Szuba et al., 2021) has led to the assumption that human septins also organize as filaments in cells, but formal evidence that this is the case is scarce. Immunogold electron microscopy in mammalian cells has shown gold-decorated septins in close apposition to cortical actin filaments and to the plasma membrane organizing in linear arrays (Hagiwara et al., 2011; Kinoshita et al., 1997). Single septin protomers along actin filaments or the membrane would, however, result in a similar pattern, as it would be the case for any actin filament- or membrane-binding protein. It is reasonable to assume that septin rings and fiber-looking segments that form in the cytoplasm of mammalian cells upon actin depolymerization correspond to septin filaments or bundles thereof (Joo et al., 2007; Kim et al., 2011; Kinoshita et al., 2002; Schmidt and Nichols, 2004). However, one cannot exclude that septins on stress fibers are not filamentous, and that single septin protomers on stress fibers spontaneously form filaments in the absence of stress fibers. Furthermore, it was not shown that the observed septin fibers upon actin disassembly originate from direct end-to-end septin complex polymerization. Whether all septins in cells function as filaments, and how hexamers and octamers contribute to septin filament formation and function are not known.

The fact that mammalian septins associate with membranes as well as with the actin and microtubule cytoskeleton has made it difficult to dissect septin function, and has at the same time naturally led to the speculation that septins mediate cytoskeleton-membrane cross-talk (Dogterom and Koenderink, 2019). There is no doubt that septins can associate both with actin and microtubules in cells (Bowen et al., 2011; Nagata et al., 2004; Nagata et al., 2003; Spiliotis et al., 2008; Spiliotis et al., 2005; Surka et al., 2002; Verdier-Pinard et al., 2017). The microtubule-binding domain on septins was identified recently (Kuzmic et al., 2022), but actin-binding domains have not yet been identified making it unclear if actin-septin binding involves direct interactions, or if such binding occurs through myosin-II (Joo et al., 2007; Mostowy et al., 2010) or/and Borg proteins (Calvo et al., 2015; Farrugia et al., 2020; Joberty et al., 2001; Liu et al., 2014; Salameh et al., 2021). Similarly, although recombinant and cell-purified mammalian septins bind lipid membranes in the absence of other physiological partners (Bridges et al., 2016; Dolat and Spiliotis, 2016; Szuba et al., 2021; Tanaka-Takiguchi et al., 2009; Yamada et al., 2016), whether there is direct septin-membrane binding in cells has not been formally shown; the identification of the membrane-binding site of septins is a matter of debate (Cavini et al., 2021). Septin-membrane association in dividing cells necessitates the presence of Anillin, which itself binds the ingressing plasma membrane and recruits septins to it (Field et al., 2005; Hickson and O’Farrell, 2008; Liu et al., 2012; Renshaw et al., 2014). Given that Anillin is nuclear in interphase cells (Field and Alberts, 1995; Oegema et al., 2000), it is not known if septin-decorated actin fibers and membranes in non-dividing cells reflect membrane-bound septins.

To elucidate the interplay between human septin organization and function in non-dividing cells, notably septin-actin-membrane interactions in cells, we used actin stress fibers in U2OS cells as a model system. Septins in mammalian cells have been reported to decorate stress fibers in multiple studies over the last 25 years (Calvo et al., 2015; Connolly et al., 2011; Dolat et al., 2014; Joo et al., 2007; Kim et al., 2011; Kinoshita et al., 2002; Kinoshita et al., 1997; Liu et al., 2014; Salameh et al., 2021; Schmidt and Nichols, 2004; Surka et al., 2002; Verdier-Pinard et al., 2017; Xie et al., 1999; Zhang et al., 1999). Subsets of stress fibers are lost upon septin disruption or septin relocalization to microtubules (Calvo et al., 2015; Kinoshita et al., 2002; Kuzmic et al., 2022; Salameh et al., 2021; Schmidt and Nichols, 2004; Targa et al., 2019) suggesting an essential, yet still unclear, role of septins in actin fiber formation or/and maintenance. To test if septins organize as filaments in cells and determine septin filament composition, we combined a tripartite split-GFP complementation assay with mutants disrupting specific septin-septin interfaces in order to selectively perturb hexamers or octamers, or abolish polymerization altogether. Atomic force microscopy nanoindentation measurements on cells were used to assess the specific contribution of hexamers vs octamers to cell stiffness. We employed super-resolution structured illumination microscopy to decipher the higher-order assembly of septin filaments. Moreover, to determine whether septin filaments are membrane-bound and if they have the capacity to bridge membrane-actin interactions, we combined nanometer-resolved distance measurements and single protein tracking in cells with cell-free reconstitution assays using supported lipid bilayers. Our findings demonstrate that all actin-associated septins in cells organize as paired membrane-bound filaments whose integrity and function depend on octamers.

## Results

### Septins associate with contractile stress fibers

Septins in mammalian cells have been shown to localize to stress fibers (SFs) in multiple studies, but whether septins associate preferentially with specific types of SFs and if septin organization differs among SFs is not known. To answer these questions, we started by examining how septins distribute with respect to the different types of SFs in U2OS cells, notably peripheral, dorsal and ventral SFs, transverse arcs and the perinuclear actin cap (Fig. 1A). Given that SFs are classified based on their subcellular localization and their anchoring at one or both ends by focal adhesions (FAs) (Tojkander et al., 2012), we acquired confocal z-stack images of cells co-stained for septins, actin filaments and the FA proteins, paxillin or vinculin. We chose to examine the distribution of three septins expressed in U2OS cells, namely SEPT2, SEPT7 and SEPT9 (Fig. 1B; Figs. S1, S2). SEPT2 and SEPT7 are common to both hexamers and octamers, whereas SEPT9 is specific to octamers (Fig. 1A). Both septin immunostainings and imaging of septin-GFP fusions showed that the distribution of all three septins with respect to SFs was identical. They all decorated myosin-II containing contractile SFs (Fig. 1Bi-iii; Fig. S1Ai-ii, iv-v; Fig. S1Bi-vi), but not the non-contractile dorsal ones (Fig. 1Biii,b; Fig. S1Aiii; Fig. S1Biv,a). Although septins extensively decorated contractile SFs throughout their length, they were systematically excluded from FAs (Fig. 1Bi,a; Fig. S1Ai,c; Fig. S1Bi,c; Fig.S2D). In addition to the localization of septins to peripheral and ventral SFs, transverse arcs and perinuclear actin caps, we also found septins associated with two types of actin nodes: geodesic actin nodes on the ventral plasma membrane and actin nodes in transverse arcs (Fig. 1Biv; Fig. S2A,C). Actin nodes were enriched in F-actin and *α*-actinin, while actin filaments interconnecting actin nodes were decorated by septins and myosin-II in an aster-like pattern.

U2OS cells express two SEPT9 isoforms, SEPT9_i1 and SEPT9_i3 (Kuzmic et al., 2022), both of which are detected by our SEPT9 antibodies. The presence of SEPT9 showed that septin octamers are present on SFs, but does not exclude the possibility that septin hexamers are also present. Furthermore, the diffraction-limited optical resolution of our setup does not allow us to distinguish single septin protomers from septin filaments. Septin decoration of SFs may therefore reflect the presence of either single protomers (hexamers and/or octamers) or of filaments driven by hexamer or octamer polymerization, or hexamer and octamer co-polymerization (Fig. 1A).

### Septins organize as filaments on contractile SFs

Both hexamers and octamers have an exposed SEPT2 NC interface at their termini (Fig. 1A) (Iv et al., 2021; Mendonca et al., 2019; Soroor et al., 2021), thus detecting direct SEPT2-SEPT2 interactions would provide a molecular readout of end-to-end septin polymerization of hexamers or/and octamers driving the formation of septin filaments. To determine whether septins are present as filaments, we designed a tripartite split-GFP complementation assay for probing SEPT2-SEPT2 interactions *in situ* in living cells (Fig. 2A,B). This protein-protein interaction assay involves the fusion of the proteins of interest to the two last beta-strands of GFP, β10 and β11: in the presence of specific protein-protein interactions in cells expressing GFP1-9 (GFP strands β1-β9), the GFP barrel is reconstituted leading to fluorescence (Cabantous et al., 2013) (see methods and Fig. S3A-C for the design of the assay). The implementation of the tripartite split-GFP complementation assay in budding yeast for probing septin positioning was shown to measure intimate short-range physical contacts and accurately reflected the spatial relationship among septin subunits within the octamer (Finnigan et al., 2016). To probe SEPT2-SEPT2 interactions, we generated β10- and β11-strand fusions with SEPT2 that we co-expressed using an inducible bidirectional vector in U2OS cells constitutively expressing GFP1-9 (Fig. 2A; Fig. S3D). The fluorescence from the reconstitution of the GFP barrel will occur only in the case of specific SEPT2-SEPT2 interactions and will thus report the subcellular localization of such interactions as a molecular signature of the presence of septin filaments.

**Figure 2.**
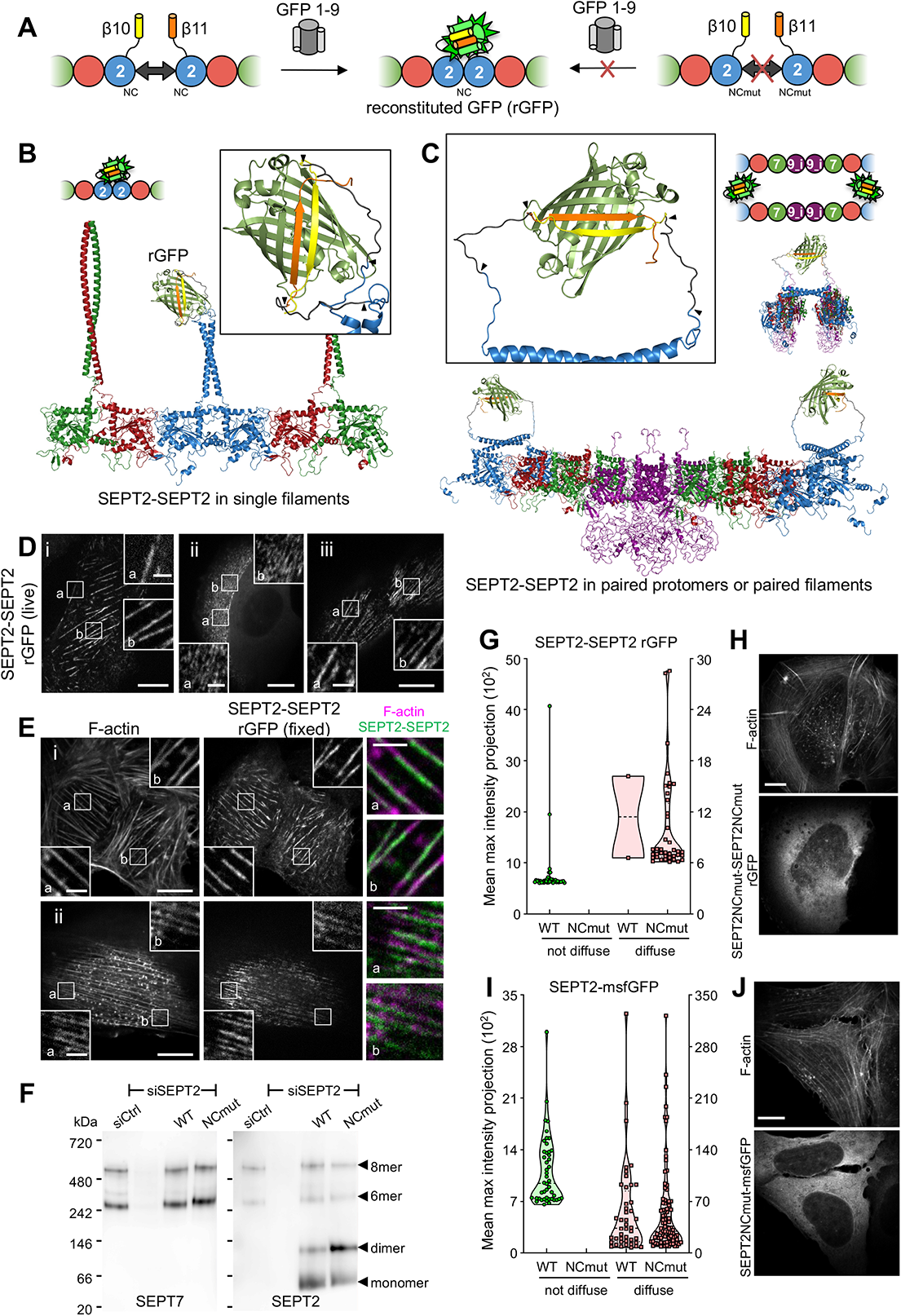
All septins on SFs organize as filaments. **(A)** Schematic representation of the tripartite split-GFP complementation assay for probing SEPT2-SEPT2 interactions. The transparency of the SEPT7 subunits is used to suggest that the polymerizing protomers can be hexamers or/and octamers. **(B-C)** Structure models of reconstituted GFP (rGFP) via direct SEPT2-SEPT2 interactions of two polymerizing septin protomers within a filament (B) or from SEPT2 in two apposed protomers (C) (octamers are shown for simplicity). Only the end-to-end interacting halves of the protomers are shown in (B) for simplicity; protomers can be hexamers or/and octamers. SEPT6 and SEPT7 coiled-coils are not shown in (C) for simplicity. The transparency of the terminal SEPT2 subunits in (C) is used to suggest that the paired protomers could be found within a filament. β10 and β11 strands are shown in yellow and orange, respectively. Linker sequences between septins and the β-strands, delimited by arrowheads, are shown in dark grey. The colors of septin subunits in the structure models correspond to the ones in the color-coded sphere representation of hexamers and octamers. The second half of the octamer is not shown in the rotated filament pair in (C) for the sake of simplicity. **(D-E)** Representative confocal micrographs of SEPT2-SEPT2 reconstituted GFP (rGFP) distribution in live cells (D) and in fixed cells (E) co-stained for F-actin (phalloidin). Examples show rGFP in live cells localizing (i) to peripheral (a) and ventral (b) SFs, (ii) to transverse arcs (a,b), and (iii) to actin caps (a,b). Examples in fixed cells show rGFP localizing (i) to ventral SFs (a,b) and (ii) to actin caps (a,b). Scale bars in large fields of views, 10 μm. Scale bars in insets, 2 μm. **(F)** Western blot following native PAGE of U2OS cell lysates probed with anti-SEPT7 (left) and anti-SEPT2 (right) antibodies upon treatment with siRNAs targeting LacZ (siCtrl), SEPT2 (siSEPT2), and targeting SEPT2 while expressing wild-type SEPT2-msfGFP (WT) or SEPT2NCmut-msfGFP (NCmut). Molecular weight markers are shown on the left. The overexpression of the msfGFP fusions leads to SEPT2 monomers and dimers in addition to hexamers and octamers (arrowheads). **(G)** Violin plots depicting the distribution of diffuse cytosolic (red datapoints) vs. non-diffuse (green datapoints) phenotypes from reconstituted GFP (rGFP) in GFP1-9 cells co-expressing wild-type SEPT2-β10 and -β11 or SEPT2NCmut-β10 and -β11. Data points are from a total of 40 cells each for wild-type and mutant SEPT2 distributed among the two phenotypes. **(H)** Representative example of a GFP1-9 cell co-expressing SEPT2NCmut-β10 and -β11 showing a diffuse cytosolic phenotype. Scale bar, 10 μm. **(I)** Violin plots depicting the distribution of diffuse cytosolic (red datapoints) vs. non-diffuse (green datapoints) phenotypes in cells expressing wild-type SEPT2-msfGFP or SEPT2NCmut-msfGFP. Data points are from a total of 90 cells each for wild-type and mutant SEPT2 distributed among the two phenotypes. **(J)** Representative example of a cell expressing SEPT2NCmut-msfGFP showing a diffuse cytosolic phenotype. Scale bar, 10 μm.

To minimize the risk of not detecting SEPT2-SEPT2 interactions due to endogenous untagged SEPT2 and given that the expression levels of SEPT2-β10/β11 fusions were kept low to minimize overexpression artifacts (Fig. S3E), we consistently knocked down endogenous SEPT2 in all subsequent experiments (Fig. S3F). Spinning disk confocal imaging in live cells and in fixed cells co-stained for actin revealed the presence of the reconstituted GFP (rGFP) on peripheral and ventral SFs, transverse arcs and perinuclear actin caps (Fig. 2D,E), with the rGFP distribution closely resembling endogenous SEPT2 immunostainings and SEPT2-GFP distribution (Fig. S1A). *In vitro* reconstitution studies have shown that fly and mammalian septins organize both as single and paired filaments (Szuba et al., 2021), prompting us to explore the origin of SEPT2-SEPT2 rGFP on SFs. To this end, we generated septin protomer structure models, and examined GFP complementation both from direct SEPT2-SEPT2 interactions within a hexamer (Fig. 2B), and from SEPT2 facing another SEPT2 in apposed hexamers or octamers of a paired septin filament (Fig. 2C). Examination of the distances and the flexibility of the SEPT2 C-termini and the linkers in such models showed that GFP reconstitution could originate either from direct SEPT2-SEPT2 interactions within one hexamer (Fig. 2B) or from SEPT2 facing another SEPT2 in apposed hexamers or octamers in a paired septin filament (Fig. 2C). Importantly, these models highlighted that paired protomers would lead to GFP reconstitution whether the protomers polymerize or not.

To test if SEPT2-SEPT2 rGFP on SFs originates from direct SEPT2-SEPT2 interactions, we designed a double point SEPT2 NC interface mutant (SEPT2 F20D, V27D, hereafter SEPT2NCmut) to prevent end-to-end association and thereby abolish polymerization (Fig. S4A) (Kuzmic et al., 2022; Sirajuddin et al., 2007). Reconstitution assays using purified recombinant hexamers and octamers bearing these mutations confirmed that this mutant abolishes septin polymerization, although it is still able to bind actin filaments *in vitro* (Fig. S4B-E). Native PAGE in cell lysates expressing SEPT2 NC interface mutants further confirmed that these mutants do not compromise protomer integrity: the expression of either wild-type SEPT2 or SEPT2NCmut in SEPT2 knockdown cells rescue equally well the hexamer and octamer distribution in control cells (Fig. 2F). Strikingly enough, using this mutant in the context of the split SEPT2-SEPT2 assay completely abolished SF localization as indicated by purely diffuse cytosolic fluorescence (Fig. 2G,H) (hereafter referred to as “diffuse cytosolic”). Given that wild-type SEPT2-SEPT2 rGFP was occasionally found as diffuse cytosolic, we quantified the distribution of diffuse cytosolic and non-diffuse phenotypes in cells expressing wild-type SEPT2-vs SEPT2NCmut-β10/β11 fusions (see methods). While 95% of wild-type SEPT2-SEPT2 rGFP localized to SFs, 100% of SEPT2NCmut-SEPT2NCmut rGFP was diffuse cytosolic (Fig. 2G,H). This result showed that direct end-to-end septin polymerization through an intact SEPT2-SEPT2 NC interface is required for septin localization to SFs and thus that septins on contractile SFs organize as single or paired septin filaments. We attribute the fact that the split assay with the NC mutant still produced fluorescence to the plasticity of septins which are able to use both NC and G interfaces when either one is compromised (Kim et al., 2012). We speculate that SEPT2NCmut forms G-homodimers, in line with earlier observations from other groups (Kim et al., 2012), thus enabling GFP complementation.

### Single septin protomers do not associate with SFs

The presence of septin filaments does not exclude that single septin protomers are also present on SFs. To test if single septin protomers associate with SFs, we examined the cellular distribution of the SEPT2 NC interface mutant fused to full-length GFP. Cells expressing this mutant exhibited a diffuse cytosolic localization, demonstrating that this mutant does not bind SFs (Fig. 2I,J). Given that wild-type SEPT2-GFP fusions also showed diffuse cytosolic localization in addition to SF localization (Fig. S1A,vi) and to confirm that the diffuse cytosolic phenotype was due to the incapacity of the NC mutant to bind SFs, we quantified the distribution of diffuse cytosolic and non-diffuse phenotypes in cells expressing wild-type SEPT2-GFP vs SEPT2NCmut-GFP fusions. While SEPT2-GFP was diffuse cytosolic in ∼50% of cells, 100% of the cells expressing SEPT2NCmut showed this phenotype (Fig. 2I,J). This result showed that single septin protomers in cells do not associate with SFs, meaning that all septins decorating SFs are filamentous.

### SF-associated septin filaments contain predominantly octamers

Our results showed that all septins decorating SFs are filamentous but did not inform us on the composition of septin filaments as SEPT2 is common to both hexamers and octamers (Fig. 1A). Recombinant hexamers and SEPT9-containing octamers have the capacity to co-polymerize *in vitro* (our own results in Fig. S5A and (Soroor et al., 2021)), but whether this is the case in cells is not known. The presence of SEPT9 on SFs (Fig. S1B) and native PAGE of cell lysates showing that all SEPT9 is incorporated in octamers (Fig. S5B) suggest that septin filaments contain octamers. To explicitly visualize the presence of octamers on SFs and to test if hexamers are also present there, we customized the tripartite complementation assay for probing specifically SEPT7-SEPT9 and SEPT9-SEPT9 as molecular signatures of octamers and SEPT7-SEPT7 as a molecular signature for hexamers. Given that both SEPT9_i1 and SEPT9_i3 isoforms localize to SFs, we probed the presence of octamers containing both these SEPT9 isoforms on SFs by detecting both SEPT7-SEPT9_i1 and SEPT9_i1-SEPT9_i1 and SEPT7-SEPT9_i3 and SEPT9_i3-SEPT9_i3 interactions (Fig. 3A,B).

**Figure 3.**
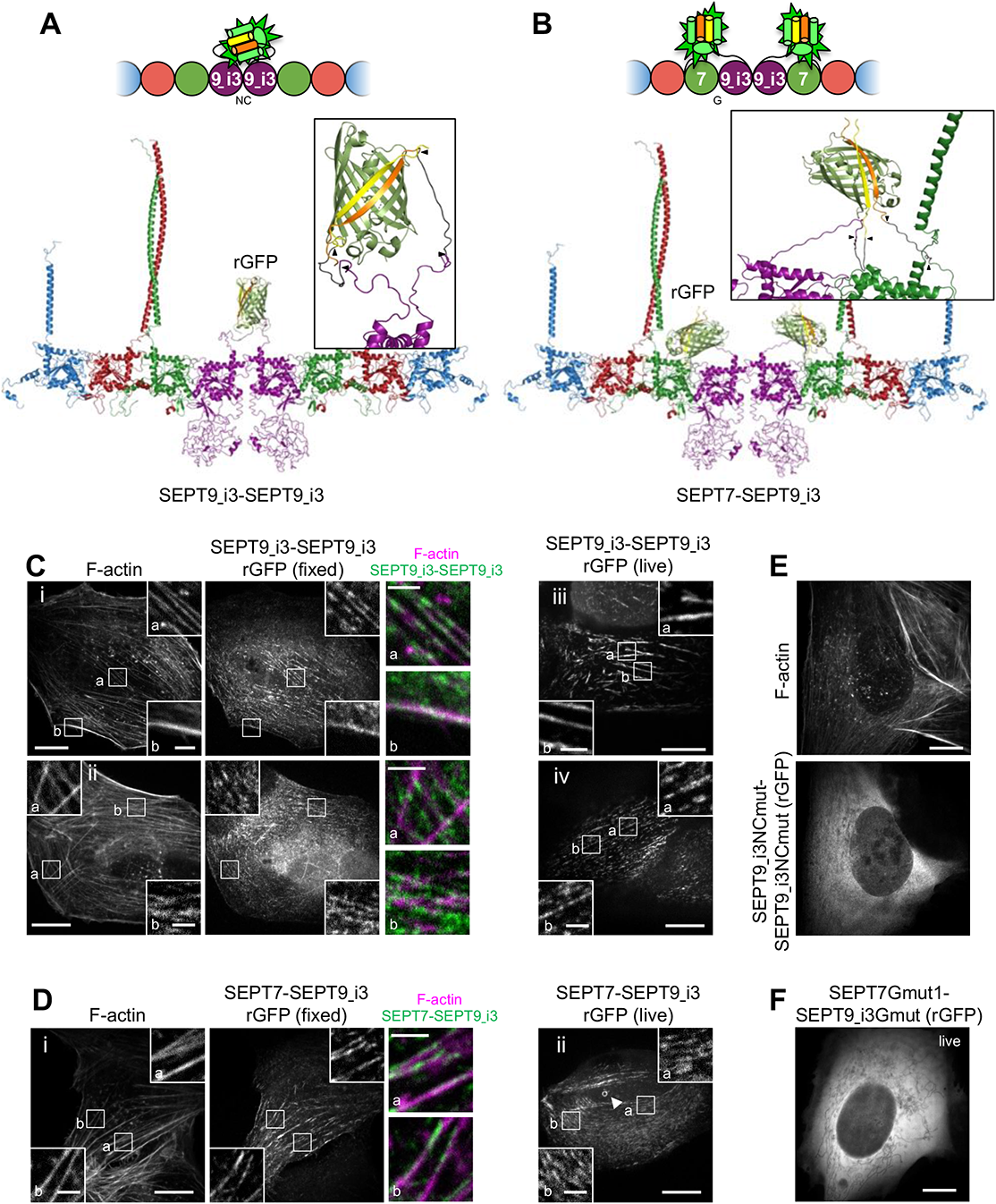
SF-associated septin filaments contain octamers. **(A-B)** Schematic (top) and respective structure model (bottom) of reconstituted GFP (rGFP) via SEPT9-SEPT9 interactions (A) and SEPT7-SEPT9 interactions (B) within an octamer. The transparency of the terminal SEPT2 subunits is used to suggest that the protomers are found within a filament. β10 and β11 strands are shown in yellow and orange, respectively. Linker sequences between septins and the β-strands, delimited by arrowheads, are shown in dark grey. The colors of septin subunits in the structure models match the ones in the color-coded sphere representation of octamers. **(C)** Representative confocal micrographs of SEPT9_i3-SEPT9_i3 rGFP distribution in live cells (right column) and in fixed cells (left and middle columns) co-stained for F-actin (phalloidin). Examples of rGFP in fixed cells localizing (i) to ventral (a) and peripheral (b) SFs and (ii) to transverse arcs (b) and excluded from dorsal SFs (a). Examples in live cells show rGFP localizing (iii) to ventral SFs (a,b) and (iv) to actin caps (a,b). Scale bars in large fields of views, 10 μm. Scale bars in insets, 2 μm. **(D)** Representative confocal micrographs of SEPT7-SEPT9_i3 reconstituted GFP (rGFP) distribution in live cells (right column) and in fixed cells (left and middle columns) co-stained for F-actin (phalloidin). Example of rGFP in fixed cells localizing (i) to ventral SFs (a,b). Example in live cells showing rGFP localizing (ii) to transverse arcs (a,b). The arrowhead points to a ring (see other examples in Fig. S6C). Such cytoplasmic rings were ∼0.5-1.6 μm in diameter (0.9 μm on average from 19 measured rings). Scale bars in large fields of views, 10 μm. Scale bars in insets, 2 μm. **(E)** Representative example of a GFP1-9 cell co-expressing SEPT9_i3NCmut-β10 and -β11 showing a diffuse cytosolic phenotype. Scale bar, 10 μm. **(F)** Representative example of GFP1-9 cell co-expressing β11-SEPT7Gmut1 and SEPT9_i3Gmut-β10 showing a diffuse cytosolic phenotype. Scale bar, 10 μm.

Expression levels of all β10/β11 fusions were kept low to minimize overexpression artifacts (Fig. S3E), and endogenous SEPT7 and SEPT9 were consistently knocked down in all subsequent experiments (Fig. S3G,H). As expected, rGFP from SEPT9_i3-SEPT9_i3 localized to contractile SFs (Fig. 3C), similarly to SEPT9 immunostainings (Fig. S1B), confirming that septin filaments contain SEPT9_i3-octamers. Split assays probing SEPT7-SEPT9_i3 interactions entirely recapitulated these findings (Fig. 3D), with rGFP additionally labeling cytoplasmic rings of ∼0.9 μm in diameter (Fig. 3Dii; Fig. S6C). To confirm that SF-localized rGFP from SEPT7-SEPT9_i3 and SEPT9_i3-SEPT9_i3 depend on the integrity of these interfaces and thus reflect direct SEPT7-SEPT9_i3 and SEPT9_i3-SEPT9_i3 interactions, we designed a double point SEPT9_i3 NC interface mutant (SEPT9_i3 M263D, I270D, hereafter SEPT9_i3NCmut) (Fig. S4A), a double point SEPT9_i3 G interface mutant (SEPT9_i3 W502A, H512D, hereafter SEPT9_i3Gmut) (Fig. S4A) and a double point SEPT7 G interface mutant (SEPT7 W269A, H279D, hereafter SEPT7Gmut1) (Fig. S4A) (Kuzmic et al., 2022; Sirajuddin et al., 2007; Zent et al., 2011). Native PAGE of these mutants with full-length GFP fusions confirmed that SEPT9_i3NCmut completely disrupts octamers (Fig. S5B), whereas SEPT7Gmut1 completely disrupts octamers and hexamers (Fig. S7B). Split assays using these mutants to disrupt the SEPT9_i3-SEPT9_i3 NC interface or the SEPT7-SEPT9_i3 G interface, completely abolished SF localization (Fig. 3E; Fig. S5C-E; Fig. 3F; Fig. S6A,G), confirming that SF localization requires intact SEPT7-SEPT9_i3 and SEPT9_i3-SEPT9_i3 interfaces. All above assays and mutants gave identical results for SEPT9_i1, confirming the presence of both SEPT9_i1- and SEPT9_i3-containing octamers in SF-associated septin filaments (Fig. S5B,F-G; Figs. S6Di,ii and S6G).

By contrast, rGFP from SEPT7-SEPT7 interactions was unexpectedly difficult to detect: although it localized to SFs (Fig. 4Ai, Bi), the majority was found on ectopic short, needle-like bundles (Fig. 4Aii, Bii), similar to the localization of full-length GFP-SEPT7 fusions (Fig. 4C; Fig. S7C). These ectopic bundles did not localize to SFs (Fig. 4Aii; Fig. S7C) and contained SEPT2 but not SEPT9 (Fig. 4C). These bundles thus most likely consist of hexamers, in line with the capacity of recombinant hexamers to form septin filament bundles *in vitro* (DeRose et al., 2020; Iv et al., 2021; Kinoshita et al., 2002; Leonardo et al., 2021). The presence of rGFP on the ectopic bundles thus showed that the split SEPT7-SEPT7 assay readily detects SEPT7-SEPT7 interactions originating from hexamers. To explore the origin of SEPT7-SEPT7 rGFP on SFs, we generated structure models of septin protomers in order to examine GFP complementation from SEPT7-SEPT7 interactions within one hexamer (Fig. 4D), as well as from SEPT7 facing another SEPT7 in apposed hexamers or octamers in a paired septin filament (Fig. 4E). Examination of the distances and the flexibility of the SEPT7 N-termini and the linkers in the structure models showed that GFP reconstitution can occur in the context of both single and paired filaments. Thus SEPT7-SEPT7 rGFP on SFs could originate either from SEPT7-SEPT7 interactions within a hexamer (Fig. 4D) or from SEPT7-SEPT7 across paired hexamers or octamers (Fig. 4E).

**Figure 4.**
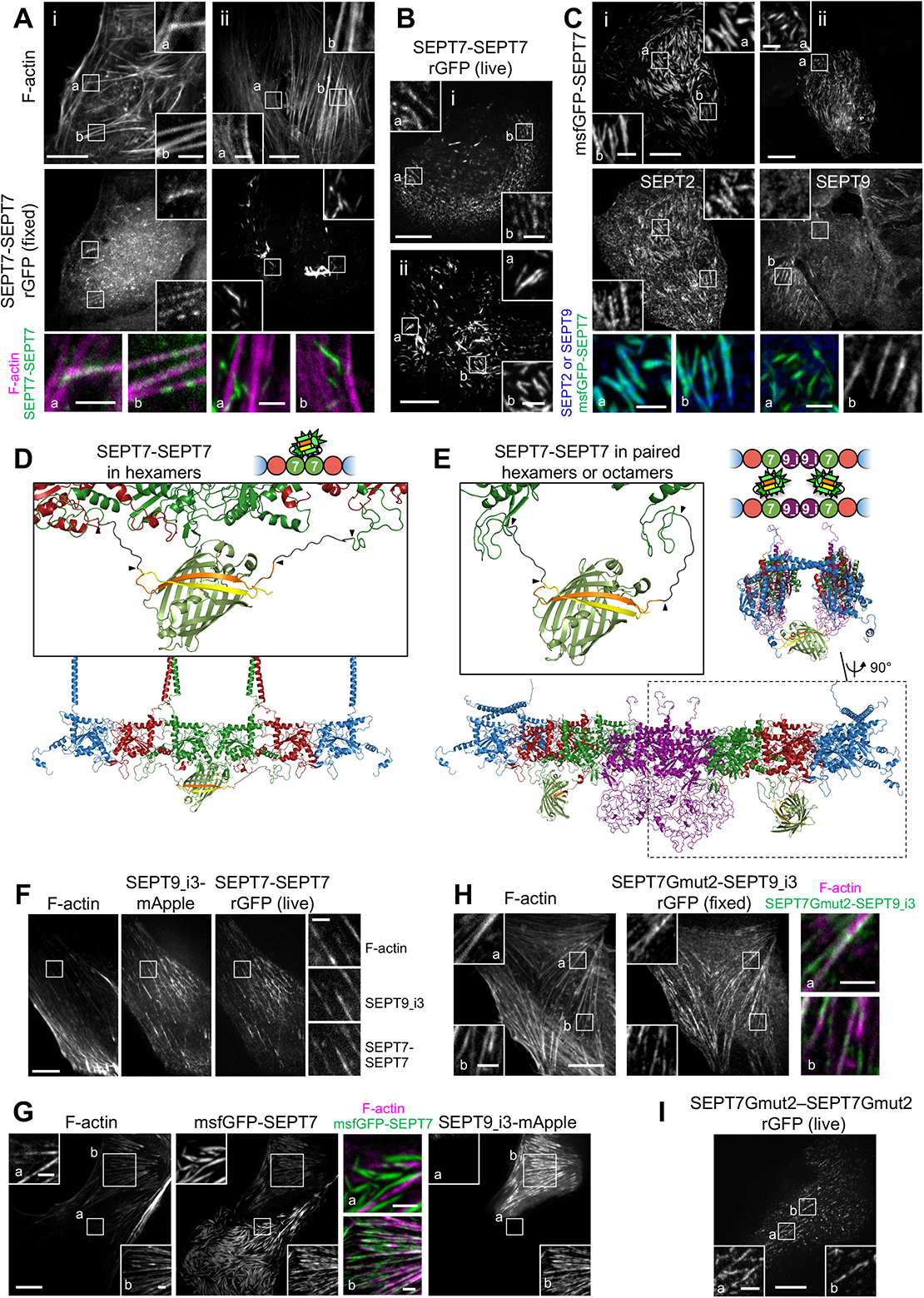
Exogenous SEPT7 and SEPT9 expression affect SEPT7 distribution. **(A)** Representative confocal micrographs of SEPT7-SEPT7 reconstituted GFP (rGFP) distribution in fixed cells co-stained for F-actin (phalloidin) localizing (i) to ventral (a,b) SFs and (ii) to ectopic bundles devoid of phalloidin staining (a,b). Scale bars in large fields of views, 10 μm. Scale bars in insets, 2 μm. **(B)** Representative confocal micrographs of SEPT7-SEPT7 reconstituted GFP (rGFP) distribution in live cells localizing (i) to transverse arcs (a,b) and (ii) to ectopic bundles (a,b). Scale bars in large fields of views, 10 μm. Scale bars in insets, 2 μm. **(C)** Examples of cells expressing msfGFP-SEPT7 and co-stained for SEPT2 (i) or for SEPT9 (ii). msfGFP-SEPT7 localizing to ectopic bundles contained SEPT2 (i;a,b) but not SEPT9 (ii;a). A non-transfected cell in (ii) shows SEPT9-stained SFs (b). Scale bars in large fields of views, 10 μm. Scale bars in insets, 2 μm. **(D-E)** Schematic (top) and respective structure model (bottom) of rGFP via SEPT7-SEPT7 interactions within a hexamer (E) or from SEPT7 in two apposed octamers within a paired filament (F). The transparency of the terminal SEPT2 subunits is used to suggest that the protomers are found within a filament. β10 and β11 strands are shown in yellow and orange, respectively. Linker sequences between septins and the β-strands, delimited by arrowheads, are shown in dark grey. The colors of septin subunits correspond to the ones in the color-coded sphere representation of hexamers and octamers. The second half of the octamer is not shown in the rotated filament pair in (F) for the sake of simplicity. Only SEPT7 subunits are shown in the zoom-in of the reconstituted GFP barrel in G for the sake of simplicity. **(F)** Representative example of a GFP1-9 cell co-expressing β10- andβ11-SEPT7, SEPT9_i3-mApple and labeled for F-actin (SiR-actin). Example shows rGFP localization to ventral SFs. Scale bars in large fields of views, 10 μm. Scale bars in insets, 2 μm. **(G)** Representative example of a cell (top right) co-expressing msfGFP-SEPT7 and SEPT9_i3-mApple and labeled for F-actin (SiR-actin). Example shows msfGFP-SEPT7 localizing to ventral SFs (b). A cell expressing only msfGFP-SEPT7 (bottom left) in (ii) shows msfGFP-SEPT7 localizing to ectopic bundles that are devoid of F-actin (a). Scale bars in large fields of views, 10 μm. Scale bars in insets, 2 μm. **(H)** Representative example of a fixed GFP1-9 cell co-expressing β11-SEPT7Gmut2 and SEPT9_i3-β10 and co-stained for F-actin (phalloidin). Example shows rGFP localizing to ventral SFs (a,b). Scale bars in large fields of views, 10 μm. Scale bars in insets, 2 μm. **(I)** Representative example of a GFP1-9 cell co-expressing β10- and β11-SEPT7Gmut2 showing rGFP localizing to SFs. Scale bars in large fields of views, 10 μm. Scale bars in insets, 2 μm.

An observation that could explain the difficulty to detect SEPT7-SEPT7 on SFs was the dependence of SEPT7 localization on SEPT9 expression levels. We consistently detected ectopic bundles when we exogenously expressed only SEPT7, either GFP-SEPT7 or split SEPT7-SEPT7 (Fig. 4A-C; Fig. S7C), but not when we co-expressed SEPT9 (Fig. 4F,G). We reasoned that in the absence of exogenous SEPT9, the slightest excess of SEPT7 leads to ectopic hexamer-based bundles, also reducing the availability of SEPT7 for forming octamers to bind SFs. SEPT7-SEPT7 interfaces being more stable than SEPT7-SEPT9 ones (Rosa et al., 2020) would facilitate this process. Exogenous co-expression of SEPT9, on the other hand, would cause incorporation of the exogenous SEPT7 into octamers, thus preventing the formation of ectopic hexamer bundles. Consistent with this hypothesis, SEPT7-SEPT7 rGFP was readily detectable on SFs under conditions of exogenous SEPT9 co-expression (Fig. 4F; Fig. S7H). Furthermore, it was difficult to find SEPT9-decorated SFs in cells also displaying ectopic hexamer-based bundles (Fig. 4G,a). These observations raised the possibility that septin filaments on SFs contain mostly, if not exclusively, octamers.

To identify the sources of the SEPT7-SEPT7 rGFP signal on SFs, we aimed at perturbing hexamers while preserving octamers. To this end, we generated a single point SEPT7 G interface mutant (SEPT7 H279D, hereafter SEPT7Gmut2) (Fig. S4A) that we reasoned should destabilize the SEPT7-SEPT7 G-interface when present in both SEPT7 subunits, but preserve the SEPT7-SEPT9 G-interface if SEPT7 is mutated but SEPT9 is wild-type. In line with these predictions, native PAGE showed that octamers are not affected by the expression of SEPT7Gmut2 (Fig. S7B), and rGFP from SEPT7Gmut2-SEPT9_i3 readily recapitulated normal septin localization on SFs (Fig. 4H; Fig. S6B,G). Importantly, rGFP from SEPT7Gmut2-SEPT7Gmut2 localized to SFs but did not show any ectopic bundles (Fig. 4I; Fig. S7G). The absence of ectopic hexamer bundles implies that SEPT7Gmut2 completely abolished SEPT7-SEPT7 interactions within hexamers in the bundles. We thus reasoned that the SF-localized rGFP from SEPT7Gmut2-SEPT7Gmut2 originates from paired octamers (Fig. 4E). We cannot exclude that the rGFP signal could originate from SEPT7Gmut2-SEPT7Gmut2 across paired hexamers containing wild-type SEPT7 and SEPT7Gmut2 subunits next to each other, but we deem this scenario highly unlikely: endogenous wild-type SEPT7 is largely knocked down (Fig. S3H), and any remaining wild-type SEPT7 would have to interact with SEPT7-β10Gmut2 that would also need to encounter SEPT7-β11Gmut2 in an apposed hexamer within a septin filament pair.

These observations altogether strongly indicate that the detected rGFP from SEPT7-SEPT7 on SFs originates from paired octamers. Split assays using the SEPT7Gmut1 mutant resulted in diffuse cytosolic distributions (Fig. S6A,G; Fig. S7A,E), confirming that SF localization requires intact SEPT7 G interfaces. These results thus support a scenario whereby septin filaments contain mostly, or even exclusively, octamers. These findings further suggest that septins organize as paired filaments, or bundles thereof.

### Septin octamers, but not hexamers, are essential for the integrity and function of SF-associated septin filaments

To further test the contribution of octamers and hexamers to septin filament formation, we examined septin filaments under three conditions: (a) the presence of hexamers and octamers (control condition), (b) the absence of octamers, by knocking down SEPT9, and (c) the presence of octamers only, by expressing SEPT7Gmut2 to disrupt hexamers while preserving octamers. As a readout of septin filaments, we examined rGFP from SEPT2-SEPT2 in live cells while at the same time imaging stress fibers (Fig. 5A-C). To assess the effects of the perturbations, we quantified the distribution of non-diffuse (SF-associated and punctate) vs. diffuse cytosolic phenotypes and calculated Pearson and Manders correlation coefficients for actin-septin co-localization in all three conditions (Fig. 5D,E). Strikingly, removing octamers by knocking down SEPT9 entirely removed the SEPT2-SEPT2 rGFP signal from all SFs, leaving behind a punctate pattern not localizing to SFs, suggesting that filamentous septin integrity depends entirely on octamers. On the other hand, preserving octamers in the absence of hexamers, through the expression of SEPT7Gmut2, preserved septin filament localization to SFs, showing that the absence of hexamers does not compromise septin filament integrity. Altogether, these results strongly support that septin filaments contain mostly, or even exclusively, octamers, with their integrity depending on octamers but not on hexamers.

**Figure 5.**
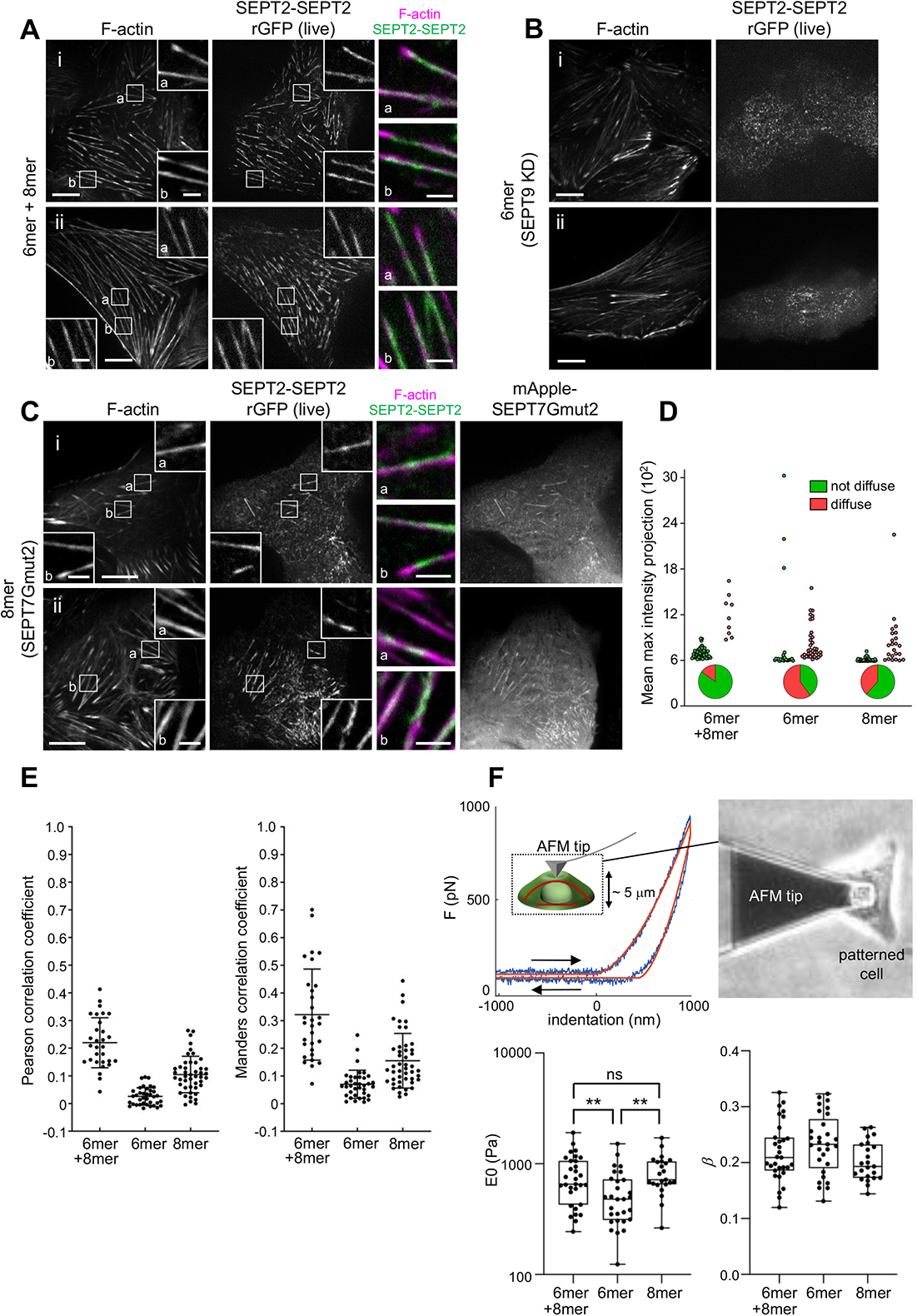
Septin octamers are essential for the integrity of SF-associated septin filaments. **(A-C)** Representative confocal micrographs of SEPT2-SEPT2 reconstituted GFP (rGFP) distribution in live cells co-labeled for F-actin (SiR-actin). Cells were treated with siRNA targeting SEPT2 (A), with siRNAs targeting both SEPT2 and SEPT9 (B), or with siRNA targeting both SEPT2 and SEPT7 and co-transfected with mApple-SEPT7Gmut2 (C). Examples in (A) and (C) show rGFP localizing to ventral SFs (a,b). Scale bars in large fields of views, 10 μm. Scale bars in insets, 2 μm. **(D)** Scatter dot plots depicting the distribution of diffuse cytosolic (red datapoints) vs. non-diffuse (green datapoints) phenotypes in cells under the same conditions as in (A-C), also shown as pie graphs. Data points are from a total of 59 cells for wild-type and 60 cells for each perturbation condition, distributed among the two phenotypes. **(E)** Scatter dot plots (mean ± SD) depicting the distributions of calculated Pearson (left) and Manders (right) correlation coefficients for actin-septin colocalization in cells under the same conditions as in (A-C). Data points for each plot, from left to right, are from a total of 30, 37 and 46 cells, respectively. **(F)** Atomic force microscopy nanoindentation on cells under the same conditions as in A-C. Top, Example of an experimental force-indentation curve. Right and left arrows correspond to the approach and retraction curves, respectively. The solid red lines represent the fits. The image on the right shows the cantilever tip indenting the dorsal membrane of a micropatterned wild-type cell. The cartoon on the left depicts the indentation of the cell, also showing ventral and dorsal SFs in red. Bottom, box plots showing the distributions of cell stiffness (*E*_0_) and cell fluidity (β). *E*_0_ values are plotted on a log scale. The data points are plotted on top of the respective box plots; each data point corresponds to one cell. On each box, the central mark indicates the median, and the bottom and top edges of the box indicate the 25th and 75th percentiles, respectively. The whiskers extend to the minimum and maximum values. The number of measurements in each box plot, from left to right, is n = 31, 29, 23. The respective median cell stiffness values are 656 Pa, 479 Pa, and 719 Pa, and the respective median cell fluidity values are 0.21, 0.23, and 0.19. One-way ANOVA for log(*E*_0_); ns=not significant; ** P<0.01.

To question the functional contribution of hexamers vs octamers in our cells, we turned to atomic force microscopy (AFM) nanoindentation for measuring cell stiffness. The principle of a nanoindentation experiment is to physically indent a cell with an AFM cantilever tip, measure the applied force, and fit the experimental force-indentation curves to a viscoelastic model in order to extract the elastic modulus (*E*_0_) and the fluidity (*β*) of the cell (Fig. 5F, see methods). Septin depletion has been previously shown to reduce cell stiffness, using AFM, in cultured mammalian cells (Mostowy et al., 2011), but the specific contribution of hexamers vs octamers to cell mechanics was not explored. To determine the contribution of hexamers vs octamers to cell stiffness, we compared nanoindentation measurements under the same conditions as earlier: (a) the presence of hexamers and octamers (control condition), (b) the absence of octamers, by knocking down SEPT9, and (c) the presence of octamers only, by expressing SEPT7Gmut2 (Fig. 5A-C). Measurements were made on cells plated on Y shape-micropatterned substrates in order to minimize variability due to size and shape differences among cells (Rigato et al., 2015). While removing hexamers did not have any effect, the depletion of octamers resulted in a statistically significant decrease in cell stiffness and a corresponding increase in cell fluidity (Fig. 5F). We conclude that octamers are essential not only for the integrity of SF-associated septin filaments, but also for their function. These data also further support the scenario that septin filaments contain mostly, or even exclusively, octamers.

### Super-resolution microscopy reveals septin fibers running longitudinally along and around SFs and interconnecting SFs

Having shown that all septins associated with SFs are filamentous, we naturally aimed at visualizing how septin filaments organize in cells on the different types of SFs. To this end, we employed super-resolution structured illumination (SIM) microscopy in cells co-stained for SEPT7 (as a pan-septin filament marker), actin filaments, and *α*-actinin or myosin-II heavy chain (MHCA). We examined septin filament organization on perinuclear actin caps, transverse arcs, including at arc nodes, on ventral SFs and at ventral actin nodes (Fig. 6A-E). Regardless of the type of SFs that septins associated with, we noticed that septin filament morphology was very different from that of the actin filament bundles in SFs. While actin filament bundles typically appeared as straight, rigid fibers throughout the cell and for different SF types, septin fibers consistently appeared less straight and with lower orientational persistence *(Note: we choose to use septin “fibers” instead of septin “filaments” in this section to avoid confusion with single or paired septin filaments or bundles thereof; we discuss the composition of septin fibers below).* Unlike core SF components like myosin and the actin crosslinker, *α*-actinin, which displayed a sarcomere-like punctate distribution (Fig. 6E, MHCA), septin fibers were distinctly separate from SFs, organizing in three manners: (a) septin fibers running longitudinally along SFs, either on the side of SFs with their signal segregated from the F-actin signal, or overlapping with SFs with the septin and F-actin signals merging (Fig. 6Aa’; Ei,e’; Eiii,f’,g’), (b) septin fibers running longitudinally along SFs and diagonally across their width, as if wrapping around the SFs (Fig. 6Ab’; Eii,h’,i’), and (c) septin fibers running longitudinally along segments of SFs while interconnecting different SFs and also forming connections to other septin fibers, thus forming a fiber meshwork (Fig. 6Cc’; Dd’; Eiv,k’). When septin fibers connected SFs, the septin fiber segments in between SFs frequently colocalized with F-actin signal, but in many instances there was no detectable F-actin signal along these segments.

**Figure 6.**
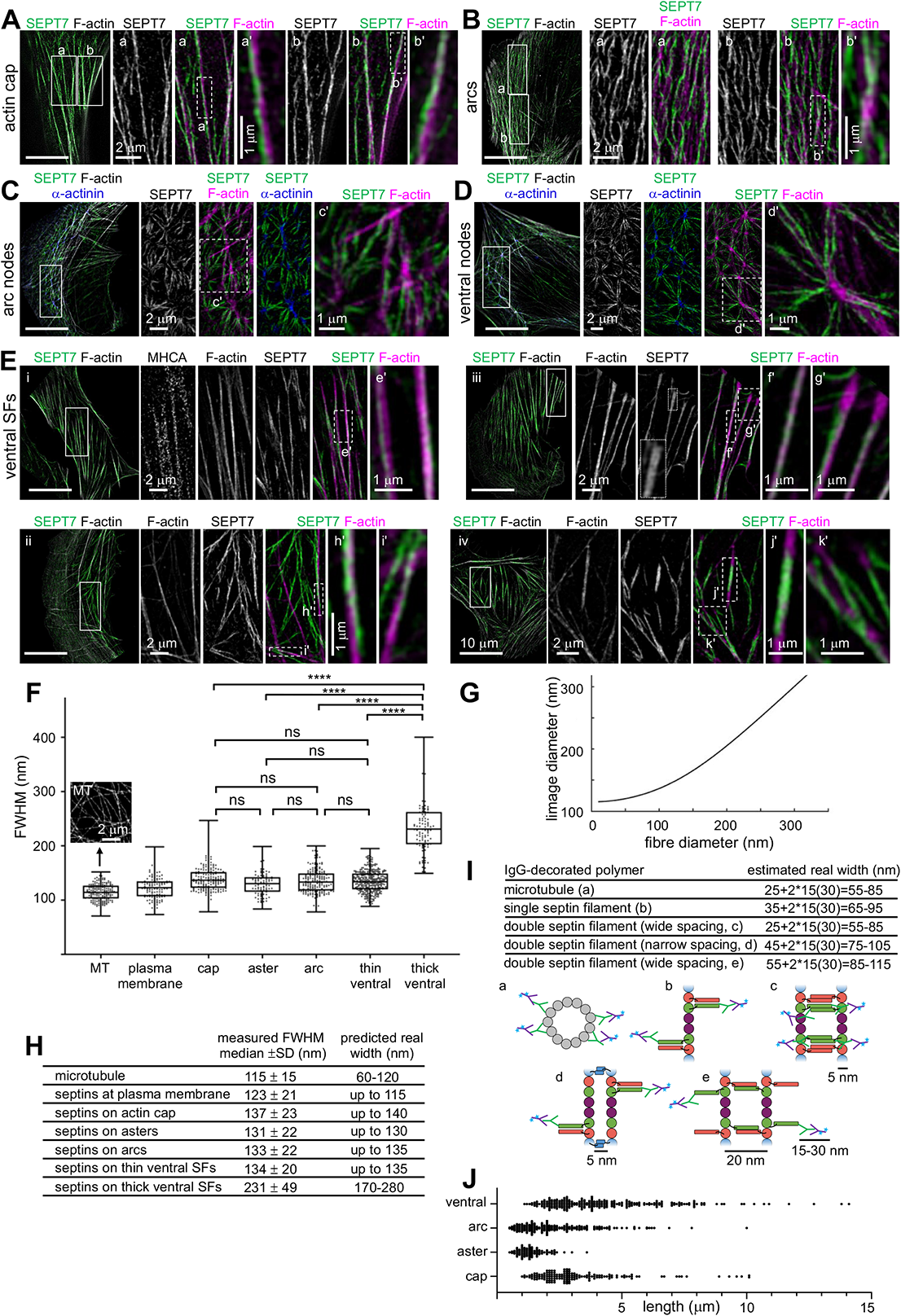
Super-resolution structured illumination (SIM) microscopy of septin filaments on SFs. **(A-D)** Representative SIM micrographs of SEPT7 immunostained cells co-stained for F-actin (phalloidin) (A-E), and additionally for *α*-actinin (C,D) or myosin heavy chain (MHCA) (E). Examples show septin filament localization to perinuclear actin caps (A), arcs (B) and arc nodes (C), ventral nodes (D), and ventral SFs (E, cells i-iv). Scale bars in all large fields of views, 10 μm. Scale bars in insets, 1 or 2 μm as indicated. **(F-I)** Fiber width measurements and real size estimations from SIM images. Box plots in (F) depict the distributions of measured widths, as the full width at half maximum (FWHM), of microtubules (MT) (an example of a SIM image of MTs is shown) and septins associated with peripheral SFs (“plasma membrane”), perinuclear actin caps (“cap”), arc and ventral actin nodes (“aster”), arcs and ventral SFs; widths from thin and thick septin fibers were plotted separately. The data points are plotted on top of the respective box plots; data points correspond to width measurements at multiple positions along MT and septin fibers and in multiple MT and septin fibers per cell in a total of 10 cells for MT and 10 cells for septin fiber measurements. On each box, the central mark indicates the median, and the bottom and top edges of the box indicate the 25th and 75th percentiles, respectively. The whiskers extend to the minimum and maximum values. The number of measurements in each box plot, from left to right, is n = 180, 123, 175, 88, 184, 330, 114. The respective median values are 115 nm, 123 nm, 137 nm, 131 nm, 133 nm, 134 nm, and 231 nm. Kruskal-Wallis test; ns=not significant; **** P<0.0001. (G) Numerical simulations of the expected FWHM in SIM images (“image diameter”) as a function of the real fiber diameter. A Gaussian point spread function (PSF) of 115 nm was used in the shown graph. The curve was generated from the convolution of the PSF with an increasing fiber size. Fiber sizes above ∼200 nm scale linearly with the image sizes. These simulations were used together with FWHM measurements in SIM images (F) to estimate an upper width limit for septin fibers associated with the different types of SFs (H). These estimations were then compared to the real width ranges one expects from IgG antibody-decorated septins organizing as single or double filaments (I). Primary and fluorophore (cyan asterisk)-conjugated secondary antibodies are depicted in green and magenta, respectively. The primary SEPT7 antibody used in our immunostainings binds the very C-terminus of SEPT7. The narrow and wide spacings of paired filaments, the presence of homodimeric coiled coils for SEPT2, SEPT6 and SEPT7, and of heterodimeric coiled coils for SEPT6 and SEPT7 are based on experimental evidence from (de Almeida Marques et al., 2012; Leonardo et al., 2021; Low and Macara, 2006; Sala et al., 2016). **(J)** Scatter dot plots of length distributions for septin fibers on the indicated types of SFs. Bars depict median values. The number of measurements in each plot, from left to right, is n = 151, 97, 227, 249. The respective median values are 2.8 μm, 1.3 μm, 2.0 μm, and 3.8 μm.

Regardless of the type of SFs, the majority of septin fibers appeared thinner than their associated SFs. We noticed that septin fibers were often thicker on the SF segments adjacent to FAs (Fig. 6Eiii,g’; Eiv,j’), but thicker septin fibers were occasionally also found on arcs, caps and ventral SFs. These thicker septin fibers did not exceed the width of the associated SF, and appeared either as single thick fibers, or what looked like two closely-apposed thin fibers (dashed rectangle in the SEPT7 channel of Fig. 6Eiii). To compare septin fiber thicknesses across the different SF types we measured the width of septin fibers at multiple positions along their length and in multiple septin fibers for each SF type. The full width at half maximum (FWHM) was calculated from the fluorescence intensity profiles of the measured widths (Fig. 6F). Thick septin fiber widths from ventral SFs, where thicker septin fibers were found more frequently, were plotted separately. The comparison of the width distributions showed that all thin septin fiber populations were very similar to each other, with median FWHM values in the range of 123-137 nm across the different SF types, and distinct from the thick ones that showed an almost 2-fold higher median FWHM value of 231 nm (Fig. 6F,H). There was no statistically significant difference between thin septin fiber widths on caps, asters, arcs and ventral SFs (Fig. 6F).

In an effort to determine whether the septin fibers are single or paired septin filaments (“double septin filaments”), or bundles thereof, we compared the width values with the widths of microtubules in the same cells. FWHM values of single microtubules (MTs), which are 25-nm wide tubes, are routinely used as the gold standard for assessing the performance of super-resolution microscopy techniques. MTs were stained using whole primary and fluorophore-coupled secondary IgG antibodies, just like for septin stainings, leading to an estimated real MT width of ∼60 nm (Fig. 6I) (Weber et al., 1978). Measurements of MT widths in our cells with SIM resulted in an average FWHM value of 115 nm, in line with reported FWHM values for MTs by SIM (Hamel et al., 2014; Wegel et al., 2016) given that the lateral resolution of SIM is roughly half of the diffraction limit, i.e., ∼110 nm. Given that the observed size in our images is the convolution of the real object size with the point spread function (PSF) of the SIM microscope, we simulated the predicted image size as a function of the real fiber size (Fig. 6G) (see methods). The comparison of the estimated real widths of primary and secondary IgG-decorated septins, assumed to organize as single or as paired filaments with either narrow (∼5 nm) or wide (∼20 nm) spacing (Leonardo et al., 2021) (Fig. 6I), with the widths predicted from our FWHM measurements of immunostained septins (Fig. 6H), suggests that the thin septin fiber widths are compatible with single or paired septin filaments, whereas the thick septin fibers could correspond to two single or two double septin filaments.

We also wondered about the length of the SF-associated septin fibers. Although our results show that all septins are filamentous, the lateral resolution limit of SIM did not allow us to distinguish if what appears as continuous fiber signal originates from a single fiber or from adjacent fibers partially overlapping at their ends. The uniform thickness across thin septin fibers let us speculate that these could correspond to single septin fibers. Length measurements of presumably single septin fibers showed that short septin fibers associated with and interconnecting actin nodes were on the order of 0.5-3.5 μm in length, whereas septin fibers on arcs, actin caps and ventral SFs were as short as 0.5-1μm in length and as long as 10-15 μm (Fig. 6J).

### SF-associated septin filaments are closely apposed to the plasma membrane

Having shown that all SF-associated septins are filamentous and given the extensive, intimate association of septin filaments with actin fibers observed by SIM, we wondered how septin function relates to septins being filamentous. Recombinant human septins can bind and cross-link actin filaments, but can also bind lipid membranes, raising the hypothesis that septin filaments in cells anchor SFs to the plasma membrane. Although it is often assumed that human septins are found at the plasma membrane, there is no formal proof of direct septin-membrane binding in cells. Mutants of the putative membrane-binding polybasic stretch of residues in the septin *α*0 helices (Cavini et al., 2021; Zhang et al., 1999) also disrupt the respective NC interfaces and thus the integrity of the protomers (Bertin et al., 2010; Kuzmic et al., 2022), and thus do not allow us to conclude on direct septin-membrane interactions. A first indication that SF-associated septins might be membrane-bound came from live cell extraction experiments. Short (30s-1 min) incubations of live cells with low concentrations of Triton X-100 detergent right before fixation is routinely used for removing cytosolic pools of cytoskeletal proteins, for example myosin, while preserving pools that are stably associated with the actin cytoskeleton, with the aim to reduce diffuse cytosolic signal and enhance the signal on SFs to reveal better its sarcomere-like distribution. Strikingly, while extracting the plasma membrane after fixation entirely preserved septin localization to SFs, live-cell extraction removed septins from all SFs while preserving actin, myosin and the actin crosslinker, *α*-actinin, on SFs (Fig. 7A, data not shown for *α*-actinin). Septins thus did not seem to behave like core components of SFs such as myosin and actin crosslinkers, and their sensitivity to the detergent suggested that they might be bound to the membrane.

**Figure 7.**
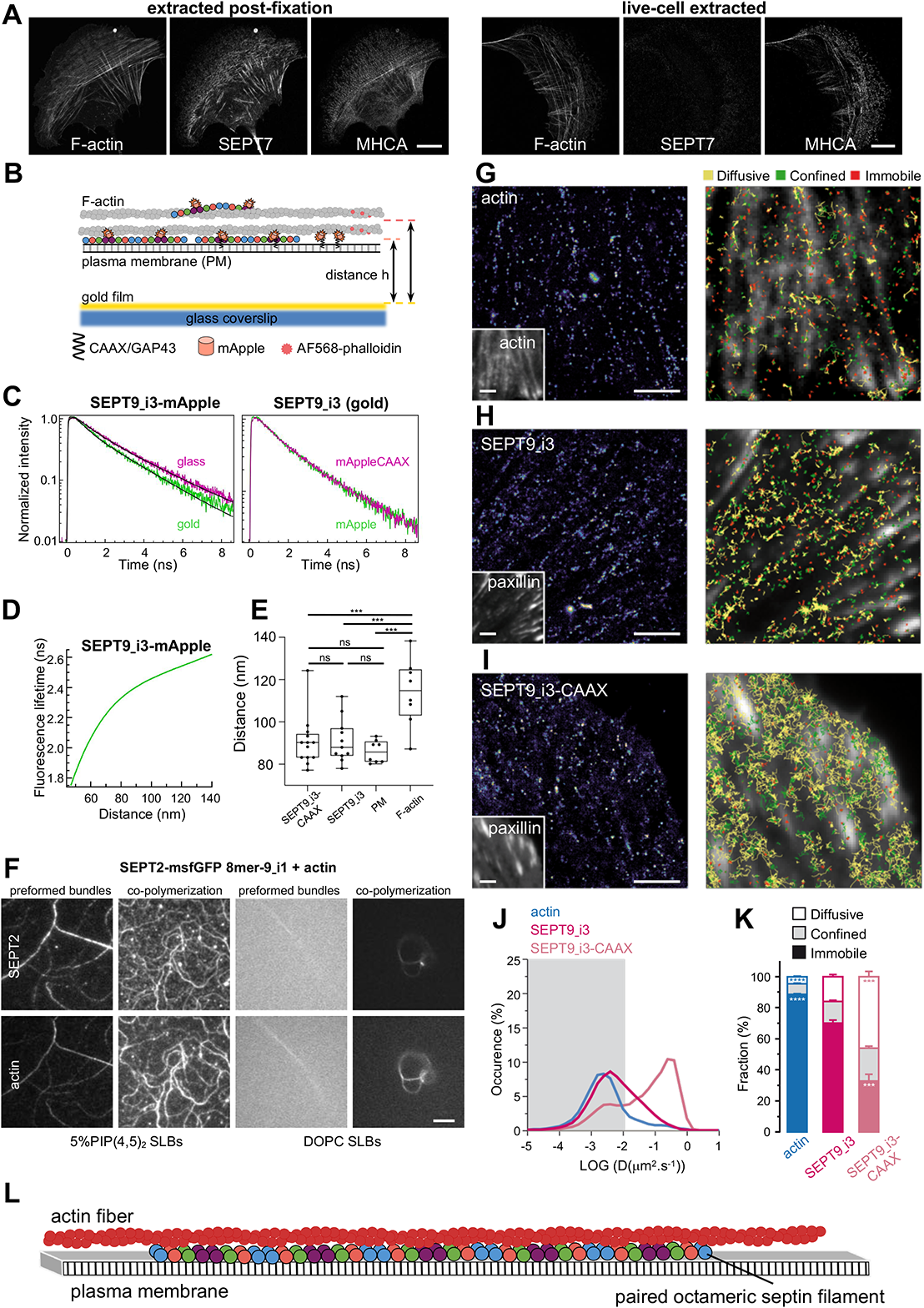
Septin filaments are closely apposed to the plasma membrane, are largely immobilized on actin stress fibers, and can mediate actin-membrane anchoring. **(A)** Representative confocal micrographs of SEPT7 immunostained cells co-stained for F-actin and myosin heavy chain (MHCA). Cells were either extracted after fixation (left panel) or were live-extracted right before fixation (right panel). Scale bars, 10 μm. **(B)** Schematic of the metal-induced energy transfer (MIET) assay for probing fluorophore (mApple or AlexaFluor 568) distances from a gold-coated coverslip using fluorescence lifetime measurements. **(C-E)** C depicts representative examples of lifetime decay traces for SEPT9_i3-mApple on glass and in the presence of gold (left) and for SEPT9_i3-mApple and SEPT9_i3-mApple-CAAX in the presence of gold (right). The solid lines represent the numerical fits, showing the lifetime reduction due to the MIET process. The calculated lifetime-distance dependence for SEPT9_i3-mApple (D, see methods) was used to calculate the distance of SEPT9_i3-fused mApple, with or without the CAAX lipid anchor, from the coverslip (E). Lifetime decay traces and lifetime-distance dependence curves for GAP43-mApple (plasma membrane) and AF568-phalloidin (F-actin) are shown in Fig. S8A,B. Box plots in (E) depict the distributions of calculated distances for SEPT9_i3-mApple-CAAX, SEPT9_i3, GAP43-mApple (plasma membrane, PM) and AF568-phalloidin (F-actin). The data points are plotted on top of the respective box plots; each data point corresponds to one cell. On each box, the central mark indicates the median, and the bottom and top edges of the box indicate the 25th and 75th percentiles, respectively. The whiskers extend to the minimum and maximum values. The number of measurements in each box plot, from left to right, is n = 13, 11, 8, 8. The respective median values are 90 nm, 88 nm, 86 nm, and 115 nm. One-way ANOVA; ns=not significant; *** P<0.001. **(F)** TIRF images of SEPT2-msfGFP 8mer-9_i1 and F-actin, either co-polymerized on top of a supported lipid bilayer (SLB), or co-polymerized in solution to form preformed bundles that were then flushed onto the supported lipid bilayer. The supported lipid bilayer was composed either of 5% of PI(4,5)P_2_, a septin-interacting lipid, and 95% DOPC (top panels), or 100% DOPC (bottom panels). Due to the shallow penetration depth (∼100 nm) of TIRF together with the absence of crowding agents, only truly membrane-associated structures are visible. Scale bar, 5μm. **(G-K)** Septins are primarily immobilized and confined in actin stress fibers but also undergo very slow lateral free diffusion in the vicinity of the plasma membrane. (G-I) Left: Super-resolution PALM intensity images of mEos2-Actin (G), SEPT9_i3-mEos3.2 (H) and SEPT9_i3-mEos3.2-CAAX (I) in mouse embryonic fibroblasts obtained from a sptPALM sequence (50 Hz, >80 s). Insets: low resolution images of GFP-actin (G) or GFP-paxillin (H-I), which were co-expressed for FA labelling. Scale bars, 3 µm. Right: color-coded trajectories overlaid on FAs labelled by GFP-paxillin or on FAs and SFs labelled by GFP-actin (grayscale) show the diffusion modes: free diffusion (yellow), confined diffusion (green) and immobilization (red). (J) Distributions of the diffusion coefficient D computed from the trajectories of mEos2-actin (blue), SEPT9_i3-mEos3.2 (magenta) and SEPT9_i3-CAAX-mEos3.2 (light magenta) obtained outside FAs, are shown in a logarithmic scale. The gray area including D values inferior to 0.011 µm².s^-1^ corresponds to immobilized proteins. Values represent the average of the distributions obtained from different cells. (K) Fraction of mEos2-actin (blue), SEPT9_i3-mEos3.2 (magenta) and SEPT9_i3-mEos3.2-CAAX (light magenta) undergoing free diffusion, confined diffusion or immobilization outside FAs. Values represent the average of the fractions obtained from different cells (error bars: SEM). Results for SEPT9_i3-mEos3.2 (14 cells) correspond to pooled data from two independent experiments with n, the number of trajectories analyzed: SEPT9_i3-mEos3.2 n_SEPT9_i3_ = 72,720. Results for mEos2-actin (9 cells) and SEPT9_i3-mEos3.2-CAAX (5 cells) correspond each to data from one experiment with n, the number of trajectories analyzed: mEos2-actin n_actin_ = 34,715; SEPT9_i3-mEos3.2-CAAX n_SEPT9_i3-CAAX_ = 37,339. Statistical significance in (K) was obtained using two-tailed, non-parametric Mann–Whitney rank sum test. The different conditions were compared to the SEPT9_i3-mEos3.2 condition. The resulting P values are indicated as follows: *** P<0.001; **** P<0.0001. **(L)** Working model supported by the results of this study. Septins in cells organize as paired, octamer-based filaments mediating actin-membrane anchoring.

To directly test if septins on SFs are close to the plasma membrane, we employed a metal-induced energy transfer (MIET) assay in cells (Chizhik et al., 2014). In MIET, the fluorescence lifetime is dependent on the distance of fluorophores from a metal layer, allowing us to use fluorophore lifetime measurements for deducing the axial distance of fluorophores from a gold-coated coverslip surface with an axial resolution of a few nanometers (Fig. 7B). We hypothesized that septins could either associate with the plasma membrane while interacting with SFs, or that septins interact with SFs in the absence of any septin-membrane association. To distinguish these scenarios, we compared distances of the fluorescent protein, mApple, in three conditions: (a) mApple N-terminally fused to the 20 N-terminal residues of neuromodulin/GAP43 that contains palmitoylated cysteines and functions as a membrane targeting signal (GAP43-mApple); this fusion thus served as a reference for fluorophores localizing directly at the plasma membrane, (b) SEPT9_i3-mApple as a reference for ventral SF-associated septin octamers, and (c) SEPT9_i3-mApple-CAAX as a reference for septins targeted to the plasma membrane through the H-Ras CAAX motif, i.e. the 20 C-terminal residues of H-Ras that contain palmitoylated and farnesylated cysteines and function as a membrane targeting signal. Representative lifetime decay traces are shown in Fig. 7C and Fig. S8A,B. Strikingly, the distance of mApple from the metal surface, derived from the lifetime-distance dependence curve (Fig. 7D and methods), was the same for SF-associated septins, membrane-bound mApple, and membrane-bound septins, strongly indicating that septins are closely apposed to the plasma membrane (Fig. 7E). Lifetime measurements of AF568-phalloidin bound to ventral SFs under the same conditions placed SFs significantly further away, by ∼25 nm, from the plasma membrane (Fig. 7E). MIET assays being limited to probing interactions within 200 nm from the metal surface, it was not feasible to probe septin populations on SFs localized further away, notably transverse arcs and perinuclear actin caps. Given that arcs and actin caps are most likely coupled to the dorsal plasma membrane (Burnette et al., 2014; Maninova et al., 2017), it is conceivable that septins associated with these SF pools are also membrane-bound.

### Septin filaments anchor actin filaments to lipid membranes

Since SF-associated septins are closely apposed to the membrane, we naturally wondered if septin filaments could function to anchor stress fibers to the plasma membrane. In the absence of available septin membrane-binding and actin-binding mutants, we turned to reconstitution assays on supported lipid bilayers (SLBs), comparing phosphatidylcholine-(PC) vs phosphatidylinositol(4,5)-bisphosphate (PI(4,5)P_2_)-containing membranes (Fig. 7F; Fig. S8C), PI(4,5)P_2_ being a septin-interacting lipid (Szuba et al., 2021). To image only truly membrane-associated structures, we used total internal reflection fluorescence (TIRF) microscopy in the absence of crowding agents. Actin filaments alone did not bind lipid membranes, whereas septin octamers alone specifically bound PI(4,5)P_2_-containing membranes (data not shown), in line with previous reports for mammalian septin hexamers (Szuba et al., 2021). To test if septins can anchor actin to membranes, we either preformed actin-septin bundles in solution and then added them to SLBs, or co-polymerized septins and actin on SLBs. In both cases, and specifically on PI(4,5)P_2_-containing membranes but not on membranes composed of PC, actin filaments and actin filament bundles were anchored to the lipid bilayers (Fig. 7F; Fig. S8C), showing that septin filaments can indeed at the same time bind membranes and actin and thus mediate membrane-actin anchoring.

### Single protein tracking reveals that septins are immobilized on actin stress fibers

Since septin filaments are able to anchor actin filaments to lipid membranes in vitro and given that SF-associated septins are in close proximity to the plasma membrane, we questioned whether the molecular dynamics of septins at the plasma membrane could reveal the potential SF-anchoring function of septins in living cells.

To determine the molecular behavior of septins in the vicinity of the plasma membrane and actin stress fibers, we combined photoactivated localization microscopy (PALM) with live-cell single protein tracking of SEPT9_i3 and actin fused to photoswitchable mEos fluorescent proteins (mEos3.2 and mEos2, respectively) using sptPALM (Manley et al., 2008; Rossier et al., 2012). Cells were co-transfected with mEos-fused proteins and GFP-paxillin as a FA reporter or GFP-actin as a SF and FA reporter. In brief, this method uses TIRF microscopy to detect and track numerous sparse photo-activated proteins of interest within 200 nm above the coverslip surface at high-frequency (50Hz acquisition), allowing us to reconstruct thousands of protein trajectories (Fig. 7G,H,I). For trajectories lasting at least 260 ms, we compute the mean square displacement (MSD), which describes the diffusion properties of a molecule. We then sort trajectories according to their diffusion modes (immobile, confined, free-diffusive; Fig. 7K), and extract diffusion coefficients (D_diff_, D_conf_) (Fig. 7J; Fig. S8D,E; see methods). We first looked at the dynamic behavior of mEos2-actin in SFs labelled with GFP-actin (Fig. 7G). mEos2-actin was found inside FAs and also linearly organised along SFs between FAs, as expected. Actin mostly displayed immobilized and confined behaviors, as illustrated by the large fractions of immobilization and confined diffusion (Fig. 7G,K; immobile: 88.5 ± 0.5%, confined: 6.8 ± 0.3%, mean ± SEM) and a distribution of diffusion coefficients centered around 1.5-2.5.10^-3^ µm^2^.s^-1^ (Fig. 7J). In line with septin immunostainings (Fig. 1Bi,a; Fig. S1Ai,c; Fig. S1Bi,c; Fig.S2D), single SEPT9_i3-mEos3.2 molecules were rarely found inside FAs, but were linearly organized between FAs decorating SFs (Fig. 7H). Like actin, also SEPT9_i3-mEos3.2 was found to be primarily immobilized and confined (Fig. 7H,K; immobile: 70.0 ± 1.9%, confined: 13.9 ± 0.8%, mean ± SEM). Contrary to actin, however, SEPT9_i3-mEos3.2 also displayed a significant freely diffusing population (Fig. 7H,K; diffusive: 16.1 ± 1.3%, mean ± SEM). However, septin free-diffusion was very slow (Fig. 7J; Fig. S8D) with a diffusion constant D_diff_ = 0.087 ± 0.001 µm^2^.s^-1^ (mean ± SEM) that is comparable to that of free diffusing transmembrane proteins (integrins: (Rossier et al., 2012)) or of a lipid-anchored protein bound to the plasma membrane by its PH domain (kindlin:(Orre et al., 2021). Confined SEPT9_i3-mEos3.2 was also diffusing very slowly (Fig. S8E) with a diffusion constant D_conf_ = 0.044 ± 0.001 µm^2^.s^-1^ that was comparable to that of mEos2-actin (0.057 ± 0.003 µm^2^.s^-1^; means ± SEM). Overall these results suggest that septins, when immobilized and confined, could indeed be anchoring actin SFs to the plasma membrane, while the free-diffusing septins display a diffusivity that is consistent with them being membrane-anchored.

Similarly to the MIET experiments, we used SEPT9_i3-mEos3.2-CAAX as a reference for septins targeted to the plasma membrane. SEPT9_i3-mEos3.2-CAAX did not localize specifically to SFs but decorated the whole plasma membrane (Fig. 7I). In comparison with the behavior of SEPT9_i3-mEos3.2, SEPT9_i3-mEos3.2-CAAX displayed a smaller immobilized fraction (Fig. 7I,J,K; immobile: 32.6 ± 0.5%) but increased free-diffusion and confined diffusion fractions with an increased diffusion constant (D_diff_: 0.328 ± 0.002 µm^2^.s^-1^; D_conf_: 0.202 ± 0.002 µm^2^.s^-1^, mean ± SEM) (Fig. S8D,E). Interestingly, being stably anchored to the plasma membrane allowed SEPT9_i3-mEos3.2-CAAX to diffuse inside FAs (Fig. 7I). The much lower diffusion coefficient of SEPT9_i3-mEos3.2 compared to that of SEPT9_i3-mEos3.2-CAAX is in line with freely diffusing SEPT9_i3 being fully incorporated into septin filaments (our results from the split assays) and could reflect hop diffusion of septin filaments alternating between SFs and the plasma membrane. Altogether, our combined findings from SLB assays, MIET and sptPALM in cells support a scenario in which septin filaments in cells function by anchoring SFs to the plasma membrane (working model in Fig. 7L).

### Microtubule-associated septins organize as filaments containing predominantly octamers

While SF-associated septin filaments contain both SEPT9_i1 and SEPT9_i3-octamers, only octamers containing SEPT9_i1 associate with microtubules (Kuzmic et al., 2022). Co-stainings of septins and MTs in U2OS cells revealed occasionally septin localization to MTs (Fig. 8A), in line with the very low amounts of the SEPT9_i1 isoform in this cell line (Kuzmic et al., 2022). Split-GFP complementation assays probing SEPT9_i1-SEPT9_i1 and SEPT7-SEPT9_i1 interactions, together with mutants disrupting the SEPT9_i1-SEPT9_i1 NC interface and the SEPT9_i1-SEPT7_i1 G interface confirmed that MT localization requires intact SEPT7-SEPT9_i1 and SEPT9_i1-SEPT9_i1 interfaces (Fig. 8B; Fig. S5F-I; Figs. S6Diii,iv and S6E,G). To test whether all septins on MTs also organize as filaments that contain mostly, if not exclusively octamers, we probed SEPT2-SEPT2 interactions on MTs and also selectively removed hexamers using SEPT7Gmut2. To facilitate the study of MT-associated septins in U2OS cells, we increased their amount by co-expressing SEPT9_i1 (Kuzmic et al., 2022; Nagata et al., 2004; Nagata et al., 2003; Surka et al., 2002). Examining the rGFP from SEPT2-SEPT2 in live cells, while at the same time imaging MTs, revealed rGFP signal decorating MTs, revealing that MT-associated septins are indeed filamentous (Fig. 8C). The SEPT2 NC interface mutant in the context of the split SEPT2-SEPT2 assay completely abolished MT localization, confirming that an intact SEPT2-SEPT2 NC interface and thus direct SEPT2-SEPT2 interactions are required for septin localization to MTs (Fig. 8D). These results, together with the fact that SEPT2NCmut-mApple fusions do not associate with MTs (Kuzmic et al., 2022), lead us to conclude that all septins decorating MTs are filamentous.

**Figure 8.**
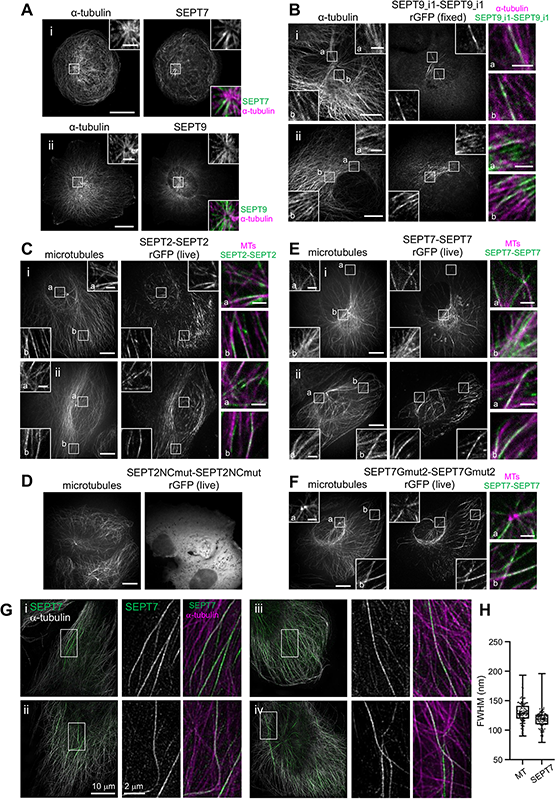
Septins on microtubules organize as octamer-based filaments. **(A)** Representative confocal micrographs of SEPT7 (i) SEPT9 (ii) immunostained cells co-stained for microtubules (*α*-tubulin). Examples show septins localizing to microtubules close to the microtubule organizing center. **(B)** Representative confocal micrographs of SEPT9_i1-SEPT9_i1 reconstituted GFP (rGFP) distribution in fixed cells (i, ii) co-stained for microtubules (*α*-tubulin). **(C)** Representative confocal micrographs of SEPT2-SEPT2 rGFP distribution in live cells (i, ii) co-expressing mCherry-SEPT9_i1 (not shown) and labeled for microtubules (SiR-tubulin). **(D)** Representative confocal micrograph of a GFP1-9 cell co-expressing SEPT2NCmut-β10 and -β11, mCherry-SEPT9_i1 (not shown) and labeled for microtubules (SiR-tubulin), showing a diffuse cytosolic phenotype. **(E)** Representative confocal micrographs of SEPT7-SEPT7 rGFP distribution in live cells (i, ii) co-expressing mCherry-SEPT9_i1 (not shown) and labeled for microtubules (SiR-tubulin). **(F)** Representative example of a GFP1-9 cell co-expressing β10- and β11-SEPT7Gmut2 co-expressing mCherry-SEPT9_i1 (not shown) and labeled for microtubules (SiR-tubulin). (A-F) Scale bars in large fields of views, 10 μm. Scale bars in insets, 2 μm. **(G)** Representative SIM micrographs of cells (i-iv) expressing mCherry-SEPT9_i1 (not shown) co-stained for SEPT7 and *α*-tubulin. Scale bars in large fields of views, 10 μm. Scale bars in insets, 2 μm. **(H)** Box plots depict the distributions of measured widths, as the full width at half maximum (FWHM), of microtubules (MT) and MT-associated septins (SEPT7). The data points are plotted on top of the respective box plots; data points correspond to width measurements at multiple positions along MT and septin fibers and in multiple MT and septin fibers per cell in a total of 8 cells. On each box, the central mark indicates the median, and the bottom and top edges of the box indicate the 25th and 75th percentiles, respectively. The whiskers extend to the minimum and maximum values. The number of measurements is n = 128 and 112 for MTs and septins, respectively. The respective median values are 128 nm and 119 nm for MTs and septins, respectively.

Although SEPT9_i1 is essential for septins to bind MTs (Kuzmic et al., 2022), whether hexamers are important for septin filament localization on MTs is not known. In line with our results on exogenously expressed SEPT7, it was difficult to find SEPT9-decorated MTs in cells also displaying ectopic hexamer-based bundles (Fig. S7D,a). rGFP from SEPT7-SEPT7 was, however, readily detected on MTs under our conditions of exogenous SEPT9_i1 co-expression (Fig. 8E), suggesting that the reconstituted split-GFP signal may originate from paired octamers (Fig. 4E). When we expressed SEPT7Gmut2 to selectively disrupt hexamers while preserving octamers, rGFP from SEPT7Gmut2-SEPT7Gmut2 and SEPT7Gmut2-SEPT9_i1 were readily detected on MTs (Fig. 8F; Fig. S6F), reflecting indeed SEPT7-SEPT7 rGFP from paired octamers (Fig. 4E), and also showing that hexamers are not essential for septin-MT association, in line with our findings for actin-associated septin filaments. SIM imaging of MT-associated septin filaments revealed thin septin fibers running along MTs over several micrometers (Fig. 8G). Different from the presence of both thin and thick actin-associated septin fibers, all MT-associated septin fibers we observed appeared homogeneous in their width. FWHM measurements of MT and septin fiber widths (Fig. 8H), combined with numerical simulations for estimating the real fiber size in SIM images (Fig. 6G), predicted MT and septin fiber widths in the ranges of 65-135 nm and 85-130 nm, respectively. These predictions are compatible with MT-associated septin filaments organizing as paired septin filaments (Fig. 6I).

## Discussion

We showed that human septins in U2OS cells extensively associate with myosin-II containing contractile SFs, but not with non-contractile dorsal SFs nor with focal adhesions. Combining a tripartite split-GFP complementation assay to probe SEPT2-SEPT2 interactions as a molecular readout for septin polymerization with mutants that disrupt SEPT2-SEPT2 end-to-end polymerization, we showed that all septins decorating SFs organize as filaments. Moreover, complementation assays probing SEPT9-SEPT9 and SEPT7-SEPT9 interactions, together with mutants disrupting the SEPT9-SEPT9 NC interface and the SEPT9-SEPT7 G interface, showed that septin filaments rely on intact octamers. The use of distinct SEPT7 G interface mutants, which either disrupt both the SEPT7-SEPT7 and the SEPT7-SEPT9 G interfaces, or that disrupt specifically the SEPT7-SEPT7 but not the SEPT7-SEPT9 G interface, allowed us to explore the composition of septin filaments and distinguish the contributions of hexamers vs octamers to septin filament integrity and function. Our results showed that septin filaments contain mostly, or even exclusively, octamers, and that octamers, but not hexamers, are essential for the integrity of septin filaments and their function in cell stiffness. Super-resolution structured illumination microscopy revealed that septin filaments organize as fibers running longitudinally along and around SFs, interconnecting SFs and forming a meshwork. Careful analysis of the filament widths in SIM images, using microtubules as a reference, revealed that thin septin fibers most likely correspond to paired septin filaments. To understand the function of septin filaments on SFs, we examined septin localization with respect to the plasma membrane. Nanometer-resolved distance measurements showed that septin filaments associated with ventral SFs are closely apposed to the plasma membrane, in line with reconstitution assays on lipid membranes showing that septins can mediate membrane-actin anchoring and with single protein tracking measurements revealing that septins are largely immobilized and confined on actin fibers. Finally, our results showed that all microtubule-associated septins also organize as filaments whose integrity depend on octamers but not on hexamers.

Our study shows for the first time that all SF-associated septins organize as filaments mediated by direct SEPT2-SEPT2 end-to-end polymerization. Whether septins organize invariably as filaments when associated with other actin-based structures in cells and tissues, notably at the cell cortex, at cell-cell junctions and in the cytokinetic ring, can now be tested formally using the tools we have developed in this study, namely a combination of split-GFP assays and polymerization-disrupting mutants. Given the high conservation of residues in the NC interface, our mutants and split assays are well placed for being adapted for other septins within the SEPT2 group when required by cell- and tissue-specific septin expression (Karlsson et al., 2021; Uhlen et al., 2015). Importantly, split assays can test if septins organize as filaments *in situ* in cellular contexts not necessarily linked to actin, and can also be used as readouts for filament formation in the context of future studies aiming to identify regulators of septin polymerization, drugs interfering with septin polymerization, or septin mutants related to disease. These tools can further be adapted for studying septin organization in a tissue context in genetic animal model systems.

An unexpected result of our study is the fact that human septin filament integrity in cells depends entirely on octamers, raising the possibility that septin filaments contain predominantly, if not exclusively, octamers, and naturally questioning the functional importance of hexamers. The embryonic lethality of *Sept9* mouse knock-outs (Fuchtbauer et al., 2011) might point to the essential, if not exclusive, contribution of octamers to septin function. Our single point mutant (SEPT7 H279D) that disrupts hexamers while preserving octamers promises to help elucidate the relative functional contribution of hexamers vs octamers in other cellular contexts. Unlike SEPT3 and SEPT12, whose expression is restricted to specific tissues, SEPT9 is expressed in practically all tissues and cell lines (Karlsson et al., 2021; Uhlen et al., 2015), thus octamers are expected to be found ubiquitously. We speculate that it is the SEPT3 group septin, which is absent from hexamers, that dictates septin function; for example, SEPT9_i3 would mediate septin-actin interactions, and SEPT9_i1 would drive septin-microtubule interactions. The fact that recombinant hexamers can also bind actin and membranes (Iv et al., 2021; Szuba et al., 2021) is most likely due to actin- and membrane-binding domains on septins being conserved across septins from different groups and might reflect a fundamental capacity of human septins to interact with these scaffolds; in the presence of the SEPT3 group septin, additional physiological signals, for example the very N-terminus of SEPT9_i1 important for septin-MT association (Kuzmic et al., 2022), would work together or in addition to the actin- and membrane-binding domains for determining septin localization and function. It is also possible that the actin/membrane binding of hexamers observed *in vitro* does not happen in cells. The observation that SEPT7 is the septin whose localization and assembly are most sensitive to SEPT7 and SEPT9 expression levels, and the fact that SEPT7-SEPT7 interactions are stronger than SEPT7-SEPT9 ones (Rosa et al., 2020) might suggest that SEPT9, and possibly the other SEPT3 group septins, helps prevent SEPT7 from forming ectopic bundles. It is intriguing that *Drosophila* does not have any SEPT3 group septins and thus contains only hexamers (Field et al., 1996). Interestingly, endogenous *Drosophila* septins in cells occasionally form cytoplasmic tube-like bundles devoid of Anillin, in addition to their normal localization to the cortex in interphase cells and to the cleavage furrow in dividing cells (Hickson and O’Farrell, 2008). Such cytoplasmic bundles also form in the absence of Anillin’s septin-binding domain that recruits septins to the plasma membrane (Kechad et al., 2012). We speculate that the formation of these cytoplasmic bundles is analogous to the ectopic hexamer bundles we observed in human cells and that they form in similar conditions, i.e., in the absence of, or at limiting amounts of, a physiological partner (SEPT9 in our case, Anillin for *Drosophila*).

Our findings suggest that actin-associated septin filaments in mammalian cells organize as paired septin filaments, with thicker septin fibers, e.g. close to FAs, likely consisting of 2-3 double septin filaments. Recombinant and cell-isolated septin protomers in solution and on membranes form single and paired filaments, as well as straight, curved and interconnected bundles of varying thicknesses (DeRose et al., 2020; Iv et al., 2021; Soroor et al., 2021; Szuba et al., 2021). Septins at the budding yeast neck organize in arrays made of single and paired filaments (Bertin et al., 2012; Ong et al., 2014); bundles of more than two filaments were not observed. However, single-molecule localization microscopy of SF-associated septins in U2OS cells, i.e., in the same system we use in our study, reported that septin bundles may comprise as few as 25 to as many as 150 filaments (Vissa et al., 2019). However, we note that this study assumed septin GTP-binding domains associating laterally without any spacing, without considering the ∼5-20 nm spacing occupied by coiled-coil pairing used for bundling (Leonardo et al., 2021), so the number of filaments could be substantially overestimated. The SIM images of SEPT2-GFP in cancer-associated fibroblasts (Calvo et al., 2015) showing thin septin fibers along SFs are in line with our results that septin fibers contain a few rather than tens of filaments. This same study also reported septins appearing to wrap around actin fibers. These observations support the hypothesis that human septins in cells, like budding yeast septins, organize in a rather narrow range of assembly geometries. We speculate that the wide range of assembly geometries found for septins in solution reflects their plasticity, but that the presence of a physiological partner, like actin filaments and the plasma membrane, leads to their native assembly into paired filaments (Bertin et al., 2010; Ong et al., 2014; Szuba et al., 2021), but not to bundles of more than a few filaments, as indicated in our data (Fig. 6H,I).

Our results from the MIET assays on ventral SF-associated septins showed that septin filaments are closely apposed to the plasma membrane. We also found septins localized along stretches of peripheral SFs and at curved segments of the membrane that appeared devoid of phalloidin staining (Fig. 6Eiii; Fig. S2B), but these observations are not enough to conclude on direct septin-membrane binding. The strongest evidence, in our opinion, that septins are membrane-bound in cells, or at least closely apposed to the plasma membrane, comes from electron microscopy evidence in budding yeast (Bertin et al., 2012; Ong et al., 2014; Rodal et al., 2005). Our finding that recombinant septin octamer-containing filaments simultaneously bind lipid membranes and actin filaments, thus providing actin-membrane anchoring, leads us to speculate that all SF-associated septins in cells bind actin fibers to the adjacent plasma membrane. Although detectable free-diffusion of septins with sptPALM suggests septin-membrane association, whether septins bind membranes directly or indirectly through membrane-associated interacting partners remains to be shown. It will be interesting to explore if cortical actin meshworks are also membrane-attached via septins.

Mammalian septins are distinct and unique from other membrane-bound actin-binding and -crosslinking proteins, like Anillin and ERM proteins, in that they form filaments. Their capacity to form filaments, catalyzed by membrane binding (Szuba et al., 2021) and coupled with their ability to bind and cross-link actin filaments (Iv et al., 2021; Mavrakis et al., 2014), provides them with the unique potential to stabilize actin filament bundles and meshes at the plasma membrane over considerable distances (working model in Fig. 7L). The important fractions of immobilized and confined septins on SFs from sptPALM are in line with such a stabilization function. This hypothesis is further consistent with our SIM imaging showing that septins are intimately associated with actin SFs over many micrometers and also interconnect contractile SFs forming meshes. We propose that septins function precisely by binding ventral SFs, transverse arcs and perinuclear actin caps to the respective ventral and dorsal plasma membrane, stabilizing them at the membrane and at the same time interconnecting the respective SFs, participating in their generation or/and maintenance. Our AFM data are in line with such a role, whereby septins stiffen the cell cortex. Although septins are absent from FAs, multiple reports showed that their absence impacts FA maturation (Calvo et al., 2015; Dolat et al., 2014; Kang et al., 2021). Our observation that septin filaments associate as thick fibers adjacent to FAs lets us think that they contribute a stabilization function at the connection between FAs and the core of SFs and thus impact FA maturation indirectly by affecting the accumulation of mechanical tension on SFs.

The findings of this study lead to several remaining open questions. In the absence of the identification of actin-binding domains on septins, it is still unclear if SF decoration by septins in cells reflects direct septin-actin interactions, or if such binding occurs through myosin-II or/and Borg proteins (Calvo et al., 2015; Farrugia et al., 2020; Joberty et al., 2001; Joo et al., 2007; Liu et al., 2014; Mostowy et al., 2010; Salameh et al., 2021). The fact that septins are found only on contractile SFs lets one suppose that myosin-II related signaling might be involved in their recruitment to SFs but this remains to be shown. Also what regulates septin polymerization in cells is still unknown. Cell-free assays have suggested cooperativity in septin-actin binding (Iv et al., 2021), which might drive SF generation and/or maintenance. Cell-free reconstitution approaches and animal model systems promise to provide important further insights into the link between animal septin organization and function.

## Materials and methods

### Design of septin fusions for the tripartite split-GFP complementation assay

For the tripartite complementation assay to report SEPT-SEPT interactions with stringency, the amino acid linker length between SEPT and the β10- and β11-strands should not be too short in order to allow for the necessary proximity and flexibility for the β10- and β11-strands to orient in an antiparallel fashion for complementing GFP1-9, but it should be short enough to minimize reporting longer-range interactions. We used fluorescence imaging to test the dependence of split-GFP complementation on the linker length and on the position of the β10- and β11-tags by screening different homo- and hetero-septin combinations as shown in Fig. S3A-C. All the combinations we tested resulted in fluorescence, reflecting the inherent flexibility of the N- and C-termini of SEPT2, 7 and 9. To allow for the most stringent complementation, we chose to use C-terminal fusions with 14-residue linkers for SEPT2-β10- and -β11 tags and for SEPT9-β10- and -β11 tags, and N-terminal fusions with 14-residue linkers for β10- and β11-SEPT7 tags. This short linker is comparable in length to the 10-residue-long β10- and β11-strands and thus long enough to allow the antiparallel arrangement of the latter. Protein structure models of human septin hexamers and octamers bearing full-length β10- and β11-tagged septins (see method section “Modeling of human septin complexes”) confirmed the efficiency of GFP complementation for the final chosen linker length and β10/β11-tag positioning (Fig. 2B,C; Fig. 3A,B; Fig. 4D,E).

### Plasmids and cloning

Septin and msfGFP cDNAs were as described in (Iv et al., 2021). mApple and sfCherry2 cDNAs were PCR-amplified from Addgene plasmids #54862 and #83031, respectively. Three types of mammalian expression plasmids were used in this study. A pCMV backbone (Clontech) was used for the expression of full-length fluorescent protein (msfGFP/mApple) fusions. A pcDNA3.1 backbone (ThermoFisher Scientific), also with a CMV promoter, was used for the expression of β10-or β11-tagged septins. Finally, a pTRIP TRE Bi vector, modified from pTRE-Tight-BI (Takara-Bio) (Koraichi et al., 2018), bearing a bidirectional tetracycline response element (TRE) promoter and an IRES-TagBFP cassette downstream β10-tagged septins, was used for the doxycycline-inducible co-expression of β10- and β11-tagged septins (Fig. S3D,E). pCMV and pTRIP TRE Bi plasmids were used for all results presented in the figures. The pcDNA3.1 plasmids were used only for the initial screening (Fig. S3A-C).

All pCMV plasmids, SEPT2 constructs in pTRIP TRE Bi plasmids and all interface mutants in pTRIP TRE Bi plasmids were cloned using seamless cloning (In-Fusion HD Cloning Plus Kit from Takara Bio, 638910). All pcDNAs and all wild-type SEPT7 and SEPT9-containing constructs in pTRIP TRE Bi plasmids were generated with classical cloning. In this latter case, DNA fragments were amplified by PCR using the PCR Master Mix from ThermoFisher Scientific (K0171), TaqFast DNA polymerase (Applied Biological Materials G277) or Phusion High-Fidelity DNA Polymerase (New England Biolabs M0530S) and ligated into double digested plasmids with the Rapid DNA Ligation Kit from ThermoFisher Scientific (K1422). pCMV constructs were cloned into a NheI/BamHI linearized vector. pcDNA constructs were cloned into a NheI/XbaI linearized vector. pTRIP TRE Bi constructs were cloned in two steps: first the β10-tagged septins were cloned into a SacII/NheI digested vector, then the β11-tagged septins were cloned into a NdeI/XbaI digested vector carrying the β10-tagged septin. The starting methionine of septin sequences is included in the N-terminal β10- and β11-tagged versions.

Bacterial expression plasmids for generating wild-type SEPT2-msfGFP hexamers and octamers-9_i3 were described in (Iv et al., 2021) and are available through Addgene (#174492, 174498, 174499, 174501). pnEA-vH plasmids for the bacterial expression of SEPT2NCmut-msfGFP and SEPT2-sfCherry2 were generated using seamless cloning following the same strategy described in (Iv et al., 2021). All primers for seamless cloning were Cloning Oligo (<60 bp) or EXTREmer (>60 bp) synthesis and purification quality from Eurofins Genomics, Germany and are listed in Table S1. All restriction enzymes were FastDigest enzymes from Thermo Scientific or from New England Biolabs. All plasmids were verified by sequencing (Eurofins Genomics, Germany) after each cloning step. We have deposited all plasmids with the nonprofit repository Addgene. Note that the SEPT9_i1NCmut in this study is the same as the SEPT9_i1NCmut2 in (Kuzmic et al., 2022). Plasmid mCherry-SEPT9_i1 was from Addgene (#71622).

### Cell lines, cell culture and transfection

U2OS osteosarcoma cells for the expression of full-length fluorescent protein septin fusions were from ATCC (HBT-96). For the inducible co-expression of β10- and β11-tagged septins in the context of the tripartite split-GFP complementation system, we generated an inducible U2OS-Tet-On-GFP1-9 cell line which expresses constitutively a GFP1-9 fragment and an anti-GFP VHH intrabody that enhances split-GFP fluorescence. To generate this cell line, U2OS cells were successively transduced with lentiviruses encoding rtTA, GFP1-9 and anti-GFP VHH G4 and tested for complementation efficiency using transient expression of a GFP10-zipper-GFP11 domain (Koraïchi et al., 2018). One additional round of transduction with GFP1-9 lentivirus lead to an optimized U2OS-Tet-On-GFP1-9 cell line that showed 80% GFP positive cells upon expression of the GFP10-zipper-GFP11 domain. An IRES-TagBFP cassette downstream β10-tagged septins was used for monitoring septin expression. Cells were maintained in McCoy’s medium (Gibco 16600082) supplemented with 10% fetal bovine serum (Dominique Dutscher S181H), 100 U/mL penicillin and 100 μg/mL streptomycin antibiotics (P4333, Sigma) in a humidified atmosphere at 37°C containing 5% CO_2_.

Transfections with pcDNAs, for the screening of β10- and β11-tag combinations (Fig. S3A-C), were performed 16 h prior to immunostainings using jetPRIME (PolyPlus 101000015). To obtain single cells for imaging, 50×10^3^ U2OS-Tet-On-GFP1-9 cells were typically grown on 18 mm coverslips (Knittel Glass MS0010), previously cleaned by sonication in 70% ethanol, and placed into a 12-well plate a day prior to the day of transfection, for allowing an optimal number of cells to attach and spread. A total of 0.4 μg of DNA and a 4:1 ratio of jetPRIME (μL) : DNA (μg) were used per reaction. To minimize septin overexpression artifacts, the total amount of DNA was composed by 30 ng of β10-septin, 30 ng of β11-septin and 340ng of empty vector.

Transfections with either pCMV or pTRIP TRE Bi plasmids and siRNAs were performed through electroporation using the Neon Transfection System (Thermo Fisher Scientific MPK5000). For pCMVs, a single 100-μL reaction using 1.8×10^6^ U2OS cells, 300 pmol of each siRNA and 6 μg of each DNA was electroporated within the dedicated tip (Thermo Fisher Scientific, MPK10096). Electroporation parameters consisted in 4 pulses of 10 ms width and a voltage of 1230 V. The electroporated cells were then inoculated in 5 mL of culture medium without antibiotics, and immediately divided for native-PAGE, SDS-PAGE/western blots and immunostaining as follows: 3 mL in a 6-cm dish containing 2 mL of medium without antibiotics, 2 mL in a 6-cm dish containing 3 mL of medium without antibiotics, and 100 μL in the well of a 12-well plate containing a 18-mm coverslip in 900 μL of medium without antibiotics, respectively. A satisfactory septin knockdown efficiency was achieved within 48-96 h after electroporation. Typically, immunostaining and protein extraction were performed 72h post electroporation.

For pTRIP TRE Bi plasmids, a single 100-μL reaction using 1.8×10^6^ U2OS-Tet-On-GFP1-9 cells, 300 pmol of each siRNA and 6 μg of each DNA was electroporated within the dedicated tip using the same electroporation parameters described previously. The electroporated cells were then inoculated in 5.5 mL of culture medium without antibiotics and immediately divided for SDS-PAGE/western blots, immunostaining and live cell imaging as follows: 5 mL in a 6-cm dish, 400 μL in the well of a 12-well plate containing a 18-mm coverslip in 600 μL of medium without antibiotics, and 200 μL in the well of a 24-well glass bottom plate (Cellvis, P24-1.5H-N) containing 800 μL of medium without antibiotics, respectively. After either 48 h, for samples intended for live cell imaging or immunostainings, or 72 h, for samples for biochemical analysis, protein expression was induced using 1 μg/mL of doxycycline (Sigma, D9891) for 16 h.

### Septin mutant phenotype classification

The diffuse cytosolic vs non-diffuse phenotype classification analysis for mutant characterization with pCMV plasmids (Fig. 2I; Fig. S5C,F; Fig. S7A) was done from 3 independent experiments. Transfected cells were fixed and co-stained for actin and α-tubulin. Each round of experiments was composed of the 30 first fluorescent cells found randomly in the sample, with the exception of one round containing 11 cells for msfGFP-SEPT7 and 8 cells for msfGFP-SEPT7Gmut1. Acquired images were classified as “diffuse cytosolic” in the presence of purely diffuse cytosolic signal or as “non-diffuse” in the presence of structure-like signal; no differentiation was applied for SF-, microtubule-, membrane-like or punctate signals in the latter case. The violin graphs representing the phenotype distributions show the mean intensity distribution calculated on the whole field of view from maximum intensity projections of all z-planes. The phenotype classification for split constructs was identical, in terms of the used criteria and graph display, but the data was generated from 2 independent experiments, with each experimental round composed of the 20 first fluorescent cells, in live cell imaging, with the exception of one round containing 13 cells for SEPT7Gmut1-SEPT7Gmut1 and 9 cells for SEPT7Gmut2-SEPT7Gmut2. For the septin-actin colocalization analysis, the diffuse cytosolic vs non-diffuse phenotype sorting is displayed both as scatter dot plots and as a pie graph to highlight the diffuse cytosolic vs non-diffuse proportion from each condition. Bars in scatter dot plots depict means and error bars SD. Violin plots, scatter dot plots and pie graphs were prepared using GraphPad Prism. The number of cells used to assess the phenotypes for each condition is indicated in the respective legends.

### RNA interference

Control synthetic small interfering RNA (siRNA) targeting the coding region of LacZ (5’-GCGGCUGCCGGAAUUUACC-3’) and siRNA targeting the 3’UTR region of all SEPT9 mRNA variants (5’-GGAUCUGAUUGAGGAUAAA-3’) were previously validated (Verdier-Pinard et al., 2017). The siRNA sequences targeting the 3’UTR regions of SEPT2 and SEPT7 were 5’-ACACUUUCCUGGAUAAAAA-3’ and 5’-GCAUUUAGCUGUAUUCAUA-3’, respectively. All siRNAs were designed to hybridize with 19-bp sequences in the 3’UTR regions of septin genes, thus knocking down endogenous septins while allowing the expression of the transfected plasmids. 21mer siRNAs, 20 nmol each, were synthesized with dTdT overhangs by Eurofins, and delivered as annealed and ready-to-use siRNA duplexes.

### SDS-PAGE and western blotting of cell lysates

The dish containing the cells was placed on ice and the cells were washed twice with PBS, without Ca^2+^ and Mg^2+^, before being detached with 40 μL of ice-cold lysis buffer (10 mM Tris-HCl pH 7.5, 148.5 mM NaCl, 0.5 mM EDTA, 0.5% NP-40, 1x PhosSTOP Roche, 5x cOmplete protease inhibitor cocktail Roche, 1 mM DTT) using a cell scraper (TPP 99003). The lysate was collected in a 1.5 mL tube and incubated on ice for 30 min. The lysates were then centrifuged at 20,000 g for 20 minutes at 4°C for removing cell debris. An aliquot of 6 μL was collected for protein quantification using the BCA Protein Assay (ThermoFisher Scientific 23227) and the remaining clarified lysates were kept at −20°C until SDS-PAGE analysis.

The lysates were analyzed by 4-20% SDS-PAGE using Mini-PROTEAN TGX^TM^ Precast Protein Gels (BioRad 4561095). Molecular mass markers were Precision Plus Protein All Blue Standards (BioRad 1610373) or Amersham ECL Rainbow Marker (Cytiva RPN800E). For the western blot, the gel, the PVDF Immobilon-P^SQ^ membrane (MERCK ISEQ85R), filter pads and filter papers were all incubated in transfer buffer (25 mM Tris, 192 mM glycine and 20% of methanol) for 15 min before assembly in the Mini Trans-Blot transfer cell (BioRad 1703935). The transfer was done at 4°C for 16 h at 110 mA constant current. The transfer efficiency was checked by Ponceau S staining (Sigma P7170). The membrane was then blocked in a 3% w/v dry milk TBS-T solution (20 mM Tris-HCl pH7.5, 200 mM NaCl and 0.1% v/v Tween20) for 90 min under constant agitation. Primary and secondary antibodies were diluted in the same blocking solution and incubated over the membrane for 90 and 60 min, respectively. In between antibody incubations, membranes were washed three times for 10 min with TBS-T, and the very last wash right before ECL detection was done only with TBS.

The loaded amount of extracted protein in the gels was adapted depending on the expression promoter and the analyzed septin. For pCMV plasmids used to assess the knockdown efficiency (Fig. S3F-H), a total of 4 μg of extracted protein was used for detecting endogenous SEPT2, 8 μg for endogenous SEPT7, and 4 μg for endogenous SEPT9. For pTRIP TRE Bi plasmids, a total of 8 μg of extracted protein was used for all analysis. To detect specific septins, we used rabbit anti-SEPT2 (1:2500, Sigma HPA018481), rabbit anti-SEPT7 (1:200, Santa Cruz Biotechnology sc-20620) and rabbit anti-SEPT9 (1:4000, Proteintech 10769-1-AP). For detecting β10- and β11-tag expression, we used rabbit anti-β10 (1:5,000) and rabbit anti-β11 (1:5,000) (Koraichi et al., 2018). For detecting tubulin as a loading control, we used mouse anti-α-tubulin (1:2,500, Sigma T9026). Secondary HRP-conjugated antibodies were either anti-rabbit-IgG (1:10,000, Cytiva GENA934) or anti-mouse-IgG (1:10,000, Cytiva GENA931). Chemiluminescent detection was performed with an Amersham ImageQuant 800 imager (Cytiva 29399481) using Amersham ECL Select Western Blotting Detection Reagent (Cytiva RPN2235) diluted five times in Milli-Q water. The membrane was incubated with the diluted reagent for 30 s, and washed for 10 s in TBS right before image acquisition. Images were collected in time series mode every 10 s, for a total of 50 images, and processed with ImageQuantTL software for quantification of the band intensities to measure expression levels. Expression quantification graphs for assessing knockdown efficiency were prepared using GraphPad Prism and are shown as mean values (normalized to 1 for siCtrl) with the error bar representing the standard deviation. Data are from at least 3 independent siRNA treatments.

### Native PAGE and western blotting of cell lysates

The dish containing the cells was placed on ice and the cells were washed twice with PBS, without Ca^2+^ and Mg^2+^, before being detached with 40 μL of ice-cold native lysis buffer (80 mM PIPES pH 6.9, 2 mM MgCl_2_, 4 mM EGTA, 0.2% saponin, 5x cOmplete protease inhibitor cocktail Roche). The lysate was collected in 1.5 mL tube and incubated on ice for 10 min. The lysates were then centrifuged at 14,000 g for 10 min at 4°C for removing cell debris. To prevent septin polymerization, clarified lysates were supplemented with NaCl, adding 10 μL of NaCl 5 M for each 100 μL of lysate. After 15 min of incubation on ice, the lysates were clarified in a second centrifugation step of 10 min, 14,000 g at 4°C. An aliquot of 12 μL was collected for protein quantification using the BCA Protein Assay (ThermoFisherScientific 23227), and the remaining clarified lysates were kept at −20°C until Native PAGE analysis.

The lysates were analyzed by 4-16% Native PAGE using precast Bis-Tris Mini Protein Protein Gels (Invitrogen BN1003BOX) following the manufacturer’s instructions. The molecular mass marker was NativeMark™ Unstained Protein Standard (Invitrogen, LC0725). For the western blot, the gel, the PVDF Immobilon-P^SQ^ membrane, filter pads and filter papers were all incubated in NuPAGE transfer buffer for 15 min before assembly. The transfer was done at 4°C for 16 h at 20 V constant voltage. The transfer efficiency was checked by destaining the membrane with an aqueous solution containing 25% of methanol and 10% of acetic acid. The protein marker was identified and the membrane completely destained with pure methanol for 3 min. The membrane was then blocked and stained with the respective antibodies as described for SDS-PAGE western blots.

The loaded amount of extracted protein in the gels was again adapted depending on the analyzed septin. A total of 10 μg of extracted protein was used for detecting endogenous or exogenously expressed SEPT2 and SEPT7, and 10 or 4 μg for endogenous or exogenously expressed SEPT9, respectively. To detect specific septins, we used mouse anti-SEPT2 (1:7,500, Proteintech 60075-1), rabbit anti-SEPT7 (1:200, Santa Cruz Biotechnology sc-20620) and rabbit anti-SEPT9 (1:2,000, Proteintech 10769-1-AP). Secondary HRP-conjugated antibodies were either anti-rabbit IgG (1:10,000, Cytiva GENA934) or anti-mouse IgG (1:10,000, Cytiva GENA931). Chemiluminescent detection was done with an Amersham ImageQuant 800 imager (Cytiva 29399481) using Amersham ECL Select Western Blotting Detection Reagent (Cytiva RPN2235) as described previously for SDS-PAGE western blot.

### Immunofluorescence

Cells were fixed for 15 min with 4% paraformaldehyde (Electron Microscopy Sciences 15714) in 37°C-prewarmed cytoskeleton buffer (10 mM MES pH 6.1, 150 mM NaCl, 5 mM EGTA, 5 mM MgCl_2_, 5 mM glucose), followed by 2 x 5 min wash steps in phosphate-buffered saline (PBS) solution, and a subsequent permeabilization and blocking step with PBS containing 0.1% saponin and 1% IgG-free/protease free bovine serum albumin (BSA) (Jackson ImmunoResearch 001-000-161) for 1 h at RT. Cells were incubated successively with primary antibodies for 16 h at 4°C in a humidified chamber, followed by secondary Alexa Fluor-conjugated IgG antibodies combined with 0.165 μM Alexa Fluor 647-phalloidin (ThermoFisher Scientific A22287) for 2 h at RT. Antibody solutions were prepared in PBS containing 0.1% saponin and 1% BSA, and 3 x 10 min wash steps in the same buffer were performed in between antibody incubations. Coverslips with stained cells were washed 2 x 5 min in PBS and then mounted with 15 μL Fluoromount (Sigma F4680) for image acquisition. Primary antibodies were rabbit anti-SEPT2 (1:500, Sigma HPA018481), rabbit anti-SEPT7 (1:500, IBL 18991), rabbit anti-SEPT9 (1:200, Proteintech 10769-1-AP), mouse anti-*α*-tubulin (1:10,000, Sigma T9026), mouse anti-paxillin (1:500, Merck Millipore 05-417). Secondary antibodies were donkey AlexaFluor488-conjugated anti-rabbit IgGs (1:500, Thermo Fisher Scientific A10037) and donkey AlexaFluor568-conjugated anti-mouse IgGs (1:500, Thermo Fisher Scientific A21206).

### Immunostaining after live-cell extraction vs after extraction post-fixation

To live-extract cells (Fig. 7A), we incubated cells in 37°C-prewarmed cytoskeleton buffer containing 0.1% v/v Triton X-100 for 1 min, then replaced immediately with 37°C-prewarmed cytoskeleton buffer containing 4% paraformaldehyde (Alfa Aesar 43368) and fixed cells for 15 min. Cells were rinsed with PBS and incubated in a permeabilization/blocking solution of PBS containing 0.1% saponin and 5% goat serum (ThermoFisher Scientific 16210064) overnight at 4°C. Cells were incubated successively with primary antibodies for 2 h at RT in a humidified chamber, followed by secondary Alexa Fluor-conjugated IgG antibodies combined with 0.165 μM Alexa Fluor 546-phalloidin (ThermoFisher Scientific A22283) for 1 h at RT. Antibody solutions were prepared in PBS containing 0.1% saponin and 5% goat serum, and 3 x 10 min wash steps in the same buffer were performed in between antibody incubations. Coverslips with stained cells were washed 2 x 5 min in PBS and then mounted with Fluoromount-G (Southern Biotech 0100-01) for image acquisition. Primary antibodies were rabbit anti-SEPT7 (1:400, IBL 18991) and mouse anti-non-muscle Myosin IIA (1:200, abcam ab55456). Secondary antibodies were goat AlexaFluor488-conjugated anti-mouse IgGs (1:400, Thermo Fisher Scientific A11001) and goat AlexaFluor633-conjugated anti-rabbit IgGs (1:400, Thermo Fisher Scientific A21070). The respective control experiment, i.e. extracting cells post-fixation (Fig. 7A), involved fixing cells in 37°C-prewarmed cytoskeleton buffer containing 4% paraformaldehyde for 15 min, then extracting cells with PBS containing 0.5% Triton X-100 and 5% goat serum for 10 min before overnight permeabilization/blocking and antibody incubations as described above. This last protocol was also used for the immunostainings shown in Fig. 1Biv, Fig. 6 and Fig. S2 using the additional primary antibodies mouse anti-*α*-actinin-1 (1:200, Thermo Scientific clone BM 75.2) and mouse anti-vinculin (1:200, clone hVin-1 Sigma V9264). Cells shown in Fig. 1Biv, Fig. 6, Fig. 7A and Fig. S2 were plated on fibronectin-coated coverslips and left to attach and spread for 6 h before immunostainings. Human plasma fibronectin was from Millipore (FC010) and was used at 20 μg/mL in 100 mM bicarbonate buffer pH 8.5 for coating coverslips overnight at 4°C.

### Confocal fluorescence microscopy of cells and image processing

For live cell imaging, right before microscopy and due to the absence of CO_2_ control on our microscope setup, the culture medium was exchanged by Leibovitz medium (Gibco 21083027) supplemented with 10% fetal bovine serum and antibiotics. Cells were kept at 37°C in a heating chamber (OkoLab H301-T-UNIT-BL). Fluorescence images of live or fixed cells were acquired using a spinning disk unit (CSU-X1-M1 from Yokogawa) connected to the side-port of an inverted microscope (Eclipse Ti2-E from Nikon Instruments) using a Nikon Plan Apo ×100/1.45 NA oil immersion objective lens, 488-561- and 641-nm laser lines (Coherent) and an iXon Ultra 888 EMCCD camera (1024×1024 pixels, 13×13 μm pixel size, Andor, Oxford Instruments) resulting in an image pixel size of 65 nm. Z-stacks were acquired with a Δz interval of 0.4 μm. Exposure times were in the range of 0.5-3.0 s depending on the exact condition. For the non-diffuse vs diffuse cytosolic phenotype classification for septin mutant characterization, acquisition parameters were kept the same among imaging sessions. For actin or microtubule co-labeling in live cells, cells were incubated for 30 min with 0.5 μM of SiR-actin or 60 min with 0.5 μM of SiR-tubulin and 10 μM of verapamil in culture medium (SiR Cytoskeleton Kit, Spirochrome SC006).

Images were processed with the open-source image processing software ImageJ/Fiji. All shown images, except for the ones used for the septin-actin co-localization analysis that were acquired as single z-planes, are maximum intensity projections of two consecutive z-planes contrasted manually in order to optimize the image display. For septin-actin co-localization measurements, acquired channels of single z-planes, for septin and actin, were individually processed as follows: images were subjected to automatic contrast enhancement, allowing 0.1% of saturated pixels, then to a blurring with a Gaussian filter of radius 1.0 and a subsequent background subtraction using a rolling ball radius of 7 pixels. A manual intensity threshold was used when calculating Pearson and Manders co-localization coefficients, using the JACoP plugin for ImageJ (Bolte and Cordelières 2006).

Images shown in Fig. 1Biv, Fig. 7A and Fig. S2 were acquired on a Zeiss LSM 710 laser scanning confocal microscope using a PlanApochromat 100x/1.4 NA oil immersion objective lens, 488-543- and 633-nm laser lines for excitation, with all channels at 1AU for the pinholes. Z-stacks were acquired with a Δz interval of 0.48 μm. All shown images are single z-planes and were processed with ImageJ/Fiji.

### Super-resolution structured illumination microscopy

#### Sample preparation and image acquisition

Cells for super-resolution structured illumination (SIM) microscopy were plated on high precision (170 ± 5 µm thick) 18×18mm glass coverslips from Zeiss (474030-9000-000) and prepared for immunostainings as detailed in the section “Immunostaining after live-cell extraction vs after extraction post-fixation”; all images shown in Fig. 6 and Fig. 8G employ extraction post-fixation. For microtubule stainings used for microtubule width measurements in Fig. 6F,H, cells were fixed with −20°C-prechilled methanol for 2 min at −20°C and rinsed with PBS before overnight permeabilization/blocking and antibody incubations as described in the above section. Primary antibodies were mouse tubulin (1:1,000, Sigma T9026) and rabbit anti-SEPT7 (1:500, IBL 18991). Secondary antibodies were goat AlexaFluor488-conjugated anti-mouse IgGs (1:400, Thermo Fisher Scientific A11001) and goat AlexaFluor633-conjugated anti-rabbit IgGs (1:400, Thermo Fisher Scientific A21070). Images in Fig. 6 were acquired on a Zeiss Elyra PS.1 super-resolution microscope using an alpha PlanApochromat 100x/1.46 NA DIC M27 Elyra oil immersion objective lens, 488-, 561-, and 642-nm laser lines for excitation and respective BP495-550, BP570-620 and LP655 emission filters. Z-stacks were acquired with a Δz interval of 0.101 μm. Images were processed and channel-aligned with the Zeiss ZEN Black software. Images in Fig. 8G were acquired on a DeltaVision OMX SR (Leica Microsystems/Cytiva) super-resolution microscope using an Olympus PlanApo N 60x/1.42 NA oil immersion objective lens, 488- and 640-nm laser lines for excitation and respective 528/48 and 683/40 emission filters. Z-stacks were acquired with a Δz interval of 0.125 μm. Images were processed and channel-aligned with the DeltaVision softWoRx 7.0.0 software. All shown images are single z-planes and were prepared with ImageJ/Fiji.

#### Septin fiber diameter and length measurements

Full width at half maximum (FWHM) measurements for measuring the diameter of microtubules (MT) and septin fibers in SIM images (Fig. 6F,H) were made with a custom-generated Matlab code (FilamentAnalysis.mlx), the source code of which is available at gitHub.com/cchandre/Polarimetry. A line was drawn perpendicular to the axis of the MT or to the long axis of the septin fiber, and the FWHM was extracted from the intensity profile using the *findpeaks* Matlab function. We measured the width of MTs and septin fibers at multiple positions along their length and in multiple microtubules and multiple septin fibers for each SF type per cell. Box plots depicting the distribution of FWHM measurements (Fig. 6F) were prepared using GraphPad Prism (one data point corresponds to one width measurement). The central mark indicates the median, and the bottom and top edges of the box indicate the 25th and 75th percentiles, respectively. The whiskers extend to the minimum and maximum values. The number of measurements per condition (MT or SF type) is indicated in the respective legend. A Kruskal-Wallis test followed by a multiple comparison test was used for comparing the distributions.

Length measurements were implemented in the same custom-generated code. A line was drawn parallel to the long axis of the septin fiber and the length extracted with the *curveLength* Matlab function. We measured the length of multiple septin fibers for each SF type per cell. Scatter dot plots depicting the distribution of length measurements (Fig. 6J) were prepared using GraphPad Prism (one data point corresponds to one length measurement). The number of measurements per SF type is indicated in the respective legend. Bars depict median values.

#### Numerical simulations for fiber size estimation

Numerical simulations of the expected FWHM in SIM images (“image diameter” in Fig. 6G) as a function of the real fiber diameter (“fiber diameter” in Fig. 6G) were made with a custom-generated Matlab code (Convolution_1D.m), the source code of which is available at gitHub.com/cchandre/Polarimetry. A Gaussian point spread function (PSF) was used, and the curve was generated from the convolution of this PSF with an increasing fiber diameter size, using the conv Matlab function. Assuming a real antibody-decorated MT diameter size of ∼60 nm (Weber et al., 1978), the convolution curve permits to deduce the PSF size from the measured FWHM in isolated microtubule fibers (the median value is used). This PSF size being linearly dependent on the emission wavelength, it is then rescaled to account for the wavelength difference used in MT vs septin imaging: MTs and septins were imaged at 488 and 642 nm, respectively, for SIM in Fig. 6, whereas MTs and septins for SIM in Fig. 8G were imaged at 640 and 488 nm, respectively. To predict the real width of the respective septin fiber diameters (Fig. 6H), the convolution curve was finally used for the estimated PSF, using as input the measured septin FWHM.

#### Production and purification of recombinant human septin complexes

Wild-type nonfluorescent and SEPT2-msfGFP hexamers and octamers-9_i3, SEPT2NCmut-msfGFP hexamers and octamers-9_i3, and SEPT2-sfCherry2 octamers-9_i3 were produced and purified as described in (Iv et al., 2021). Briefly, plasmids expressing SEPT2, SEPT2-msfGFP or SEPT2NCmut-msfGFP, and plasmids co-expressing SEPT2, SEPT2-msfGFP or SEPT2NCmut-msfGFP and SEPT6, were co-transformed with plasmids co-expressing SEPT6 and SEPT7 (Addgene #174499), or SEPT7 and SEPT9_i3 (Addgene #174501), for generating recombinant nonfluorescent, SEPT2-msfGFP, or SEPT2NCmut-msfGFP hexamers and octamers-9_i3 (Iv et al., 2021). Plasmids co-expressing SEPT2-sfCherry2 and SEPT6 were co-transformed with plasmids co-expressing SEPT7 and SEPT9_i3 (Addgene #174501) to generate recombinant SEPT2-sfCherry2 octamers-9_i3. The N-terminus of SEPT2 is tagged with a His_6_-tag, and the C-terminus of SEPT7 (for isolation of hexamers), or the C-terminus of SEPT9 (for isolation of octamers), is tagged with a Strep-tag. A purification scheme comprising a Strep-Tactin affinity column to capture Strep-tagged complexes, followed by a nickel affinity column to retain the Strep-tagged complexes that also bear His_6_-tagged septins isolates hexamers and octamers (Iv et al., 2021).

Co-transformed *E. coli* BL21(DE3) were selected on LB agar plates with carbenicillin and spectinomycin each at 100µg/mL. A single colony was selected to prepare an overnight LB medium preculture at 37°C with antibiotics at 100 µg/mL. Terrific broth with antibiotics at 50 µg/mL, typically 3.5-5 L, was inoculated with the pre-culture and incubated at 37°C. Bacteria were left to grow to A_600nm_ ∼ 0.6-0.8 before inducing expression with 0.5 mM IPTG for overnight expression at 17°C. The culture was stopped by centrifuging at 3,400 g for 15 min and 4°C, and the supernatants were pooled and further centrifuged at 5,000 g for 10 min and 4°C. Bacteria pellets were stored at −20°C until protein purification. Bacteria expressing msfGFP- and sfCherry2-tagged septins yield yellow-greenish and pink-reddish pellets, respectively.

On the day of purification, the pellet was resuspended in ice-cold lysis buffer (50 mM Tris-HCl pH 8, 300 mM KCl, 5 mM MgCl_2_, 0.25 mg/mL lysozyme, 1 mM PMSF, cOmplete™ protease inhibitor cocktail (1 tablet per 50 mL), 10 mg/L DNase I, 20 mM MgSO_4_) and lysed on ice using a tip sonicator with 5 cycles of 30 s “ON”, 15 s “OFF”. The lysate was clarified by centrifugation for 30 min at 20,000 g and 4°C, and the supernatant loaded on a StrepTrap HP column. Strep-tag-II-containing septin complexes were eluted with 50 mM Tris-HCl pH 8, 300 mM KCl, 5 mM MgCl_2_, and 2.5 mM desthiobiotin. The pooled fractions were then loaded to a HisTrap HP column, and His_6_-tag-containing complexes eluted with 50 mM Tris-HCl at pH 8, 300 mM KCl, 5 mM MgCl_2_, and 250 mM imidazole. Only the highest-concentration peak fractions were collected. Both affinity steps were performed on an ÄKTA pure protein purification system at 4°C (Cytiva). To remove imidazole, we either performed overnight dialysis or used a PD-10 column, also including DTT in this last step. The final elution buffer, in which septins are stored, was 50 mM Tris-HCl pH 8, 300 mM KCl, 5 mM MgCl_2_, and 1 mM DTT. Protein concentration was assessed with absorbance measurements at 280 nm from the calculated extinction coefficients using ExPASy, and protein aliquots were flash-frozen in liquid nitrogen and stored at −80°C until further use.

Chemicals used for recombinant septin complex production and purification are as follows. *E. coli* BL21(DE3) from Agilent (200131). Carbenicillin (C3416), spectinomycin (S4014), LB broth medium (L3022), LB agar (L2897), SOC medium (S1797) from Sigma. Terrific Broth from MP Biomedicals (091012017). IPTG (EU0008-C) and lysozyme (5933) from Euromedex. Imidazole from Fisher Scientific (Fisher Chemical I/0010/53). PMSF (78830), cOmplete™ Protease Inhibitor Cocktail Tablets (Roche, 11836145001), DNase I (Roche, 10104159001), *d*-Desthiobiotin (D1411), and DTT (D0632) from Sigma. HisTrap HP 1 mL columns (17524701) and StrepTrap HP 1 mL columns from Cytiva (28907546). 20K MWCO Slide-A-Lyzer cassettes from Thermo Scientific (87735). PD-10 desalting columns from Cytiva (17085101).

#### Sample preparation for fluorescence microscopy of *in vitro* reconstituted actin and septins

To prepare flow cells, glass slides and coverslips were cleaned for 15 min in base-piranha solution (Milli-Q water, 30% ammonium hydroxide, 35% hydrogen peroxide at a 5:1:1 volume ratio), rinsed with Milli-Q water and stored in 0.1 M KOH up to one month. Right before assembling flow cells, slides and coverslips were rinsed with Milli-Q water and dried with synthetic air. Flow cells with ∼10 μL channels were assembled by sandwiching ∼2-mm-wide and ∼2.5-cm-long strips of Parafilm between a cleaned glass slide and coverslip and melting on a hot plate at 120°C. The resulting chambers were passivated by incubating for 45 min with 1 M KOH, rinsing with actin polymerization buffer (5 mM Tris-HCl pH 8, 50 mM KCl, 1 mM MgCl_2_, 0.2 mM Na_2_ATP, 1 mM DTT), incubating for another 45 min with 0.2 mg/mL PLL-PEG, and rinsing with actin polymerization buffer. Flow cells were placed in a Petri-dish along with tissue paper soaked in water to prevent flow channels from drying during the incubation steps and until use.

Lyophilized rabbit skeletal muscle G-actin was resuspended to 5 mg/mL (119 μM) in G-buffer (5 mM Tris-HCl pH 8, 0.2 mM Na_2_ATP, 0.1 mM CaCl_2_, 1 mM DTT), aliquots snap-frozen in liquid nitrogen and stored at −80°C. Frozen aliquots were thawed and centrifuged for 30 min at 120,000 g in a benchtop Beckman air-driven ultracentrifuge (Beckman Coulter Airfuge, 340401) to clear the solution from aggregates. Clarified G-actin was kept at 4°C and used within 3-4 weeks.

For actin-septin reconstitution experiments, thawed septin aliquots were cleared for 15 min at 120,000 g in a Beckman airfuge right before use. To polymerize G-actin in the presence of septins, we mixed G-actin, previously diluted with G-buffer to 5 μM, with septins, either nonfluorescent ones or msfGFP-labeled septins (at 20% msfGFP molar ratio for wild-type septins, and 100% GFP for SEPT2NC septins) to a final actin concentration of 1 μM and a final septin concentration of 0.3 μM, right before polymerization in actin polymerization buffer, additionally containing 1 mM Trolox, 2 mM protocatechuic acid (PCA), 0.1 μM protocatechuate 3,4-dioxygenase (PCD) and 0.1% w/v methylcellulose. To fluorescently label actin filaments, we polymerized G-actin in the presence of 1 μM Alexa Fluor 568-conjugated phalloidin.

Actin-septin samples were prepared with a final volume of 10 μL, were loaded immediately into passivated flow channels upon mixing of the components to start polymerization, and flow channels were sealed with VALAP (1:1:1 vasoline:lanoline:paraffin). The contributions of KCl and MgCl_2_ from the septin elution buffer were taken into account to yield the same final composition of actin polymerization buffer. Actin-septin samples were incubated overnight at room temperature (RT) in the dark before observation. To polymerize septins in the absence of actin, we followed the same procedure as above, but replaced the G-actin solution with G-buffer. Septins were used at 20% msfGFP and 20% sfCherry2 molar ratio for wild-type septins and at 100% GFP for SEPT2NC septins.

The sources and identifiers for proteins, materials and chemicals are as follows. Glass slides (26×76 mm) (AA00000102E01FST20) and glass coverslips (24×60 mm) (BB02400600A113FST0) from Thermo Scientific. Ammonium hydroxide solution (221228) and hydrogen peroxide solution (95299) from SIGMA. PLL-PEG from SuSoS AG (PLL(20)-g[3.5]-PEG(2)). Rabbit skeletal muscle G-actin from Cytoskeleton, Inc. (AKL99). Alexa Fluor 568-phalloidin from Thermo Scientific (A12380). Methylcellulose (M0512), Trolox (238813), protocatechuic acid (03930590), protocatechuate 3,4-dioxygenase (P8279) from Sigma.

#### Confocal fluorescence microscopy of reconstituted actin-septins and image processing

Reconstituted actin-septin assemblies were imaged on the same spinning disk microscope setup described for imaging cells using the same objective lens and camera. Images were acquired with an exposure time of 0.1 s. Actin-septin bundles were imaged close to the surface. Septin filament bundles were also found at the surface, but the clusters of interconnected filament bundles were observed floating in the bulk of the flow channels. To capture such clusters, z-stacks were acquired over 10-50 μm using a Δz interval of 0.5 μm. Images were processed with ImageJ/Fiji. Images of actin-septin bundles are from single planes. Images of septin filament bundles are from maximum-intensity z projections. The contrast of all images shown was adjusted post-acquisition so that both dim and bright structures are visible without saturation. All images use an inverted grayscale, with bright signals appearing black in a white background.

### Metal-induced energy transfer assays

U2OS cells were transfected with SEPT9_i3-mApple, SEPT9_i3-mApple-CAAX, or GAP43-mApple with FuGeneHD (Promega E2311). 16h post-transfection, cells were plated on glass coverslips (for obtaining reference lifetime measurements, see below) and on gold-coated glass coverslips, previously cleaned with 70% ethanol. Cells were left to attach and spread for 24h, then fixed for 15 min using 4% paraformaldehyde (Electron Microscopy Sciences 15714) in cytoskeleton buffer (10 mM MES – pH 6.1 with NaOH, 150 mM NaCl, 5 mM EGTA, 5 mM glucose, 5 mM MgCl_2_). The excess of cytosolic protein content was washed out with a permeabilization/blocking step (0,1% saponin,1% BSA in PBS) for 1h at room temperature. Labelling of F-actin in U2OS cells was achieved with AlexaFluor568-phalloidin (Invitrogen A12380) at 165 nM in permeabilization/blocking solution for 1h. The samples were maintained in PBS until and throughout the measurements.

Metal-induced energy transfer (MIET) was performed following the concept introduced by Enderlein and coworkers (Chizhik et al., 2014). Briefly, we measured the fluorescence lifetime of emitters in the vicinity of a 18 nm-thick gold film. From the calibration of the fluorescence lifetime dependence with the distance to the gold film (Chizhik et al., 2014), the distance between the fluorophore and the metal is recovered. The MIET calibration curve was computed using the MIET-GUI Matlab code developed by the Enderlein group (https://projects.gwdg.de/projects/miet/repository/raw/MIET_GUI.zip?rev=ZIP). For mApple, we used a peak emission wavelength at 610 nm and a quantum yield of 49%. For Alexa Fluor 568, the emission peak was 603 nm and the quantum yield 69%. Our calculation used the fluorescence lifetime of the dyes measured on a glass coverslip, in the absence of the metal layer, to account for the slight 0.1 ns lifetime change induced by the functionalization of the dye to septin, GAP43 or phalloidin. An isotropic orientation of the fluorophores is assumed (Chizhik et al., 2014).

We used a gold film of 18 nm thickness deposited by electron-beam assisted evaporation of gold on a borosilicate glass coverslip (Bühler Syrus Pro 710). A 2 nm-thick chromium layer is used to promote the adhesion of gold on the glass coverslip. For the MIET calibration, the refractive indexes of the gold and chromium layers were taken from (Rosenblatt et al., 2020) and (Johnson and Christy, 1974), respectively while the refractive index of 1.52 for the borosilicate glass coverslip was provided by the supplier (D 263 M glass by Schott AG).

The fluorescence lifetime measurements were performed with a home built confocal microscope with a 557 nm iChrome-TVIS laser (Toptica GmbH, pulse duration 3 ps, 40 MHz repetition rate) and a Zeiss C-Apochromat 63x, 1.2 NA water immersion objective. The excitation power remained below 2 μW on the sample to avoid photobleaching during the measurement. The fluorescence light was collected by the same microscope objective and filtered using a dichroic mirror (ZT 405/488/561/640rpc, Chroma), long-pass filter (ET570LP, Chroma) and bandpass filter (ET595/50m, Chroma). The confocal pinhole diameter was 50 μm. The photon counting detection used an avalanche photodiode (MPD-5CTC, Picoquant) connected to a time correlated counting module (HydraHarp400, PicoQuant). The temporal resolution (full width at half maximum of the instrument response function) was measured to be 38 ps. The fluorescence lifetime histograms were fitted using SymPhoTime 64 software (PicoQuant GmbH) with a reconvolution taking into account the measured instrument response function. All the histograms were fitted using a biexponential function which provided a better fit to the intensity decay than a single exponential decay. About 20% of the total detected intensity corresponded to the short lifetime component (below 0.5 ns) which was not considered further for the analysis. The MIET distance measurements were taken on the long lifetime component which represented more than 80% of the total detected photons. The distribution of calculated distances from lifetime measurements for each condition is represented in box plots using GraphPad Prism (one data point per cell for each condition). The central mark indicates the median, and the bottom and top edges of the box indicate the 25th and 75th percentiles, respectively. The whiskers extend to the minimum and maximum values. The number of cells per condition is indicated in the respective legend. One-way ANOVA followed by a multiple comparison test was used for comparing the distributions.

### Supported lipid bilayer assays

#### Small unilamellar vesicle formation

We used three types of lipids, 1,2-dioleoyl-sn-glycero-3-phospho-(1’-myo-inositol-4’,5’-bisphosphate) (ammonium salt) (PI(4,5)P_2_) (Sigma 850155P), 1,2-dioleoyl-sn-glycero-3-phosphocholine (DOPC) (Sigma 850375C), and 1,2-dioleoyl-sn-glycero-3-phosphoethanolamine-N-(Cyanine 5) (DOPE-Cy5) (Sigma 810335C), all from Avanti Polar Lipids. The lipids were mixed in chloroform, or, in case PI(4,5)P_2_ was present, in a 20:9:1 chloroform:methanol:water mixture in a glass vial. The organic solvent was then evaporated completely using a stream of N_2_ followed by overnight incubation in a dessicator. The dried lipid film was resuspended in buffer to give a total lipid concentration of 0.25 mM. We used a sodium citrate buffer of pH 4.8 (50 mM citrate, made of equal molarity trisodium citrate and citric acid mixed in a 2:3 volume ratio, 50 mM KCl, 0.1 mM ethylenediaminetetraacetic acid) in case PI(4,5)P_2_ was present, and otherwise F-buffer of pH 7.4 (20 mM Tris-HCl, 2 mM MgCl_2_, 50 mM KCl, 1 mM DTT). The lipids were dissolved by four cycles of 1 min vortexing and 5 min incubation. Finally, small unilamellar vesicles (SUVs) were obtained by sonicating the lipid solution using an Ultrasonic homogeniser series HD 2000.2 sonicator equipped with a BR30 cup resonator (Bandelin) at 10% amplitude for 30 minutes with pulses of 5s on and 5s off to avoid excessive heating.

#### Protein preparation

Unlabelled septin octamers and SEPT2-msfGFP octamers were purified in house as previously reported (Iv et al., 2021). The protein was stored in aliquots in septin storage buffer (20 mM Tris HCl pH 7.4, 2 mM MgCl_2_, 300 mM KCl, 1 mM DTT), at −80°C. Before each experiment, unlabelled and labelled septin octamers were mixed in a 9:1 molar ratio in septin storage buffer at a total concentration of 1800 nM. Lyophilized monomeric actin (G-actin) from rabbit skeletal muscle (Hypermol 8101-03) was resuspended following the manufacturer’s instructions and dialyzed against G-buffer (5 mM Tris-HCl pH 7.8, 0.1 mM CaCl_2_, 0.2 mM ATP, and 1 mM DTT) to remove residual disaccharides from the freeze-drying process. Protein aggregates were removed by centrifugation at 148,000 x g for 1h and the supernatant was snap-frozen and stored in aliquots at −80◦C. Fluorescently tagged G-actin was prepared by covalent modification with Alexa Fluor™ 594 Carboxylic Acid (Thermo Fisher Scientific 15461054) (Alvarado and Koenderink, 2015). Before experiments, G-actin aliquots were thawed, and any aggregates were removed by leaving the protein on ice for at least 2h and subsequently centrifuging at 148,000 × g for 20 min. Unlabelled and fluorescent actin were mixed in a 9:1 molar ratio in G-buffer at a total G-actin concentration of 5 μM.

#### Sample preparation

Supported lipid bilayers (SLB) were formed in custom-made flow channels made of nr. 1 Menzel coverslips (Thermo Fisher Scientific 11961988) and glass slides (Thermo Fisher Scientific 11879022). The coverslips and glass slides were first cleaned in base piranha solution (5% hydrogen peroxide, 5% ammonium hydroxide) at 70°C for 10 minutes, extensively washed with Milli-Q water, and stored in Milli-Q water for a maximum of 5 days. Just before use, a coverslip and a slide were dried with a stream of N_2_ gas. Flow channels were prepared by sandwiching 2×20 mm parafilm strips separated by ∼3 mm between the glass slide and the coverslip. The parafilm was then melted by placing the chambers on a hot plate at 120°C and gently pressing on top with clean tweezers. After cooling down, an SUV solution (7-12 μL, depending on the distance between the parafilm strips) was pipetted into the channels and incubated in a humid chamber for at least 20 minutes to promote SUV rupture and SLB formation. Residual SUVs were removed by washing with 4 channel volumes of F-buffer for DOPC SLBs or with 2 channel volumes of sodium citrate buffer followed by 2 channel volumes of F-buffer for 5% PI(4,5)P_2_ SLBs. DOPC SLBs contained 99.7% DOPC and 0.3% DOPE-Cy5; 5%PIP2 SLBs also contained 94.7% DOPC and 0.3% DOPE-Cy5.

Septin octamers and actin were co-polymerized at room temperature in polymerization buffer (20 mM Tris-HCl pH 7.4, 2 mM MgCl_2_, 50 mM KCl, 1 mM DTT, 0.5 mM ATP, 1 mM GTP) supplemented with 1 mM Trolox to suppress blinking, and an oxygen scavenging system composed of 1 mM protocatechuic acid and 0.05 μM of procatechuate 3,4-dioxygenase to minimize photobleaching. We first prepared a 5x master buffer (100 mM Tris HCl pH 7.4, 10 mM MgCl_2_, 5 mM DTT, 2.5 mM ATP, 5 mM GTP, 5 mM Trolox and 5 mM protocatechuic acid). To prepare the sample, we mixed the master buffer (5-fold dilution), 0.05 μM of procatechuate 3,4-dioxygenase, the G-actin mix (5-fold dilution to give a final concentration of 1 μM), and the septin mix (6-fold dilution, to give a final concentration of 300 nM), in that order. The mixture was either immediately added to the flow channels containing the SLBs and incubated for 1h in a humid environment, or first incubated in the tube for 1h to promote septin-actin bundle formation and then added to the flow channels using a cut pipette tip to minimize bundle disruption. The channels were then sealed with Dow Corning® high-vacuum silicone grease (Sigma Z273554).

#### Image acquisition

The samples were immediately imaged using a Nikon Ti2-E microscope complemented with a Gataca iLAS2 azimuthal TIRF illumination system. The sample was illuminated with 488-nm and 561-nm lasers (Gataca laser combiner iLAS2) to visualize the septin and the actin signals, respectively. The fluorescence signal was split with a Cairn Research Optosplit II ByPass containing a Chroma ZT 543 rdc dichroic mirror and filtered with either a 525/50 or a 600/50 chroma bandpass filter. The images were recorded with an Andor iXon Ultra 897 EM-CCD camera using an exposure time of 50 ms. To check that the SLBs were uniform and free of defects, we examined DOPE-Cy5 distribution by illuminating with a 642-nm laser filtered with a 708/75 chroma bandpass filter and recorded using an exposure time of 20 ms. We checked SLB fluidity by fluorescence recovery after photobleaching of DOPE-Cy5.

### Atomic force microscopy

#### Sample preparation

12-mm glass coverslips were coated with 0.1 mg/mL PLL-PEG (PLL(20)-g[3.5]-PEG(2), Susos) before being illuminated with a deep-UV lamp through a quartz-chrome photomask bearing the micropattern features (Front Range Photomask) designed using AutoCAD (Autodesk). We used Y-shaped micropatterns with a spread area of ∼1500 mm^2^. Micropatterned coverslips were then incubated with 25 µg/mL fibronectin and 5 µg/mL fibrinogen-GFP, the latter for visualizing micropatterns. Wild type U2OS cells treated with siLacZ siRNA (6mer+8mer), with SEPT9 siRNA (6mer), and U2OS cells treated with SEPT7 siRNA and also transfected with msfGFP-SEPT7Gmut2 (8mer) were seeded on fibronectin-coated micropatterns 48h post-electroporation. Cells were incubated for 5-7 h to attach and spread adopting a triangular shape. The expression of msfGFP-SEPT7Gmut2 for the 8mer condition was confirmed through the detection of fluorescence in each measured cell.

#### Force Spectroscopy experiments and data analysis

Atomic force microscopy-force spectroscopy (AFM-FS) was performed on the dorsal perinuclear region of individual cells at room temperature. We used a MLCT-Bio-DC (D) cantilever featuring a 4-sided regular pyramid with a semi-open angle of 35°. The spring constant of the cantilevers was determined in air using the Sader method (Sader et al., 2012) and the optical lever sensitivity from the thermal spectrum in liquid (Sumbul et al., 2020). Force-distance curves were acquired applying a maximum force of 0.8 nN with a ramp range of 5 µm, at the same approach and retract velocity of 5 µm/s on a Nanowizard 4 AFM microscope (JPK-Bruker). The indentation depth was on the order of 1 μm. 31 cells, 29 cells and 23 cells were probed for the 6mer+8mer, 6mer and 8mer condition, respectively. For each cell, about 15-30 force curves were acquired across 3 different contact points, resulting in a total of 576, 630 and 501 force curves for the 6mer+8mer, 6mer and 8mer condition, respectively. To extract the cell viscoelastic properties, we fitted the Ting numerical viscoelastic model for a 4-sided regular pyramidal tip of semi-open angle (theta) to the experimental force-distance curves (Bilodeau, 1992; Efremov et al., 2017):

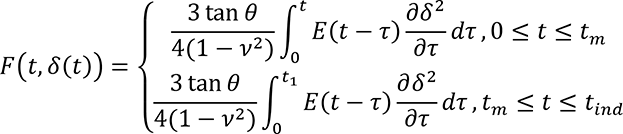

where F is the applied force; δ is the indentation; t is the time since initial contact, t_m_ is the duration of approach trace, t_ind_ is the duration of complete indentation cycle, and t_1_ determined by solving the equation

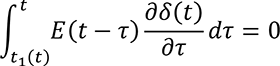

We assumed that the time-dependent Young’s modulus followed a power law relationship:

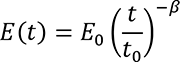

where *E*_0_ is the elastic modulus at time *t*_0_, *β* is the fluidity of the cell and *t*_0_ is the reference time, arbitrarily assumed 1s. A viscous drag force (*F*_d_) proportional to the trace velocity (*v*) was also added to the force traces using a precalibrated value of the viscous drag coefficient (*b*= 5 pN·s/µm), *F*_d_=*b*·*v*.

The values of log10(*E*_0_) and β extracted from each force measurement were pooled by cell and then averaged. The data was reproduced in 3 independent experiments and their distribution represented in box plots using GraphPad Prism (one data point per cell for each condition). The central mark indicates the median, and the bottom and top edges of the box indicate the 25th and 75th percentiles, respectively. The whiskers extend to the minimum and maximum values. The number of cells per condition is indicated in the respective legend. *E*_0_ values (in Pa) were plotted on a log scale. One-way ANOVA followed by a multiple comparison test was used for comparing the distributions of E_0_ values using the log10(*E*_0_) values, given the log-normal distribution of *E*_0_.

### Single particle tracking Photo-Activated Localization Microscopy (sptPALM)

#### Cell culture

Mouse embryonic fibroblasts were cultured in DMEM (Gibco 10313-021) with 10% fetal calf serum (FCS, Eurobio scientific CVFSVF00-01). Transient transfections of plasmids were performed 2 days before experiments using the Amaxa nucleofector (Lonza VPD-1004). The cells were detached with trypsin/EDTA, the trypsin was inactivated using DMEM with 10% FCS, and the cells were washed and suspended in serum-free Ringer solution (150 mM NaCl, 5 mM KCl, 2 mM CaCl_2_, 2 mM MgCl_2_, 10 mM HEPES-Na pH 7.4, 2 g/L glucose), then incubated for 30 min in Ringer solution before plating on fibronectin-coated glass coverslips (human plasma fibronectin at 10 µg/ml, Roche 10838039001).

#### Plasmids

SEPT9_i3-mEos3.2 and SEPT9_i3-mEos3.2-CAAX were cloned in a pCMV plasmid backbone with seamless cloning into a NheI/BamHI linearized vector (primers in Table S1). EGFP-human β-actin was provided by A. Matus (Friedrich Miescher Institute for Biomedical Research, Switzerland). The mEos2-actin construct was generated from EGFP-actin as described in (Rossier et al., 2012). GFP-human paxillin (isoform alpha) was used as described in (Rossier et al., 2012).

#### Optical setup and image acquisition

sptPALM acquisitions were steered by MetaMorph software (Molecular Devices) with an inverted motorized microscope (Nikon Ti) equipped with a temperature control system (The Cube, The Box, Life Imaging Services), a Nikon CFI Apo TIRF 100x oil, NA 1.49 objective and a Perfect Focus System, allowing long acquisition in TIRF illumination mode.

Imaging was performed at least 3 hours after seeding the cells on fibronectin-coated coverslips mounted in a Ludin chamber (Life Imaging Services). For photoactivation localization microscopy, cells expressing mEos2 and mEos3.2 tagged constructs were photoactivated using a 405 nm laser (Omicron) and the resulting photoconverted single molecule fluorescence was excited with a 561 nm laser (Cobolt Jive™). Both lasers illuminated the sample simultaneously. Their respective power was adjusted to keep the number of the stochastically activated molecules constant and well separated during the acquisition. Fluorescence was collected by the combination of a dichroic and emission filters (D101-R561 and F39-617 respectively, Chroma) and a sensitive EMCCD (electron-multiplying charge-coupled device, Evolve, Photometric). The acquisition was performed in streaming mode at 50 Hz. Either GFP-paxillin or GFP-actin were imaged using a conventional GFP filter cube (ET470/40, T495LPXR, ET525/50, Chroma). Using this filter cube does not allow spectral separation of the unconverted pool of mEos from the GFP fluorescent signal. However, with all of the constructs used, whether the mEos signal was highly or poorly enriched in FAs, we were still able to detect FAs.

#### Single molecule segmentation and tracking

A typical sptPALM experiment leads to a set of at least 4000 images per cell, analyzed in order to extract molecule localization and dynamics. Single molecule fluorescent spots were localized and tracked over time using a combination of wavelet segmentation and simulated annealing algorithms (Izeddin et al., 2012; Racine et al., 2006; Racine et al., 2007). Under the experimental conditions described above, the resolution of the system was quantified to 59 nm (Full Width at Half Maximum, FWHM). This spatial resolution depends on the image signal to noise ratio and the segmentation algorithm (Cheezum et al., 2001) and was determined using fixed mEos2 samples. We analyzed 130 2D distributions of single molecule positions belonging to long trajectories (>50 frames) by bi-dimensional Gaussian fitting, the resolution being determined as 2.3 *s_xy_*, where *s_xy_* is the pointing accuracy.

For the trajectory analysis, FAs ROIs were identified manually from GFP-paxillin or GFP-actin images. The corresponding binary mask was used to sort single-molecule data analyses to specific regions. We analyzed trajectories lasting at least 260 ms (≥13 points) with a custom Matlab routine analyzing the mean squared displacement (MSD), which describes the diffusion properties of a molecule, computed as (Eq. 1):

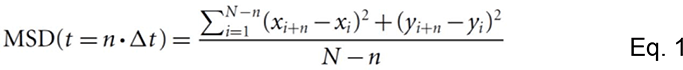

where *x_i_* and *y_i_* are the coordinates of the label position at time *I x Δt*. We defined the measured diffusion coefficient *D* as the slope of the affine regression line fitted to the *n*=1 to 4 values of the MSD(*n x Δt)*. The MSD was computed then fitted on a duration equal to 80% (minimum of 10 points, 200 ms) of the whole stretch by (Eq. 2):

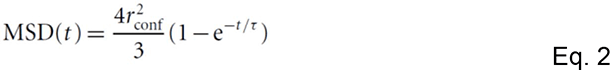

where *r*_conf_ is the measured confinement radius and *τ* the time constant *τ* = (*r*_conf_² / 3*D*_conf_). To reduce the inaccuracy of the MSD fit due to downsampling for larger time intervals, we used a weighted fit. Trajectories were sorted in 3 groups: immobile, confined diffusion and free diffusion. Immobile trajectories were defined as trajectories with *D*<0.011 μm^2^.s^-1^, corresponding to molecules which explored an area inferior to the one defined by the image spatial resolution ∼(0.05μm)² during the time used to fit the initial slope of the MSD (Rossier et al., 2012) (4 points, 80 ms): *D*_threshold_=(0.059 μm)²/(4×4×0.02s)∼0.011 μm^2^.s^-1^. To separate trajectories displaying free diffusion from confined diffusion, we used the time constant calculated *τ* for each trajectory. Confined and free diffusion events were defined as trajectories with a time constant respectively inferior and superior to half the time interval used to compute the MSD (100 ms). Statistical significance tests were prepared using GraphPad Prism.

### Modeling of human septin complexes

Models of full-length human septin complexes were built for analyzing and interpreting split-GFP experiments. The septin GTP-binding domains (GBDs) used as templates for the SEPT2, 6 and 7 models using SWISS-MODEL homology modeling software (Waterhouse et al., 2018) were from PDB 7M6J (Leonardo et al., 2021), the most complete human septin hexamer structure to date, which includes *α*0 helices for SEPT6 and 7. As solved in its integrity, the SEPT6 GBD remained unchanged and was used as is. The SEPT7 GBD structure was completed using SWISS-MODEL. As the use of the SEPT2 GBD from 7M6J for modeling SEPT2 led to clashes in the modeled SEPT2-SEPT2 NC interface, the SEPT2 GBD subunit was modeled using the SEPT7 GBD structure from 7M6J as a template. The lack of structural information for the short N-terminal extensions of SEPT2, 6, and 7 prompted us to model them as disordered segments using Phyre2 (Kelley et al., 2015). The C-terminal domains of SEPT2, 6 and 7 were modeled with CCFold (Guzenko and Strelkov, 2018) for the coiled-coil (CC) parts and Phyre2 for the flexible parts, as detailed in (Iv et al., 2021). The homodimeric parallel SEPT2CC was used unaltered with respect to (Iv et al., 2021). The previously modeled SEPT6 and 7 helices in the SEPT6-SEPT7 parallel coiled-coil in (Iv et al., 2021) were repositioned slightly after comparison with the only parallel septin CC structure to date (PDB 6WCU) (Leonardo et al., 2021). GBDs, N- and C-terminal extensions were then combined with PyMOL open-source software. When necessary, the disordered segments were manually modified to avoid steric clashes and to adjust distances. The SEPT9_i3 model used was the one built for (Iv et al., 2021) and included already N- and C-terminal extensions. Hexameric SEPT2-SEPT6-SEPT7-SEPT7-SEPT6-SEPT2 and octameric SEPT2-SEPT6-SEPT7-SEPT9-SEPT9-SEPT7-SEPT6-SEPT2 complexes were built by fitting the modeled structures to the hexamer from the PDB 7M6J.

To analyze and interpret the split-GFP experiments, the entire constructs used in the assays, including β10- and β11-tagged septins and the reconstituted GFP, were modeled. To this aim, the split GFP structure (PDB 4KF5) was added to the modeled septin complexes. The flexible linkers linking the reconstituted GFP to the septin of interest were built manually using PyMOL; their straight-ish appearance in the models is due to the polypeptide chains being built as linear structures. To mimic paired septin filaments with narrow spacing (Leonardo et al., 2021), mediated by homodimeric SEPT2 antiparallel CCs (Fig. 4E), septin complexes were duplicated and placed parallel to each other with a gap of ∼5 nm. The bent conformation of the septin was built by rotating the CC domain manually by 90 degrees relative to the GBD. The helices within the homodimeric SEPT2 antiparallel CC were positioned using the antiparallel SEPT4CC structure from PDB 6WB3 as a reference (Leonardo et al., 2021). All manual interventions were realized using PyMOL.

#### Statistics and reproducibility

The distributions of measurements, or of phenotypes in the case of septin mutant characterization, are represented with GraphPad Prism using box plots, violin plots and scatter dot plots as indicated in the respective methods sections and legends. Bars, error bars (SD or SEM) and box plot features are as indicated in the respective figure legends. The number of measurements in each plot and the numbers of experiments are indicated in the respective figure legend or methods. Statistical significance tests were performed with with GraphPad Prism. The tests applied and the obtained P values are mentioned in the respective figure legend. Experiments were repeated at least three times independently to ensure reproducibility. Experiments from Fig. 7C-F; Fig 7H; Fig. S4 were performed twice. Experiments from Fig. 7G,I were performed once. No data were excluded from the analyses.

#### Data availability

All data supporting the findings of this study are available within the article and its supporting information files. The source datasets generated and analyzed during the current study are available from the corresponding author on reasonable request.

#### Code availability

The source codes for the custom-generated Matlab codes for measurements of fiber diameter (FWHM) and length, and for numerical simulations of expected fiber diameter (FWHM) from SIM images has been deposited to Github. The respective links are mentioned in the relevant methods sections.

## Acknowledgements

We thank Josette Perrier and Cendrine Nicoletti (iSm2, Marseille, France) for generously hosting protein production and purification experiments. The authors further thank Artemis Kosta, Hugo Le Guenno and the Microscopy Core Facility of the Institut de Microbiologie de la Méditerranée (IMM) for SIM microscopy with the DeltaVision OMX SR microscope. We further thank R. Sterling for technical assistance and the IINS Cell culture facility, especially E. Verdier and N. Retailleau for technical help (IINS Cell Biology Facility, grant no. ANR-10-LABX-43). We would also like to thank J. B. Sibarita (IINS) for his support with sptPALM analysis. This research received funding from the Agence Nationale de la Recherche (ANR grants ANR-17-CE13-0014 SEPTIMORF to M.M.; ANR-17-CE09-0026-01 AntennaFRET to J.W.; ANR-20-CE42-0003 3DPolariSR to V.M. and O.R.), the Fondation ARC pour la recherche sur le cancer (grant ARCDOC42020010001242 to C.S.M.), and from the Cancéropôle PACA, Institut National du Cancer and Conseil Régional PACA (Bourse mobilité to M.M.). We further acknowledge financial support from the French Ministry of Research, CNRS and the Conseil Régional Nouvelle-Aquitaine (grant MechanoStem to O.R. and V.M.). We also acknowledge the France-BioImaging infrastructure supported by the French National Research Agency (ANR-10-INBS-04). This project has received funding from the European Research Council (ERC) under the European Union’s Horizon 2020 research and innovation programme (grant agreements No 723241 to J.W. and No 772257 to F.R.). This project has received funding from the European Union’s Horizon 2020 research and Innovation programme under the H2020-MSCA-ITN-2018 Grant Agreement n. 812772. This work was supported in part by the National Institutes of Health (R01GM122375 to S.K.). Confocal microscopy with the Zeiss LSM710 microscope and SIM with the Zeiss Elyra PS.1 microscope was performed at the UC Berkeley Biological Imaging Facility, which was supported in part by the National Institutes of Health S10 program under award numbers 1S10RR026866-01 and 1S10OD018136-01. G.C.L. and G.H.K. gratefully acknowledge financial support by the Netherlands Organization for Scientific Research (NWO/OCW) through the ‘BaSyC—Building a Synthetic Cell’ Gravitation grant (024.003.019).

The authors declare no competing interests.

---

**Figure S1.**
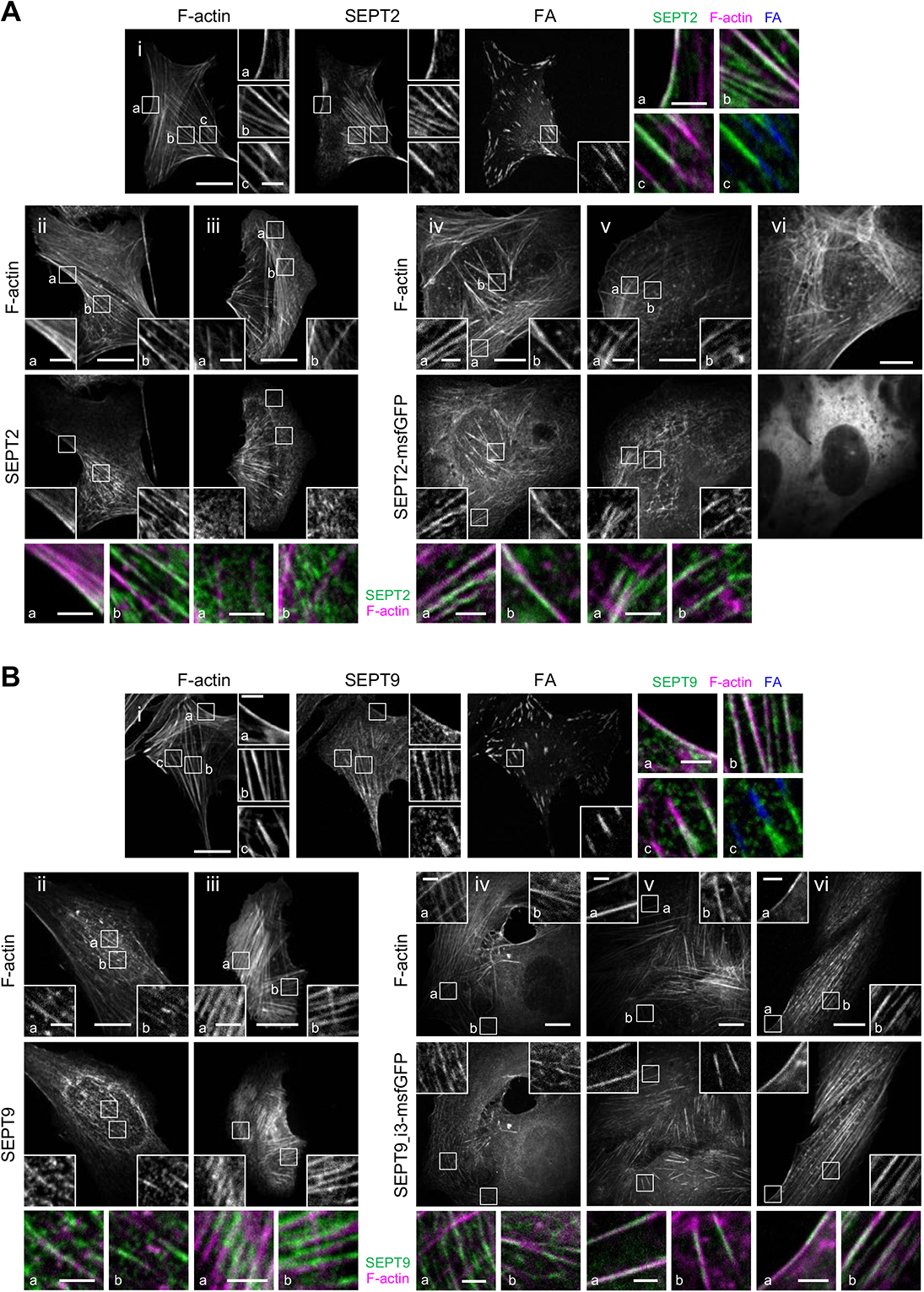
SEPT2 and SEPT9 distribution on different types of stress fibers in U2OS cells. **(A)** Representative confocal micrographs of SEPT2 immunostained cells (i-iii) and cells expressing SEPT2-msfGFP (iv-vi). SEPT2 immunostained cells are co-stained for F-actin (phalloidin) and the FA protein paxillin. Examples show SEPT2 localizing (i) to peripheral (a) and ventral (b,c) SFs and excluded from focal adhesions (FA) (c), (ii) to peripheral (a) and perinuclear actin caps (b), (iii) to transverse arcs (b) and excluded from dorsal SFs (a,b), (iv) to ventral SFs (a,b), (v) to transverse arcs (a,b) and excluded from dorsal SFs (a), and (vi) showing a diffuse cytosolic phenotype. **(B)** Representative confocal micrographs of SEPT9 immunostained cells (i-iii) and cells expressing SEPT9_i3-msfGFP (iv-vi). SEPT9 immunostained cells are co-stained for F-actin (phalloidin) and the FA protein paxillin. Examples show SEPT9 localizing (i) to peripheral (a) and ventral (b,c) SFs and excluded from focal adhesions (FA) (c), (ii) to perinuclear actin caps (a,b), (iii) to transverse arcs (a) and ventral SFs (b), (iv) to transverse arcs (a) and excluded from dorsal SFs (a) and to ventral SFs (b), (v) to ventral SFs (a,b), and (vi) to peripheral (a) and perinuclear actin caps (b). Scale bars in large fields of views, 10 μm. Scale bars in insets, 2 μm. Related to Fig. 1B.

**Figure S2.**
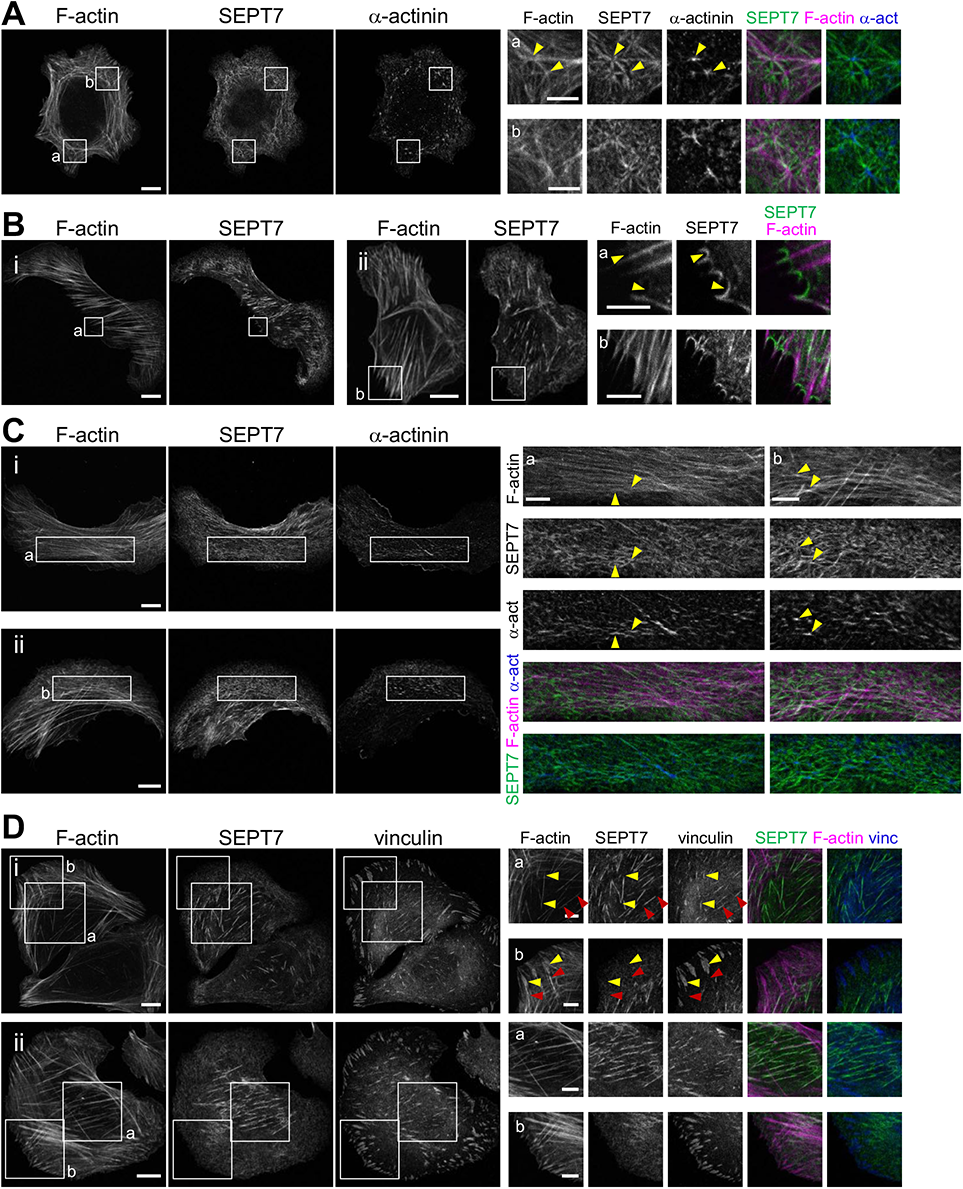
SEPT7 distribution on different types of stress fibers in U2OS cells. **(A-D)** Representative confocal micrographs of SEPT7 immunostained cells co-stained for F-actin (phalloidin) (A-D), and additionally stained for a-actinin (A,C) or the FA protein vinculin (D). Examples show SEPT7 localization to ventral actin nodes (A), to arcs and arc nodes (C), and to ventral SFs, excluded from FAs (D). Yellow arrowheads in (A) point to two *α*-actinin-enriched actin nodes. Yellow arrowheads in (B) point to curved segments of the membrane that appear devoid of phalloidin staining, but positive for SEPT7. Yellow arrowheads in (C) point to *α*-actinin-enriched arc nodes. Yellow and red arrowheads in (Di, box a) point to the septin-decorated regions of two ventral SFs, with SEPT7 excluded from the respective FAs. Yellow and red arrowheads in (Di, box b) point to two dorsal SFs, with SEPT7 excluded from them and from the respective FAs. Scale bars in large fields of views, 10 μm. Scale bars in insets, 5 μm. Related to Fig. 1B.

**Figure S3.**
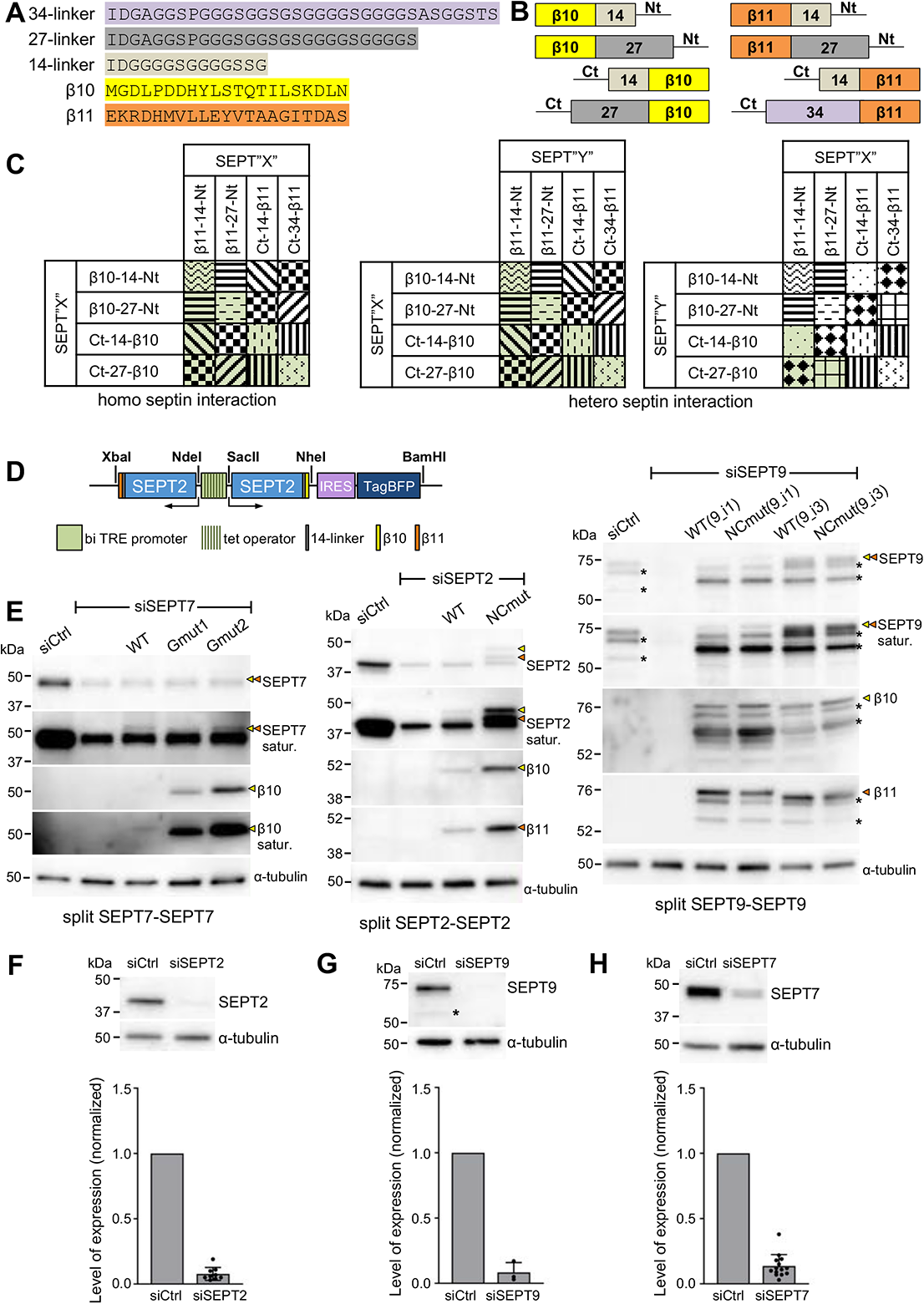
Design of the tripartite split-GFP complementation assay for probing septin organization. **(A)** Sequences of the β-10 and β11-tags used for all split assays and of the linker sequences tested in screening experiments (B,C); 14-residue linkers were used throughout this study. **(B)** Schematic of N- and C-terminal β-10 and β11-tag septin fusions tested in screening experiments (C) using short or long linkers (A). **(C)** Schematic of β-10 and β11-septin fusion combinations for screening tripartite split GFP complementation. Combinations with the same pattern were considered to be equivalent (for example, SEPT2-14-β10 / β11-14-SEPT2 and SEPT2-14-β11 / β10-14-SEPT2). The combinations in green are the ones tested experimentally. **(D)** Schematic of the pTRIP TRE Bi vector bearing a bidirectional tetracycline response element (TRE) promoter for the doxycycline-inducible co-expression of β10- and β11-tagged septins. An IRES-TagBFP cassette was used for monitoring septin expression. Restriction sites used for subcloning are indicated (see methods for details). **(E)** Left, Western blots of U2OS-Tet-On-GFP1-9 cell line lysates probed with anti-SEPT7, anti-β10 and anti-*α*-tubulin antibodies upon treatment with siRNAs targeting LacZ (siCtrl), SEPT7 (siSEPT7), and targeting SEPT7 while co-expressing wild-type β10- and β11-SEPT7 (WT), β10- and β11-SEPT7Gmut1 (Gmut1), and β10- and β11-SEPT7Gmut2 (Gmut2). Yellow and orange arrowheads point to bands correspond to β10- and β11-fusions. The SEPT7 and β10 blots are also shown saturated in purpose for displaying weaker bands. Molecular weight markers are shown on the left. Middle, Western blots of U2OS-Tet-On-GFP1-9 cell line lysates probed with anti-SEPT2, anti-β10, anti-β11 and anti-*α*-tubulin antibodies upon treatment with siRNAs targeting LacZ (siCtrl), SEPT2 (siSEPT2), and targeting SEPT2 while co-expressing wild-type SEPT2-β10 and −β11 (WT) or SEPT2NCmut-β10 and −β11 (NCmut). Yellow and orange arrowheads point to bands correspond to β10- and β11-fusions. The SEPT2 blot is also shown saturated in purpose for displaying weaker bands. Right, Western blots of U2OS-Tet-On-GFP1-9 cell line lysates probed with anti-SEPT9, anti-β10, anti-β11 and anti-*α*-tubulin antibodies upon treatment with siRNAs targeting LacZ (siCtrl), SEPT9 (siSEPT9), and targeting SEPT9 while co-expressing wild-type SEPT9-β10 and −β11 (WT) or SEPT9NCmut-β10 and -β11 (NCmut) for both SEPT9_i1 and SEPT9_i3. Yellow and orange arrowheads point to bands correspond to β10- and β11-fusions. The SEPT9 blot is also shown saturated in purpose for displaying weaker bands. Asterisks point to SEPT9 degradation products. **(F)** Western blot of U2OS cell lysates probed with anti-SEPT2 and anti-*α*-tubulin antibodies upon treatment with siRNAs targeting LacZ (siCtrl) or SEPT2 (siSEPT2). Molecular weight markers are shown on the left. Bottom, respective quantification of SEPT2 protein levels (mean+SD). Mean values (normalized to 1 for siCtrl) are from 3 independent siCtrl and 9 independent siSEPT2 treatments. SEPT2 was knocked down on average by 92%. **(G)** Same as (F) for SEPT9. The asterisk points to a SEPT9 degradation product. Mean values (normalized to 1 for siCtrl) are from 3 independent siCtrl and 3 independent siSEPT9 treatments. SEPT9 was knocked down on average by 92%. **(H)** Same as (F) for SEPT7. Mean values (normalized to 1 for siCtrl) are from 3 independent siCtrl and 12 independent siSEPT7 treatments. SEPT7 was knocked down on average by 86%.

**Figure S4.**
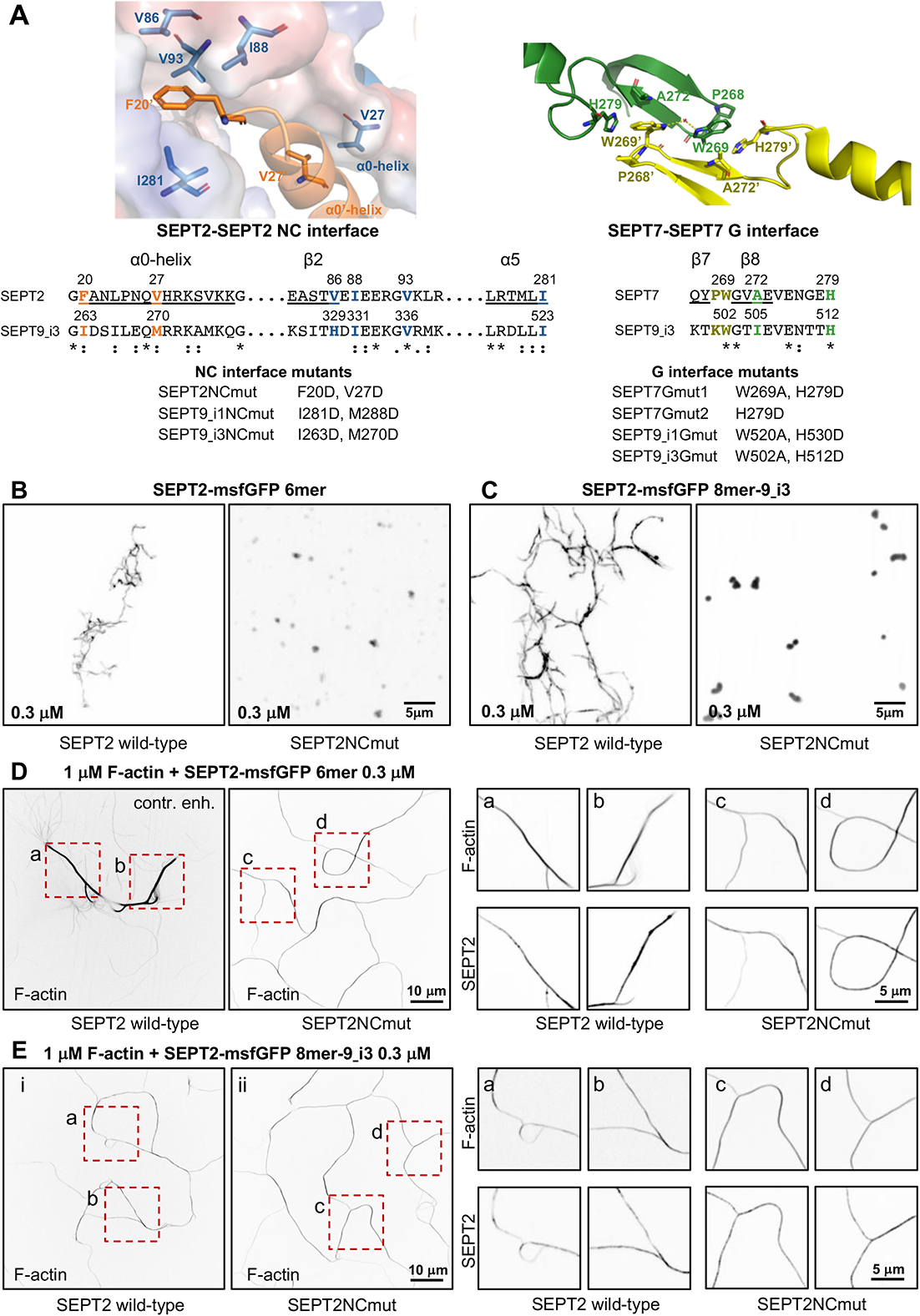
Septin interface mutants used in this study and cell-free reconstitution of septin assembly. **(A)** Left, Top, conserved residues in the SEPT2-SEPT2 NC interface are shown in the crystal structure of human SEPT2 homodimers (PDB 2QA5) (Sirajuddin et al., 2007). The backbone structure is displayed as a cartoon representation in PyMOL, with critical residues represented as sticks (deep blue and red for nitrogen and oxygen atoms, respectively). Residues F20 from the hook-loop of one SEPT2 subunit (orange) interact with the hydrophobic cleft formed by V86, I88, V93, and I281 of the adjacent SEPT2 subunit (blue). The importance of this phenylalanine in anchoring the α0 helix at the NC interface was emphasized only recently (Cavini et al., 2021). The blue subunit’s surface representation highlights the complementary of shape between the two SEPT2 in this interface. The interaction between the α0 helices of each subunit is also stabilized via a hydrophobic interaction between their respective V27. Left, Middle, sequence alignment of the regions including the residues shown in the NC interface structure for SEPT2 and SEPT9_i3. The structural elements (α0, β2, α5) related to these residues are underlined and shown above the sequences. The consensus symbols are from ClustalW alignments of all human septins (*, fully conserved residue; colon, conservation between residues with strongly similar physicochemical properties; period, conservation between residues with weakly similar physicochemical properties). We note that the residues described above are strictly or physicochemically conserved (except for V86), highlighting their importance in stabilizing the SEPT2-SEPT2 NC interface. Left, Bottom, NC interface mutants used in this study. A mutation of F20D/I263D is expected to destabilize the hydrophobic pocket depicted above, whereas a V27D/M270D is expected to destabilize the α0 helices interface. Importantly, a strictly conserved aspartate (SEPT2 E90, corresponding to SEPT6 E90 and SEPT7 E102 which are well defined in the cryo-EM structure of the SEPT6-SEPT7 NC interface (Mendonca et al., 2021)) in the loop connecting 12 and 13 is pointing to the hydrophobic cleft where the phenylalanine resides. The F20D mutation is thus expected to result in a repulsion between the aspartate and glutamate and contribute further to the destabilization of the NC interface. Right, Top, conserved residues in the SEPT7-SEPT7 G interface are shown in the crystal structure of human SEPT7 homodimers (PDB 6N0B) (Brognara et al., 2019). The backbone structure is displayed as a cartoon representation in PyMOL, with critical residues represented as sticks (deep blue and red for nitrogen and oxygen atoms, respectively). Residues W269 of one SEPT7 subunit (yellow) interact with residues W269, A272 and H279 in the adjacent SEPT7 subunit (green) (Sirajuddin et al., 2007; Zent et al., 2011). W269 from adjacent subunits interact via a water molecule bridge through hydrogen bonds. In addition, each W269 is engaged in π-π interactions with H279 and CH-π interactions with A272 of the opposite subunit. Right, Middle, sequence alignment of the regions including the residues shown in the G interface structure for SEPT7 and SEPT9_i3. The structural elements (17, 18) related to these residues are underlined and shown above the sequences. The consensus symbols are from ClustalW alignments of all human septins (*, fully conserved residue; colon, conservation between residues with strongly similar physicochemical properties). Notice that W269 and H279 are both strictly conserved, showing their importance in stabilizing this interface. Right, Bottom, G interface mutants were used in this study. The presence of both mutations W269A and H279D in SEPT7 and SEPT9 is expected to destabilize the SEPT7-SEPT7 and SEPT7-SEPT9 G-interfaces. The loss of the aromatic cycle properties in the mutant W269A does not allow the abovementioned critical interactions mediated by the wild-type Trp. W269A is expected to destabilize H279 and potentially change its orientation. In addition, the much smaller size of the alanine will poorly mimic the hydrophobic interaction between W269 and H279, weakening the G-interface. Note that W269 is in the vicinity of Tyr267 of the same subunit. This tyrosine interacts with the nucleotide buried within the G-interface. Consequently, any mutations destabilizing W269 could dramatically destabilize the overall G-interface because of a domino effect. Similarly, H279D is expected to preclude hydrophobic interactions with W269 and thus destabilize the latter. The single mutation H279D in SEPT7 is expected to destabilize the SEPT7-SEPT7 G-interface when present in both SEPT7 subunits, but not the SEPT7-SEPT9 interface with wild-type SEPT9. **(B-C)** Representative spinning disk fluorescence images of septin filament assembly upon polymerization of hexamers (B) and octamers-9_i3 (C) at the indicated final protomer concentration. Protomers contained either wild-type SEPT2 (left panels) or SEPT2NCmut (right panels). Images use an inverted grayscale. Related to Fig. 2F. **(D-E)** Representative spinning disk fluorescence images of reconstituted actin filaments, polymerizing in the presence of septin hexamers (D) and septin octamers (E). Protomers contained either wild-type SEPT2 or SEPT2NCmut. Actin filaments are visualized with AlexaFluor568-conjugated phalloidin, and septins with SEPT2-msfGFP. One example of large fields of view are shown for each condition, depicting cross-linking of actin filaments; only actin labeling is shown. The image for actin in the presence of wild-type hexamers is contrast-enhanced in purpose in order to saturate the actin bundles so that weaker-intensity single actin filaments are also visible. Insets on the right side of each panel show higher magnifications of selected regions of interest on the left (dashed squares in red). Two regions of interest (a,b for wild-type SEPT2 and c,d for SEPT2NCmut) are shown in each case, depicting both the actin (top row) and septin (bottom row) signals. Scale bars in all large fields of views, 10 μm. Scale bars in all insets, 5 μm.

**Figure S5.**
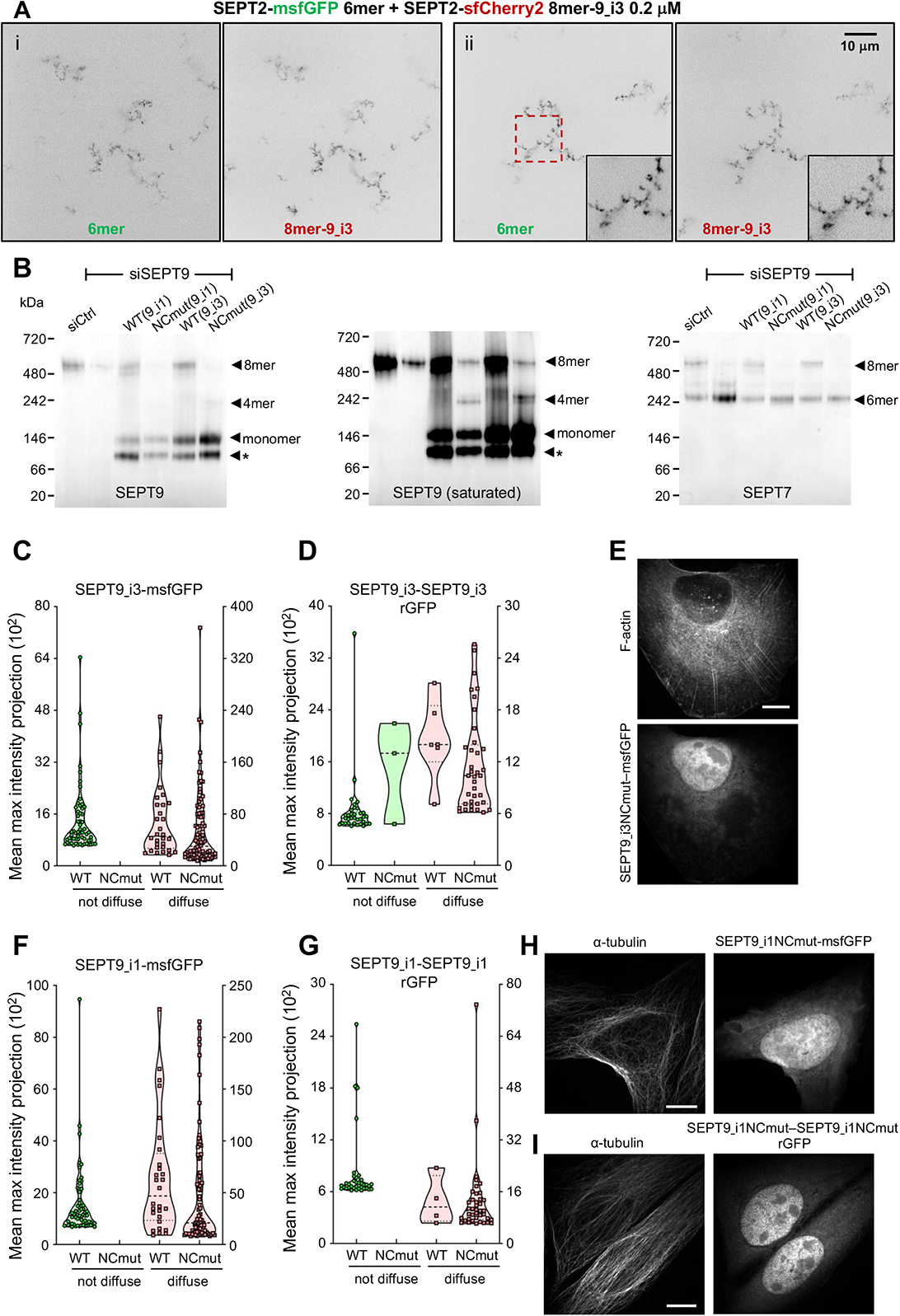
SEPT9 NC mutants used for split SEPT9-SEPT9 assays. **(A)** Representative spinning disk fluorescence images of septin filament assembly upon co-polymerization of hexamers containing SEPT2-msfGFP and octamers-9_i3 containing SEPT2-sfCherry2 at the indicated final protomer concentration. Images use an inverted grayscale. Scale bars in all large fields of views, 10 μm. **(B)** Western blot following native PAGE of U2OS cell lysates probed with anti-SEPT9 (left and middle) and anti-SEPT7 (right) antibodies upon treatment with siRNAs targeting LacZ (siCtrl), SEPT9 (siSEPT9), and targeting SEPT9 while expressing wild-type SEPT9-msfGFP (WT) or SEPT9NCmut-msfGFP (NCmut) for both SEPT9_i1 and SEPT9_i3. The SEPT9 blot is also shown saturated in purpose for displaying weaker bands. Arrowheads point to the sizes of the indicated complexes. The asterisks point to SEPT9 degradation. Molecular weight markers are shown on the left. **(C)** Violin plots depicting the distribution of diffuse cytosolic (red datapoints) vs. non-diffuse (green datapoints) phenotypes in cells expressing wild-type SEPT9_i3-msfGFP or SEPT9_i3NCmut-msfGFP. Data points are from a total of 90 cells each for wild-type and mutant SEPT9 distributed among the two phenotypes. **(D)** Violin plots depicting the distribution of diffuse cytosolic (red datapoints) vs. non-diffuse (green datapoints) phenotypes from reconstituted GFP (rGFP) in GFP1-9 cells co-expressing wild-type SEPT9_i3-110 and −111 or SEPT9_i3NCmut-110 and −111. Data points are from a total of 40 cells each for wild-type and mutant SEPT9 distributed among the two phenotypes. **(E)** Representative example of a cell expressing SEPT9_i3NCmut-msfGFP showing a diffuse cytosolic phenotype. Scale bar, 10 μm. **(F-H)** Same as (C-E) for SEPT9_i1. **(I)** Representative example of a GFP1-9 cell co-expressing SEPT9_i1NCmut-110 and −111 showing a diffuse cytosolic phenotype. Scale bar, 10 μm. Related to Fig. 3C,E.

**Figure S6.**
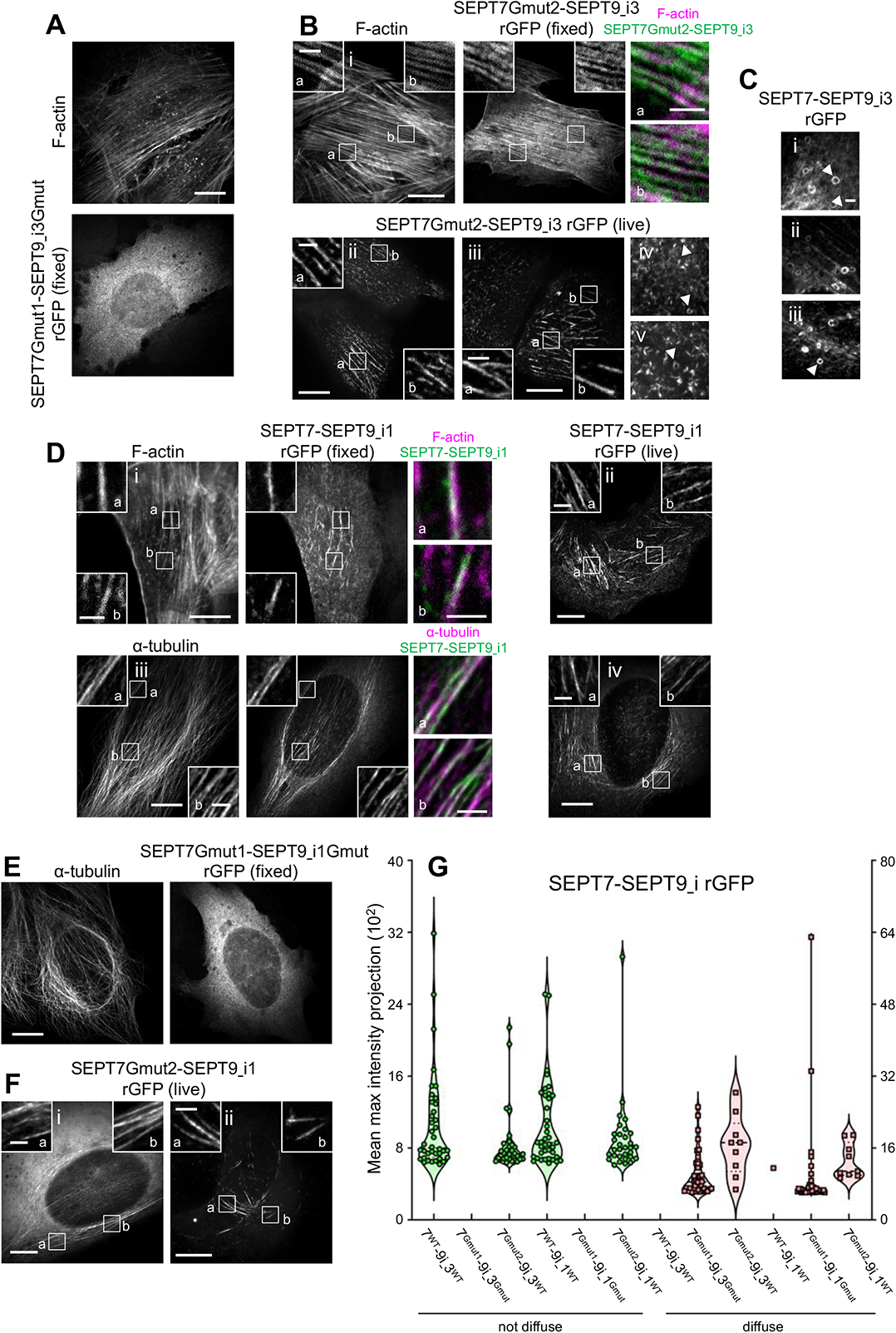
SEPT7 and SEPT9 G mutants used for split SEPT7-SEPT9 assays. **(A)** Representative example of a GFP1-9 cell co-expressing β11-SEPT7Gmut1 and SEPT9_i3Gmut-β10 showing a diffuse cytosolic phenotype. The cell is co-stained for F-actin (phalloidin). Scale bar, 10 μm. **(B)** Representative examples of GFP1-9 cells (i-v) co-expressing β11-SEPT7Gmut2 and SEPT9_i3-β10. The fixed cell is co-stained for F-actin (phalloidin). Examples show reconstituted GFP (rGFP) localizing (i) to ventral SFs (a,b), (ii) to perinuclear actin caps (a,b), (iii) to ventral SFs (a,b), and (iv-v) rings (arrowheads). Scale bars in large fields of views, 10 μm. Scale bars in insets, 2 μm. **(C)** Examples of rings (arrowheads) in GFP1-9 cells co-expressing β11-SEPT7 and SEPT9_i3-β10. Scale bar, 2 μm. **(D)** Representative examples of GFP1-9 cells co-expressing β11-SEPT7 and SEPT9_i1-β10 (i-iv). The fixed cells are co-stained for F-actin (phalloidin) (i) or for *α*-tubulin (iii). Examples show rGFP localizing to ventral SFs (i,ii) or to microtubules (iii,iv). Scale bars in large fields of views, 10 μm. Scale bars in insets, 2μm. **(E)** Representative example of GFP1-9 cell co-expressing β11-SEPT7Gmut1 and SEPT9_i1Gmut-β10 co-stained for *α*-tubulin showing a diffuse cytosolic phenotype. Scale bar, 10 μm. **(F)** Representative examples of GFP1-9 cells co-expressing β11-SEPT7Gmut2 and SEPT9_i1-β10 with localizing to microtubules. Scale bars in large fields of views, 10 μm. Scale bars in insets, 2 μm. **(G)** Violin plots depicting the distribution of diffuse cytosolic (red datapoints) vs. non-diffuse (green datapoints) phenotypes in GFP1-9 cells co-expressing the indicated combinations of β11-SEPT7 and SEPT9-β10 fusions. Data points are from a total of 40 cells each for each combination, distributed among the two phenotypes. Related to Fig. 3D.

**Figure S7.**
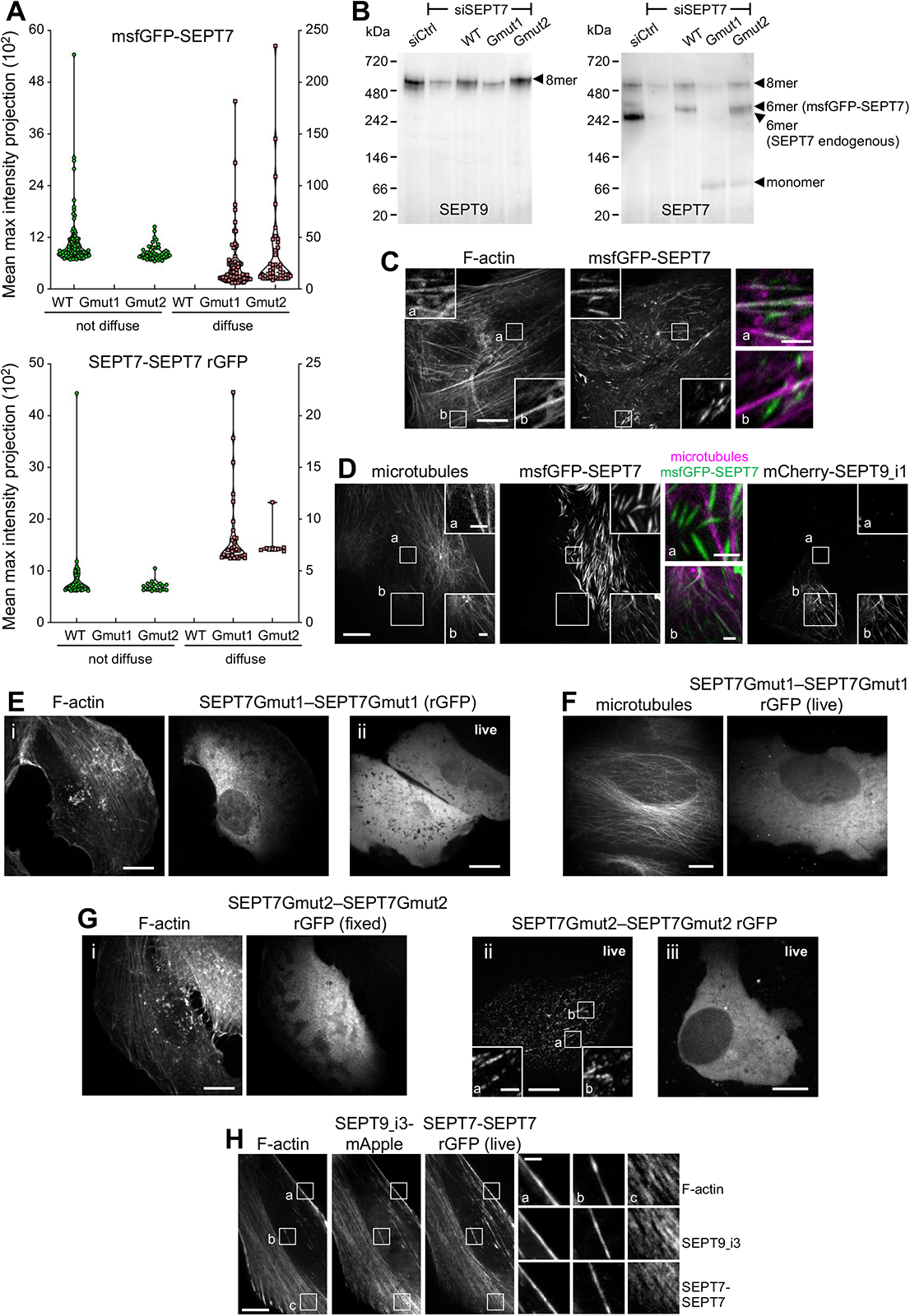
SEPT7 G mutants used for split SEPT7-SEPT7 assays. **(A)** Top, Violin plots depicting the distribution of diffuse cytosolic (red datapoints) vs. non-diffuse (green datapoints) phenotypes in cells expressing wild-type msfGFP-SEPT7 or msfGFP-SEPT7NCmut. Data points are from a total of 71 cells for wild-type, 68 cells for SEPT7Gmut1 and 90 cells for SEPT7Gmut2 distributed among the two phenotypes. Bottom, Violin plots depicting the distribution of diffuse cytosolic (red datapoints) vs. non-diffuse (green datapoints) phenotypes from reconstituted GFP (rGFP) in GFP1-9 cells co-expressing wild-type β10- and β11-SEPT7, β10- and β11-SEPT7Gmut1, or β10- and β11-SEPT7Gmut2. Data points are from a total of 40 cells for wild-type, 33 cells for β10- and β11-SEPT7Gmut1 and 29 cells for β10- and β11-SEPT7Gmut2 distributed among the two phenotypes. **(B)** Western blot following native PAGE of U2OS cell lysates probed with anti-SEPT9 (left) and anti-SEPT7 (right) antibodies upon treatment with siRNAs targeting LacZ (siCtrl), SEPT7 (siSEPT7), and targeting SEPT7 while expressing wild-type msfGFP-SEPT7 (WT), msfGFP-SEPT7Gmut1 (Gmut1), or msfGFP-SEPT7Gmut2 (Gmut2). Arrowheads point to the sizes of the indicated complexes. Molecular weight markers are shown on the left. **(C)** Representative confocal micrograph of a cell expressing msfGFP-SEPT7 co-stained for F-actin (phalloidin) localizing to ventral SFs (a) and to ectopic bundles devoid of phalloidin staining (b). Scale bars in large fields of views, 10 μm. Scale bars in insets, 2 μm. Related to Fig. 4A. **(D)** Representative example of a cell (bottom left) co-expressing msfGFP-SEPT7 and mCherry1-SEPT9_i1 and labeled for microtubules (SiR-tubulin) showing msfGFP-SEPT7 localizing to microtubules (b). A cell expressing only msfGFP-SEPT7 (top right) shows msfGFP-SEPT7 localizing to ectopic bundles not co-localizing with microtubules (a). Scale bars in large fields of views, 10 μm. Scale bars in insets, 2 μm. Related to Fig. 4G. **(E)** Representative examples of GFP1-9 cells (i,ii) co-expressing β10- and β11-SEPT7Gmut1 showing a diffuse cytosolic phenotype. The fixed cell is co-stained for F-actin (phalloidin). Scale bar, 10 μm. **(F)** Representative example of GFP1-9 cells co-expressing β10- and β11-SEPT7Gmut1, mCherry-SEPT9_i1 (not shown) and labeled for microtubules (SiR-tubulin) showing a diffuse cytosolic phenotype. Scale bar, 10 μm. **(G)** Representative examples of GFP1-9 cells (i-iii) co-expressing β10- and β11-SEPT7Gmut2. The fixed cell is co-stained for F-actin (phalloidin). Examples show diffuse cytosolic phenotypes (i,iii) of the rGFP and rGFP localizing to SFs (ii). Scale bars in large fields of views, 10 μm. Scale bars in insets, 2 μm.**(H)** Representative example of a GFP1-9 cell co-expressing β10- and β11-SEPT7, SEPT9_i3-mApple and labeled for F-actin (SiR-actin). Example shows rGFP localization to peripheral (a), ventral SFs (b) and transverse arcs (c). Scale bars in large fields of views, 10 μm. Scale bars in insets, 2 μm. Related to Fig. 4F.

**Figure S8.**
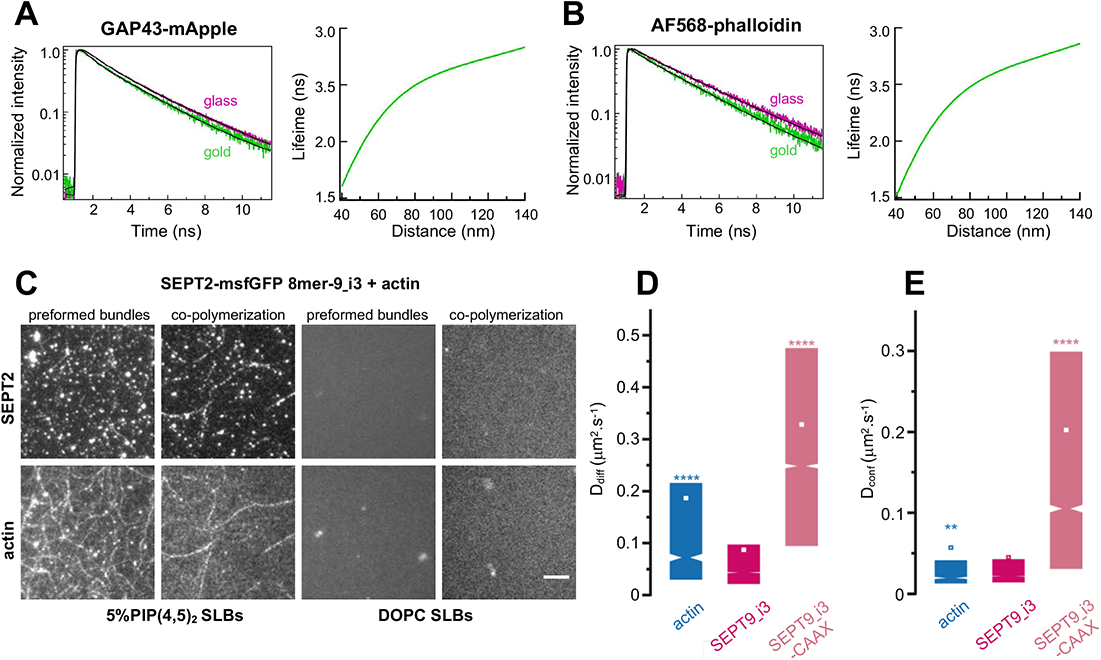
Septin filaments are closely apposed to the plasma membrane, are immobilized on actin stress fibers, and can mediate actin-membrane anchoring. **(A-B)** depict representative examples of lifetime decay traces for GAP43-mApple (A, left) and AF568-phalloidin (F-actin) (B, left) on glass and in the presence of gold. The solid lines represent the numerical fits, showing the lifetime reduction due to the MIET process. The respective calculated lifetime-distance dependences, used to calculate the distances of the fluorophores from the coverslip (Fig. 7E) are shown in the respective right panels. Related to Fig. 7B-E. **(C)** TIRF images of SEPT2-msfGFP 8mer-9_i3 and F-actin, either co-polymerized on top of a supported lipid bilayer (SLB), or co-polymerized in solution to form preformed bundles that were then flushed onto the supported lipid bilayer. The supported lipid bilayer was composed either of 5% of PI(4,5)P_2_, a septin-interacting lipid, and 95% DOPC (top panels), or 100% DOPC (bottom panels). Scale bar, 5μm. Related to Fig. 7F. **(D-E)** Box plots displaying the median (notch) and mean (square) ± percentile (25–75%) of diffusion coefficients corresponding to the free diffusion (D_diff_) (D) and confined diffusion (D_conf_) (E) trajectories outside FAs from sptPALM. Related to Fig. 7G-K. Statistical significance was obtained using two-tailed, non-parametric Mann–Whitney rank sum test. The different conditions were compared to the SEPT9_i3-mEos3.2 condition. The resulting P values are indicated as follows: ** P<0.01; **** P < 0.0001.

